# Tissue imprinting defines functional mosaic of dermal macrophages

**DOI:** 10.1101/2025.01.09.631670

**Authors:** Florens Lohrmann, Roman Sankowski, Stephan Schwer, Jana Neuber, Evi P Vlachou, Frederike Westermann, Emma Troesch, Julia Kolter, Anne Kathrin Lösslein, Kunal Das Mahapatra, Clarissa-Laura Döring, Nisreen Ghanem, Eva Kohnert, Aileen Heselich, Ori Staszewski, Gianni Monaco, Antonia Müller, Kilian Eyerich, Robert Zeiser, Marco Prinz, Burkhard Becher, Judith Zaugg, Clemens Kreutz, Sagar, Philipp Henneke

## Abstract

Dermal macrophages (macs) protect the skin from invading pathogens. They are derived from embryonic as well as hematopoietic progenitors. However, the functional impact of their diverse origin and the control networks defining different subsets remain unclear. Here, using multidimensional analysis of dermal macs, we reveal that the absence of circulating monocytes in interferon regulatory factor 8 (*Irf8*) deficient mice delays mac renewal during the steady state. Yet, the functional mosaic of dermal macs remains largely intact, i.e., major dermal mac subsets develop independently of monocyte replenishment. Thus, the tissue microenvironment is sufficient to induce alternative differentiation pathways and functional specialization of resident cells. Mycobacterial skin infection induces a steep increase in mac density due to monocyte-derived macs which execute urgent antibacterial functions and differentiate into site-adapted mac subsets in wildtype but not *Irf8^-/-^*mice, while long-term resident macs are required to initiate a tissue repair program already in early stages of infection.

In summary, we introduce a model, where an intricate network of specialized mac subsets develops to meet microanatomical needs and external cellular input is required only during immunological emergency situations.

**Highlights:**

- *Irf8^-/-^*-driven monocytopenia has negligible impact on homeostatic dermal macrophage diversity.
- Resident dermal macrophages have diverse specializations but remain flexible to adapt to challenges such as lacking monocyte influx
- Bone marrow-derived macrophages differentiate into specialized resident cells, with microenvironmental cues overriding origin-dependent programming.
- In chronic bacterial infections, distinct specialized bone-marrow-derived macrophages mount the defense, while resident macrophages activate a tissue-modifying program from early on.

## Introduction

The skin forms a microcosm of heterocellular interactions between immune cells and diverse structures such as hair bulbs, nerves, vessels and glands. It constitutes a barrier which integrates environmental cues, like mechanical irritation and those originating from microorganisms into its development and maintenance. Skin immunity never comes to rest but oscillates between states of activation and containment in a spatially diverse and dynamic fashion. Macrophages (macs) are the most abundant immune cells in the dermis as the major immune cell reservoir of the skin. Macs perform the complex task of maintaining the equilibrium at this site of interaction between host and environment^1,2^. Accordingly, dermal macs represent a heterogenous population with respect to origin, function and microanatomical localization in homeostasis and under exogenous stress^2^. Overall, tissue macs are derived from prenatally seeded yolk sac- and fetal liver-derived progenitors, and are exchanged by circulating Ly6C^hi^ (classical) monocytes (Mc) after birth in variable quantities^3,4^. In barrier tissues, such as the dermis and intestine, this process has been shown to occur more rapidly^1,5^ and to be accelerated during inflammatory processes^6^. On the other hand, we previously uncovered a small long-lived self-renewing dermal mac subset which surveils sensory nerves, highlighting the potential of macs to precisely adapt to distinct microanatomical niches with highest fidelity^2^. Other mac subsets associated with blood vessels and hair bulbs have also been reported^2,7,8^. However, spatiotemporal adaptation of macs in homeostasis and inflammation is not well resolved. Imprinting by the microenvironment has been described as a major process defining cellular properties, but the exact signals leading to the functional specialization of mac precursors remain largely unknown^9–12^. In situations of high demand, such as intradermal infections, the mac pool is expanded by incoming Mc, with quantitative involution occurring after resolution of infection^13–15^. Yet, exact knowledge on the highly dynamic differentiation process of immigrating mac progenitors into specialized resident macs is incomplete, despite important discoveries, e.g. that resident macs predominantly depend on M-CSF as a survival factor, whereas GM-CSF licenses macs for an inflammatory response^2,15–19^.

Intradermal bacterial infections including vaccination with live *Mycobacterium bovis* BCG (Bacillus-Calmette-Guérin), represent a highly instructive model to enable insights into the adaptation of innate dermal immunity to an enduring microbial trigger^20^. In healthy infants, mycobacteria infect and temporarily persist in macs but are contained at site. Inborn deficiency of *Irf8*, a master transcription factor in the development of mononuclear phagocytes in the bone marrow, leads to a profound paucity of Mc and to susceptibility for BCG dissemination after vaccination^21,22^ and probably environmental non-tuberculous mycobacteria, which cause chronic skin and soft tissue infections in children^23,24^. Mycobacterial soft tissue infections, including those related to BCG, are characterized by a heterocellular granuloma reaction, where macs are the dominant cell type^25^. In contrast, in purulent bacterial infections, neutrophils are the cellular key players^13^. In *Irf8^-/-^* mice, which – in reminiscence of the human phenotype - almost completely lack circulating Ly6C^hi^ Mc^26^, we previously found overall dermal mac numbers, as well as the highly differentiated nerve-associated macs (sNaM), unaltered, indicating compensatory maintenance mechanisms involved in establishing the resident mac pool^2^.

Here, we combined experimental dermal BCG infection with transgenic *Irf8* models to gain multidimensional insights into the functional organization of mac diversity and specific dermal immunity, particular regarding limits of resident mac plasticity and adaptability in situations of highest need. In particular, we employed tissue microscopy, spatial transcriptomics, single-cell and bulk transcriptomics, mathematical modelling along with fate mapping and bone marrow transplantation to dissect the effects of severe stress on tissue mac homeostasis such as the lack of circulating Mc (*Irf8^-/-^*) and emergency recruitment of macs during intradermal BCG infection. We find that resident macs adapt to cover the full range of transcriptional properties in homeostasis, while newly recruited Mc-derived macs are uniquely required to fight mycobacterial skin infection.

## Material and methods

### Mice

All mice were on C57BL/6 genetic background. C57BL/6J mice were purchased from Jackson Laboratories (USA) or Charles River Laboratories (Germany) and bred in the CEMT animal facility, Freiburg, Germany. CD45.1-mice (B6.SJL-*Ptprc^a^ Pepc^b^*/BoyJ), *Ccr2^-/-^*-mice (B6.129S4-Ccr2tm1Ifc/J), β*-actin-GFP*-mice (C57BL/6-Tg(CAG-EGFP)131Osb/LeySopJ), *Csf2rb^-/-^* mice (Zurich, B6.129S1-*Csf2rb^tm1Cgb^*/J), and *Irf8^-/-^* mice (B6(Cg)-Irf8tm1.2Hm/J) were purchased from Jackson Laboratories and bred in the animal facility. *Mrc1*-*CreER x Rosa26tdtomato* mice (Mrc1-cre/ERT2 x B6.Cg-Gt(ROSA)26Sortm9(CAG-tdTomato)Hze/J) have been previously described^27^. Mice were housed under specific pathogen-free conditions and kept in 12h light/dark cycles and food and water were provided *ad libitum*. If not mentioned specifically, adult mice were analyzed at 8 and 16-weeks of age. In general, both sexes were used. For transplantation experiments same-sex pairs were used; no differences were observed within the parameters analyzed. All animal experiments were approved by the Federal Ministry for Nature, Environment and Consumer’s protection of the state of Baden-Wuerttemberg (proposal numbers: G16/64, G18/83, G18/115, G19/171).

### *In vivo* experiments

#### Bone marrow transplantation

For bone marrow transplantations, recipient mice were anaesthetized with Ketamin and Xylazin and lethally irradiated with a total dose of 9 Gy. They were either fully irradiated, or the head or one ear was shielded with a customized lead device. Subsequently, mice received 1×10^7^ donor-derived bone marrow cells intravenously (i.v).

#### Intradermal infection

Infections were induced by strictly intradermal injection of approximately 10^6^ CFU of the indicated pathogen into the external side of the ear under isoflurane narcosis. Mycobacteria were resuspended in PBS containing 5% mineral oil after rigorous sonification as outlined before^20^.

#### Adoptive monocyte transfer

Adoptive Mc transfer was performed directly after intradermal BCG infection. Peripheral blood mononuclear cells from two pooled β-actin-GFP-mice were generated via Ficoll gradient (Ficoll paque Plus, Cytiva, Sigma-Aldrich) and labelled with Biotin-conjugated anti-CD115 antibodies (Invitrogen). MACS® Microbeads (Miltenyi) were added according to manufacturer’s instructions and cells were passed twice through MACS® LS columns (Miltenyi). After centrifugation cells were resuspended in 100 µl PBS and injected i.v. into *Irf8^-/-^* recipient mice.

#### Fate mapping

*Mrc1^CreERT2^:R26-tdtomato* mice were induced by fractioned subcutaneous (s.c.) Tamoxifen injections under inhalative isoflurane narcosis. 4 mg Tamoxifen, resuspended in 200 µl corn oil, was injected, distributed over 4 injection sites. The injection was repeated after 48 hours.

#### BrdU

BrdU was diluted into the drinking water with the start of intradermal infection with a concentration of 0.8 mg/ml for the indicated time periods and protected from light.

### Tissue preparation

For dermal macs, mouse ears (dermis and epidermis) were cut into small pieces and subjected to enzymatic digestion by dispase (0.25 U/ml; STEMCELL Technologies), collagenase II (1 mg/ml; Worthington), and DNase I (0.04 mg/ml; Roche) in HBSS with 10% FCS for 2 hours (h) at 1400 rpm in a shaker at 37°C. Cells were passed through a 70 µm tissue strainer. FcγII/III receptors were blocked by prior incubation with anti-CD16/32 antibody (eBioscience). All cells were resuspended in FACS buffer (PBS, 2% FBS and 2mM EDTA).

For determination of bacterial growth and tissue cytokine concentrations, the skin was cut into small pieces and subjected to tissue lysis (Quiagen^TM^ TissueLyser with 50/s oscillations for 10 min with 3 mm tungsten carbide beads). Samples for ELISA were stored at -80°C until further analysis.

Whole blood was drawn either from the Vena facialis or, for terminal analysis, from the retrobulbar plexus. It was subjected twice to eBioscience^TM^ RBC lysis buffer according to the manufacturer.

Bone marrow was flushed from the long bones with a 26G needle in PBS through a 70 µm tissue strainer and subjected to further analysis or culture.

For Mc MAC sorting, PBMCs were generated with Ficoll-Paque^TM^ Plus (Cytiva), stained with biotin-coupled anti-CD115 antibody (Invitrogen) and coincubated with Streptavidin MicroBeads® (Miltenyi). Separation was done on magnetized LS columns (Miltenyi) according to manufacturer’s instructions.

### FACS analysis and sorting

Single cell suspensions were stained with the indicated antibodies according to manufacturer’s instructions for 30 min at 4°C and subjected to FACS analysis on a Gallios^TM^ (Beckman Coulter) or to cell sorting on Moflo^TM^ (Beckmann Coulter) or Aria Fusion^TM^ (Becton Dickinson).

For intracellular staining, BD Cytofix/Cytoperm^TM^ reagents were used according to manufacturer’s instructions and the eBioscience fixable viability dye^TM^ was used alongside surface staining. For BrdU-staining, the BrdU Flow Kit (BD PharMingen) was used according to the manufacturer’s instructions, cells were permeabilized and treated with 0.3 mg/ml DNase I (Roche) for 1 h at 37°C and stained for 20 min at RT with anti-BrdU antibody (BioLegend).

### Spectral flow cytometry

Skin was left floating in 2 ml of 2.4 mg/ml dispase (dispase II, Roche 10222100) solution for 1.5 h at 37°C, cut in the plate and transferred into 2 ml of 0.4 mg/ml collagenase (Sigma C5138-1G) and DNase I (0.04 mg/ml; Roche) solution for 1.5 h at 37°C and homogenized using a G18 needle and 5 ml syringe, then filtered. Single-cell suspensions were incubated with the primary surface antibody cocktail in PBS for 30 min at 4°C, washed, and incubated in the secondary surface antibody cocktail (fluorochrome-conjugated streptavidin) for 20 min at 4°C. Acquisition on a 5L Aurora spectral analyser (Cytek Biosciences). Flow cytometry data was analyzed using the FlowJo software (version 10.8.0, Tree Star Inc.) and Rstudio (version 4.0.1). Dead cell exclusion by using a Fixable Viability Kit (Zombie Nir™, Biolegend, Cat# 423105, dilution 1:400).

GM-CSF stimulation: Following surface staining, the cells were stimulated with RPMI/DMEM complete medium + 20 ng/ml mouse recombinant GM-CSF and incubated for 20-30 min at 37°C. Fixed in 2% PFA at RT for 10 min, then permeabilized in ice-cold methanol for 30 min on ice. Then stained with anti-PSTAT5 for 15 min at 4°C. 2^nd^ Ab staining mix-including Ab conjugated to methanol sensitive fluorochromes (PE, APC, PerCP and relative tandem dyes) - on top and incubate at 4°C for additional 20 min. Washed and filtered before acquisition.

The antibodies used for this study are listed in the key resources table.

### ELISA

ELISA analysis from lysed skin tissue was performed using the legendPLEX^TM^ kit Mouse Anti-Viral Response Panel (BioLegend) according to the manufacturer’s instructions. The samples were acquired on a Gallios^TM^ (Beckman Coulter) flow cytometer on the same day. Data was analyzed using the cloud-based Data Analysis Software (qognit™) provided by the manufacturer.

### Bacterial culture

BCG-RFP was previously described^20,28^. BCG-BFP was generated by transfection with a BFP expressing plasmid via electroporation (Addgene plasmid # 30177; http://n2t.net/addgene:30177; RRID: Addgene_30177)^29^. Fluorescent protein-expressing strains were expanded in 7H9 broth with 10% OADC and 0.5% glycerol containing 50 µg/ml Hygromycin up to an OD of 1. Pelleted cultures were resuspended in medium + 15% glycerol and stored at -80°C.

Tissue lysates were plated on 7H10 plates and serial dilutions were obtained to calculate CFU/organ.

*S. aureus* (*Newman*) was cultured in LB to mid-logarithmic growth. 10^6^ CFU per infection were used. Serial dilutions of organ lysates were cultured on blood agar to calculate CFU/organ.

### qPCR

For RNA extraction of sorted mac subsets, cells were sorted into RLT buffer containing 1% β-mercaptoethanol and RNA was extracted using the QUIAGEN RNeasy Micro kit^TM^ or the Blirt ExtractMe total RNA micro kit^TM^ according to manufacturer’s instructions. RNA was reversely transcribed to cDNA with the SuperScript™ IV VILO mix (Thermo Fisher). qRT-PCR was performed with Absolute qPCR SYBR Green (Thermo Fisher Scientific) at the Roche Lightcycler^TM^ in 384 well plates. Analysis of melt curves was implemented as a quality control and values were excluded if more than one Tm peak (Temperature with sharp drop in fluorescence) was detected.

The primers are listed in table 1.

**Table 1:**
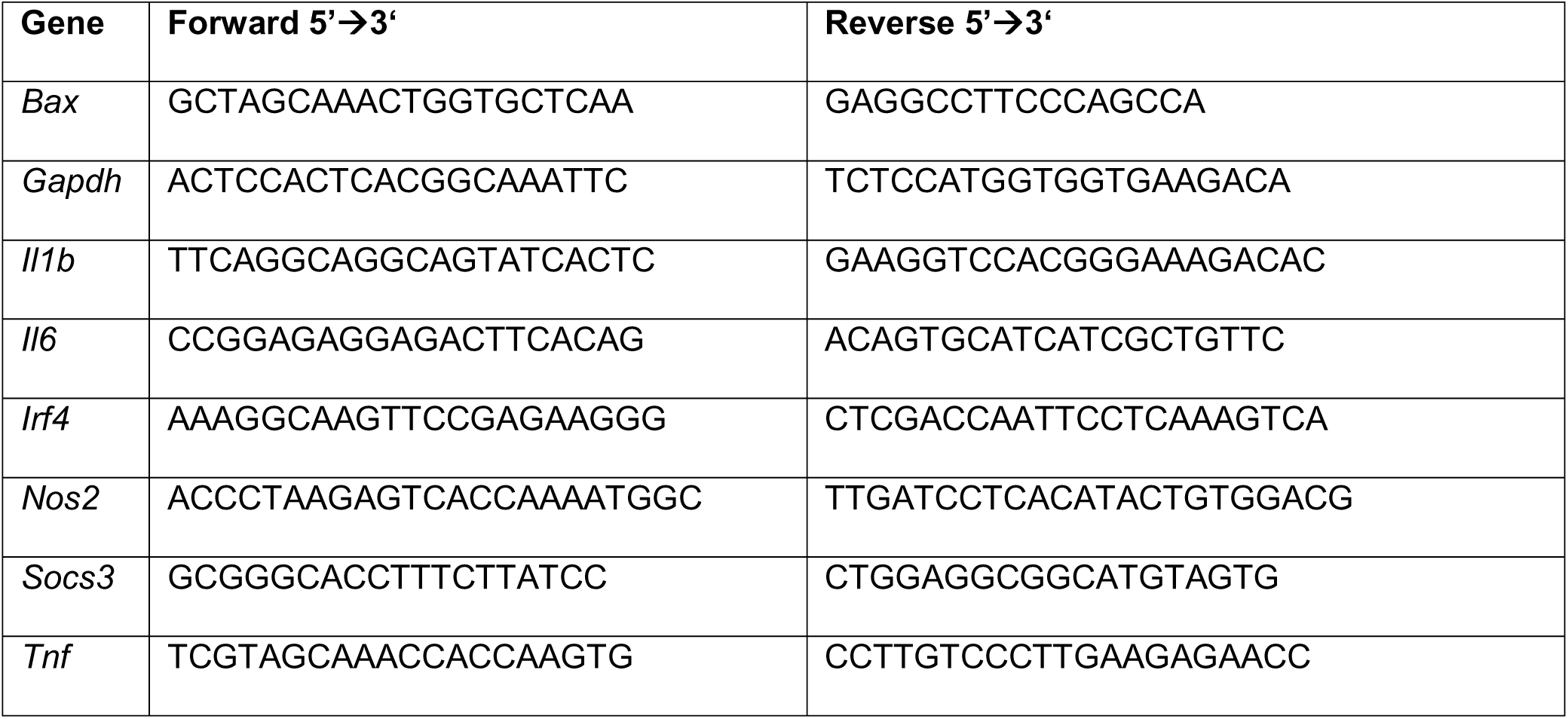
Primers used for qPCR analysis.

### Bulk RNA sequencing

Bulk RNA sequencing was performed by CeGat (Tübingen, Germany). There, whole transcriptome sequencing was performed by NovaSeq 6000 (Illumina) with 2 x 100bp read length and 50 million clusters/probe. All analyses were performed in R. Normalization, PCA plots and group comparisons were calculated with DESeq2. Shrinkage of effect size was performed with the lfcshrink function of the apeglm^30^ package. Gene set enrichment was done with the ClusterProfiler package, heatmaps were generated with the ComplexHeatmap package and volcano plots with the EnhancedVolcano package.

### Single-cell RNA sequencing

Single-cell RNA-sequencing was performed using the 10X 3’ mRNA-Sequencing kit v3.1 chemistry (cat # 1000269). Cells were multiplexed using the 10X 3’ CellPlex Kit Set A (cat # 1000261). Sequencing libraries were prepared according to the manufacturer specifications by the 10X Chromium Next GEM Single Cell 3‘ Reagents Kits v3.1 (Dual Index) Protocol with feature barcoding for cell multiplexing CG000388 Rev B utilizing the library construction (cat # 1000190) and 3’ Feature barcoding kits (cat # 1000262), for replicates 4-6 of the steady state data and for the data presented in figure 7. The gene expression and feature barcoding libraries were sequenced on an Illumina NextSeq 550 or a NextSeq1000/2000 machine with a minimum targeted sequencing depth of 20,000 reads per cell. Fastq files were aligned using the CellRanger software version 3.1 (replicates 1-3 of the steady state data presented in figure 1 and 2) or 6.1.2 (replicates 4-6 of the steady state data presented in figure 1 and 2 and data presented in figure 7).

**Figure 1:**
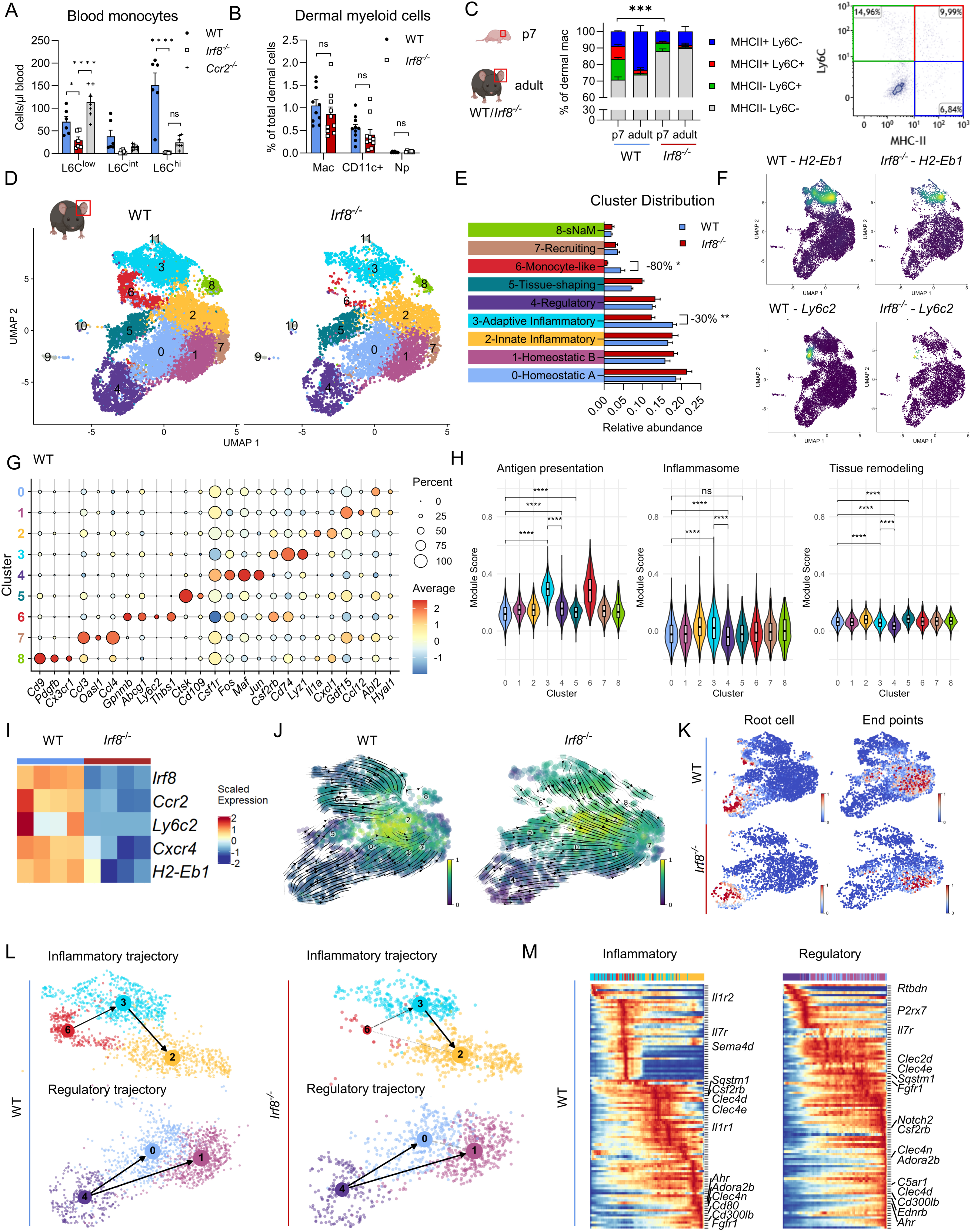
A: Quantification of blood Mc in WT, *Irf8*^-/-^ and *Ccr2*^-/-^ mice with flow cytometry. n=6-7 mice/genotype. B: Flow cytometry quantification of dermal myeloid cells as percentage of total dermal cells in WT and *Irf8*^-/-^ mice. n=10 mice. DC: Dendritic cell. Np: Neutrophil. ns: not significant. C: Flow cytometry data of Ly6C^+^ and MHCII^+^ subsets of dermal macs in young and adult WT and *Irf8^-/-^* mice. n= 7-8 mice (p7). Significance level refers to both Ly6C+ subsets respectively. D: Uniform Manifold Approximation and Projection (UMAP) representation based on gene expression profiling of sorted dermal macs of WT (left) and *Irf8*^-/-^ (right) mice. Cells from 6 mice of two independent experiments are pooled. E: Mac identities of each cluster based on expression of marker genes and gene set enrichment analysis along with the relative quantitative cluster contribution. sNaM: sensory nerve associated mac. F: Density plots displaying scaled expression levels of an exemplary MHCII-associated gene and *Ly6c2*. Blue:low, yellow:high. G: Dot plot showing key marker genes differentially expressed in clusters 0-8 of WT. Color: mean expression in the respective cluster (“Average”); Dot size: Fraction of cells expressing the gene (“Percent”). H: Violin plots showing module scores of gene sets associated with the indicated function. Wilcoxon test compares between relevant clusters. Box plots depict median with IQR. The gene sets are listed in table S9. I: Heatmap displaying exemplary genes that are significantly differentially expressed between WT and *Irf8*^-/-^ mac assessed by pseudobulk on the scRNAseq dataset using Deseq2. All 129 DEGs are listed in supp. Table 3. J: VeloVI RNA velocities derived from the dynamical pseudotime model projected onto the UMAP representation. K: UMAP representation showing the “root cells” (left, in red) and “end points” (right, in red) identified by dynamic RNA velocity analysis. L: PAGA velocity directed graph superimposed onto the UMAP representation of the “inflammatory” (above) and “regulatory” (below) differentiation trajectory. WT:left; *Irf8^-/^*^-^:right. M: Heatmap displaying top 100 dynamically expressed genes along the inflammatory and regulatory trajectories. Signaling receptors are labelled. Blue:low; red:high. Statistics in figs. 1B-C, E: students t-test including Holm-Šídák-correction; 1A: Two-way ANOVA.

Downstream analyses were conducted in the R version 4.2.2 programming environment using the Seurat package version 5 (except clustering and UMAP generation for the steady state and BMTX dataset was done using Seurat package version 4) or with Python 3.11 using the VSC interface.

All data sets were filtered to remove cells with more than 5% mitochondrial transcripts, with less than 200 or more than 5,000 detectable genes. Furthermore, to remove potentially contaminating dendritic cells or mast cells, cells expressing more than two transcripts of either *Zbtb46* or *Cpa3* were removed. The dataset after transplantation (fig. 5) was further filtered by removing one non-identifiable outlier cluster consisting of 45 cells via the UMAP coordinates. A total of ∼ 17,000 cells in steady state, of ∼ 4,000 cells after BMTX, and of ∼21,000 cells after BCG infection was included in the final analysis.

WT and *Irf8^-/-^* steady state data were integrated from two separate experiments using the SCtransform algorithm within Seurat (v5). 2000 features were identified using SelectIntegrationFeatures and used in PrepSCTIntegration and FindIntegrationAnchors. These anchors were used on IntegrateData. No Integration procedure was applied to the BMTX and BCG infection datasets and 3000 variable features were utilized. The top 30 PCs (steady state), 19 PCs (BMTX) and 30 PCs (BCG infection) were used for FindNeighbors calculation based on the result of the respective elbow plots as suggested by the package developers. Resolutions to find clusters were set to 0.45 (steady state) 0.64 (BMTX) and 0.6 (BCG infection), based on distinguishability of the resulting clusters.

Core markers were identified using the FindAllMarkers function and filtered based on significant p-values. Gene set enrichment on GO terms was done using the ClusterProfiler v4.12.6 package function gseGO with minimal and maximal gene set sizes of 5 and 800 respectively. The figures display exemplary GO terms. AddModuleScore was used to analyze involvement of a selection of genes listed in supplemental table 9. Pseudobulk analysis was performed using the AggregateExpression function on conditions, followed by the FindMarkers function and significance testing using DEseq2. The CellChat package (v1.6.0) was utilized using the default settings, for computeCommunProbPathway the data was filtered with a threshold of minimal 10 cells.

To formally compare similarity of clusters between data sets, using the quadratic programming approach, we calculated weights for all cluster medoids in one dataset for each cell in the other dataset using the solve.QP function of the quadprog R package. For example, to calculate the weights for all cluster medoids of the WT data for each cell in the BMTX data, the Seurat object was converted into the RaceID3^31^ object and the WT normalized transcript count matrix containing genes as rows and single cells as columns and the BMTX normalized transcript count matrix containing genes as rows and cluster medoids as columns were provided as inputs to the Dmat and dvec arguments of solve.QP function, respectively. Cluster medoids were calculated by the compmedoids function of the RaceID3 algorithm. The intersect of top 3,000 variable genes expressed in both datasets was used for calculating the weights using quadratic programming.

RNA velocity and PAGA analysis were done using the Python scvelo, scanpy, and VeloVI packages. The data was filtered and normalized using default settings. Velocities were calculated using scv.tl.velocity with mode set to ‘dynamical’. Pseudotime was calculated using the scv.tl.velocity_pseudotime command with default settings. Splicing dynamics were calculated by the scv.tl.recover_dynamics function. Velocity graph, root cells and end points and latent time was calculated by scv.tl.velocity_graph with mode_neighbors set to ‘connectivities’, scv.tl.terminal_states and scv.tl.latent_time after subsetting to the data to clusters 0-8. PAGA was performed using scv.tl.paga with groups set to mac clusters. Next, RNA velocity analysis was conducted using the VeloVI framework, a VeloVI model was instantiated and trained on the dataset to learn the underlying transcriptional dynamics. After training, RNA velocity estimates were extracted using the model.get_velocity function, and the velocity values were used for visualization.

### Spatial transcriptomics

Spatial transcriptomics were performed on sections of 8 halves of four ears stacked on top of each other per condition. Prior to sectioning, ears were formalin fixed and paraffin embedded (FFPE). Ears were arranged side by side to obtain transversal sections of several biological replicates. Spatial gene expression was assessed using the VisiumHD platform according to the manufacturer’s protocol with minor adjustments. 5 µm sections were placed on a high-adherence microscopy slide (Trajan Series 3, Trajan Scientific and Medical, Germany). Deparaffination and staining was performed according to the 10x Genomics Tissue preparation guide (CG000240 RevE). High-resolution tissue imaging was performed on a Keyence BZ-9000 system. For spatially-resolved transcriptome quantification, the Visium HD v1 – FFPE reagents were used along with Visium Mouse Transcriptome Probe Set v2.0. Probe release and library preparations were conducted following the manufacturer’s instructions (CG000685, RevA). The libraries were sequenced on an Illumina NextSeq 1000/2000 system (Illumina, Inc., San Diego, United States). Next, we aligned high-resolution images with the fiducial frame using Loupe Browser v8.1.0 (10X Genomics). Sequencing data along with the image alignment results were passed to Spaceranger v3.0.0 (10X Genomics) with the mm10 mouse reference transcriptome downloaded from the 10X Genomics website (mm10-2020-A). Downstream analysis was performed with Seurat v5 for clustering and gene expression analyses and SPATA2^32^ for localization analysis. For annotation of clusters, cell identity decomposition was done using the spacexr RCTD algorithm^33^ on a published reference dataset of total dermal skin cells^34^ merged to our homeostatic dermal mac dataset, creating a subset of cells assigned as antigen presenting cell/mac (APC-MAC). These were deconvoluted in a second step to the most likely cluster identity using our WT homeostatic scRNAseq dataset. Tissue borders and hair bulbs were manually assigned and used for distance analyses. Distance to border was compared with Wilcoxon test between clusters. Enrichment near hair bulb was defined as 8 µm or closer and comparisons were calculated with fisher’s exact test. Neighboring cells per spot were defined by setting a spatial radius to a value, where a mean of 6 neighboring spots were considered per mac spot. The respective mac spot itself was excluded from the neighbor analysis. The heatmap displays the top 10 differentially expressed genes (DEGs) per cluster. Enrichment of neighboring cell types was analyzed using the Seurat BuildNicheAssay function, 20 neighbor spots per central spot were used for enrichment analysis, either around cell type or around mac cluster.

### Mouse skin microscopy

For the generation of cryosections, ears were fixed for 4 h in 4% PFA and transferred to 20% sucrose overnight. Ears were embedded in Tissue Tek^TM^ (Sakura), frozen in liquid nitrogen and stored at -80°C until further processing. Samples were cut into 8 µm sections on a cryotome at - 20°C. Blocking and permeabilization steps were done in wash buffer (1% BSA, 0.1% Triton X100), followed by blocking with Fc block (CD16/32) for 30 min, and incubation with antibodies overnight. Mounted with ProLong^TM^ Diamond mounting medium and analyzed with a LSM 710 or LSM 880 confocal microscope equipped with a 20 × /0.8 NA Plan-Apochromat objective (Carl Zeiss Microimaging).

### Human skin microscopy

Paraffin-embedded skin samples from female patients with graft versus host disease or mycobacterial infection who previously underwent sex-mismatched transplantation were obtained. 5 µm thick sections were used for combined FISH-IF for simultaneous detection of Y chromosome, CD68 and iNOS. At first, the sections were deparaffinized and rehydrated by serially immersing them in Xylol (2 times, 10 min each), 100% isopropanol (2 times, 5 min each), 96% ethanol (5 min), 70% ethanol (5 min) and distilled water (5 min). Antigen retrieval was performed in sodium citrate buffer for 7 minutes in a pressure cooker. Once the slides cooled down, they were washed in 1X PBS and treated with pre-warmed pepsin at 37°C for 3 min, followed by a brief wash with 1X PBS and fixation in 1% paraformaldehyde. Fixed skin sections were dehydrated in ascending concentrations of ethanol (70%, 96% and 100% ethanol, 1 min each). After briefly air drying the slides, 20 µl of anti-Y probe mix (14 µl of hybridization buffer, 2 µl probe and 4 µl distilled water) was added to cover each of the sections and incubated at 80°C for 3 minutes, followed by incubation at 37°C overnight. The next day, the slides were washed sequentially in 1X SSC (supplemented with 0.05% Tween-20) and 0.4X SSC at 72°C for 3 minutes. After blocking with 10% donkey serum, incubation with the primary antibody mix (anti-CD68-Alexa Fluor 488 and anti-iNOS) was done overnight at 4°C. Afterwards, following three brief washes with 1X PBS, a secondary antibody against iNOS was applied to the slides for one hour at room temperature. After brief washing, the sections were incubated with DAPI (1:500) for nuclear counterstaining for 5 minutes at room temperature. Washed and air-dried slides were mounted in VectaShield medium and stored at 4°C. Images from representative areas from each section were acquired as Z stack with a Nikon LSM800 confocal microscope (40X objective). Maximum intensity projection of the Z-stack images was performed using image processing package FIJI from ImageJ (release 2.13.1). A cell was considered positive for Y-chromosome only when the signal overlapped with DAPI. Counting was performed manually using the multi-point tool in FIJI.

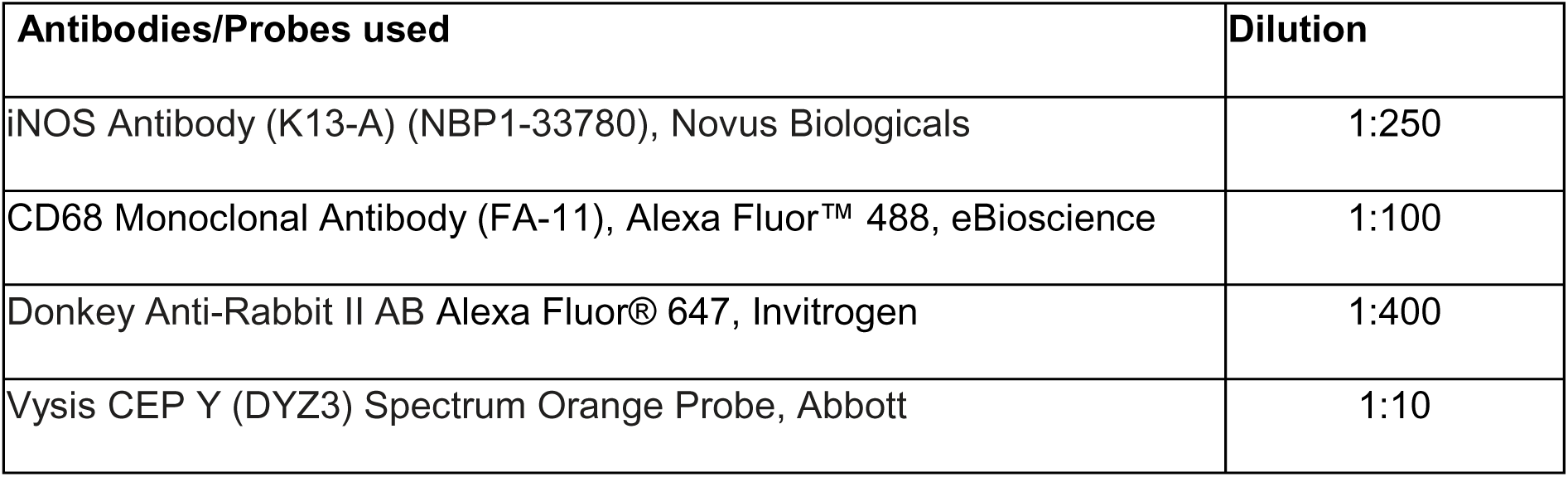

| Antibodies/Probes used | Dilution |
| --- | --- |
| iNOS Antibody (K13-A) (NBP1-33780), Novus Biologicals | 1:250 |
| CD68 Monoclonal Antibody (FA-11), Alexa Fluor™ 488, eBioscience | 1:100 |
| Donkey Anti-Rabbit IgG Alexa Fluor® 647, Invitrogen | 1:400 |
| Vysis CEP Y (DYZ3) Spectrum Orange Probe, Abbott | 1:10 |

### Cell culture experiments

For bone marrow-derived mac (BMDM) generation bone marrow cells were resuspended in DMEM medium complete (containing 10% FBS (Invitrogen), 10 µg/ml ciprofloxacin (Fresenius Kabi) and 20 ng/ml M-CSF (Peprotech)) and kept at 37°C and 5% CO_2_ for 5 days. Multinucleated giant cells were generated as described previously^20^. In short, BMDM were seeded in Opti-MEM GlutaMAX (Gibco) supplemented with 10% FBS and either with 50 ng/ml M-CSF alone, or plus BCG (MOI 20), FSL-1 [20 ng/ml (Invivogen)], or heat-fixed *M. tuberculosis* [10 µg/ml (InvivoGen)] for six days on 96 well plates [CellBind Surface (Corning)] for 6 days. Staining was performed using Hemacolor® staining of adherent cells according to manufacturer’s instructions. Images were acquired with Brightfield microscopy with at least 9 fields per condition manually quantified. Phagocytosis assay was performed with BMDM seeded in DMEM medium containing 10% FBS (Invitrogen) and 10 µg/ml ciprofloxacin (Fresenius Kabi). Either fluorescent beads (1 µm) or BCG-RFP (MOI 20) were added on the culture for the indicated time period, the negative control was supplemented with 50 µg/ml Cytochalasin B (Sigma). Then cells were put on ice for scraping, washing and surface staining.

For 3D cell culture a PEG (Polyethylenglycol)-based hydrogel with 8-arm PEG polymers was used^35^. 40 kDa PEG polymers, functionalized with vinyl sulfone groups, were manufactured by NOF Europe, Germany. Hydrogels were synthesized with linker-peptides containing a cysteine on each site of the peptide to react with the PEG polymers via a Michael-addition. For functionalization of the PEG hydrogels, RGD-peptides were included into the hydrogel setup. The peptides were manufactured by GenScript, USA. They contained an N-terminal Acetylation and a C-Terminal Amidation and had a purity of >90%.

The amino acid sequences were:

<u>Linker-peptide</u>: GCRD-LEVLFQGP-GPQGIWGQ-DRCG

<u>RGD-peptide</u>: GCGYGRGDSPG

BMDM were prepared as mentioned above, PEG and 0.3M Triethanolamine buffer were added to the cell suspension, the RGD-peptide was added and incubated for 5 min at room temperature. To start hydrogel synthesis the linker-peptide was added and 10 µl hydrogel solution was pipetted on a hydrophobic coated glass slide (using Sigmacote®) and then incubated for 10 min at room temperature. The solidified hydrogels were transferred into a 96-well plate containing DMEM medium complete, cultured for 5 days, then gels were cut into small pieces and cells were lysed in the gels, then subjected to qPCR analysis.

### Kinetic modeling

For modeling the kinetics of differentiation and exchange of dermal macs, a differential equation model with 8 dynamic states (representing resident macs, resident radioresistant macs, as well as circulating Mc, Ly6C^hi^ and Ly6C^low^ from donor and recipient) was applied. For the recipient cells, we used steady states as initial conditions. After transplantation and irradiation, recipient cells were replaced by donor cells, i.e. non-resident recipient cells were set to zero, radiosensitive resident cells quickly degraded with an estimated rate for the unshielded condition and donor cells populated the Ly6C^hi^ and Ly6C^low^ states. All unknown parameters, including the proportions of radiosensitive and radioresistant resident macs, 6 rate constants for production, turnover and degradation, 2 feedback parameters, and 9 SDs for the error of the measured quantities were jointly estimated from the data for both shielding conditions via maximum likelihood estimation. Equations and further details about this ordinary differential equation model are provided as supplemental document 1.

For estimating half-lives of the exchange of the cells in clusters derived from single-cell sequencing, we used a two-state dynamic model with one state for donor cells and one state for recipient cells and two unknown, cluster-dependent rate constants for inflow and outflow. As initial conditions, we again used steady states which were here given by the ratio of the two rate constants. A joint SD of the measurement errors in all clusters was assumed to prevent overfitting.

For both models, the Data2Dynamics modelling toolbox^36^ was used to numerically solve the differential equation models, for estimating the parameters^37^ and for performing uncertainty analyses^38^.

### Transcription factor regulon analysis

To conduct the SCENIC analysis, we employed an in-house pySCENIC pipeline^39^, focusing on a subset of (WT) cells from our homeostatic scRNAseq data. The subsetting parameters were set to a value of 5, and the analysis was performed over 50 iterations to ensure robustness in the results. This approach enabled the identification of transcription factors (TFs) and their associated regulons. Upon completion of the SCENIC analysis, we merged all identified regulons into a single unidirectional regulatory network (GRN). Given that SCENIC is based on a gradient boosting framework, the results — specifically the TF-gene pairs constituting the network — exhibited a stochastic nature. Consequently, we filtered the GRN to retain only those connections with more than 15 hits between TFs and genes. This comprehensive network served as input for the subsequent GRaNPA analysis, which evaluates the biological significance of the identified TFs.

We then performed GRaNPA (Gene Regulatory Network Performance Analysis)^40^, which assesses how effectively the bipartite TF-gene connections within our GRN can predict cell-type-specific differential gene expression. Simultaneously, this framework identifies the TFs that are critical for making accurate predictions. For the GRaNPA analysis, we utilized differentially expressed genes obtained from the bulk RNA sequencing dataset (WT Res-mac (*Irf8*^-/-^ BM) versus WT Res-mac (WT BM), fig 4). The gene list was filtered to retain only those genes with a false discovery rate (adjusted p-value) < 0.2 and a log fold change |≥ 0.4|.

### Statistical analysis

Statistical analysis was performed with Prism^TM^ 10 (GraphPad). For comparison of two groups, unpaired two-tailed Student’s t test was applied unless stated otherwise. For serial comparisons within one graph correction according to Holm-Šídák was performed for multiple t-tests. For multiple comparisons, one-way or two-way ANOVA was performed, as applicable, followed by Tukey’s or Dunnett’s multiple comparison tests and adjusted p-values are displayed. Transcript counts or counts of aggregated gene scores were compared using the non-parametric Wilcoxon-test. Differences were considered statistically significant if p-values were ≤ 0.05. (ns, not significant; *p < 0.05; **p < 0.01; ***p < 0.001; ****p < 0.0001). Data are represented as mean ± SEM if not stated otherwise. Sample sizes are indicated in the figure legends.

### Data availability

All R and python scripts are deposited online at https://github.com/lohrmanf.

All raw sequencing data is publicly accessible at GEO.

| Experiment | Main presenting figure | GEO number |
| --- | --- | --- |
| scRNAseq homeostasis | 1 |  |
| Visium HD Spatial seq homeostasis | 2 |  |
| BulkRNAseq density control | 4 |  |
| scRNAseq transplantation | 5 |  |
| BulkRNAseq infection | 7 |  |
| scRNAseq infection | 7 |  |

### Supplemental information

Supplemental figures 1-7

Supplemental document 1: Equations for dynamic modelling

Supp. table 1: Differentially expressed genes per cluster WT

Supp. table 2: Differentially expressed genes per cluster *Irf8^-/-^*

Supp. table 3: Differentially expressed genes WT versus *Irf8^-/-^* (pseudobulk)

Supp. table 4: Differentially expressed genes per cluster transplantation

Supp. table 5: Differentially expressed genes per cluster uninfected WT

Supp. table 6: Differentially expressed genes per cluster infected WT

Supp. table 7: Differentially expressed genes per cluster uninfected *Irf8^-/-^* Supp. table 8: Differentially expressed genes per cluster infected *Irf8^-/-^*

Supp. table 9: Gene sets used for gene scores, figures 1 and 5

Data2Dynamics_modeling_report

## Key resources table

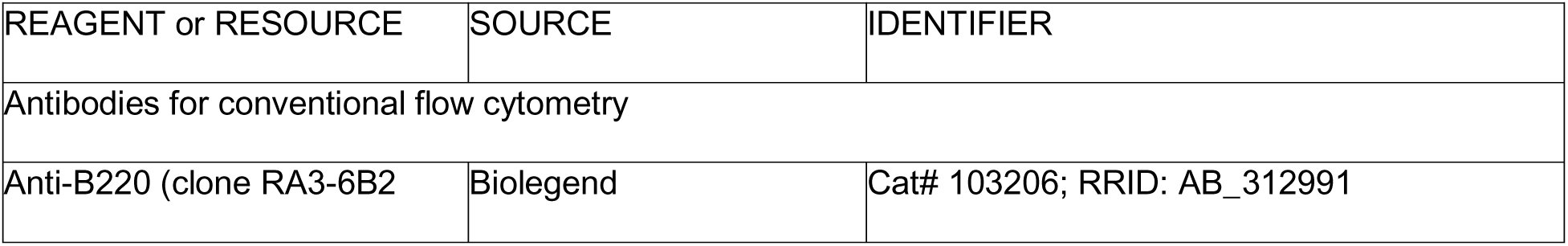
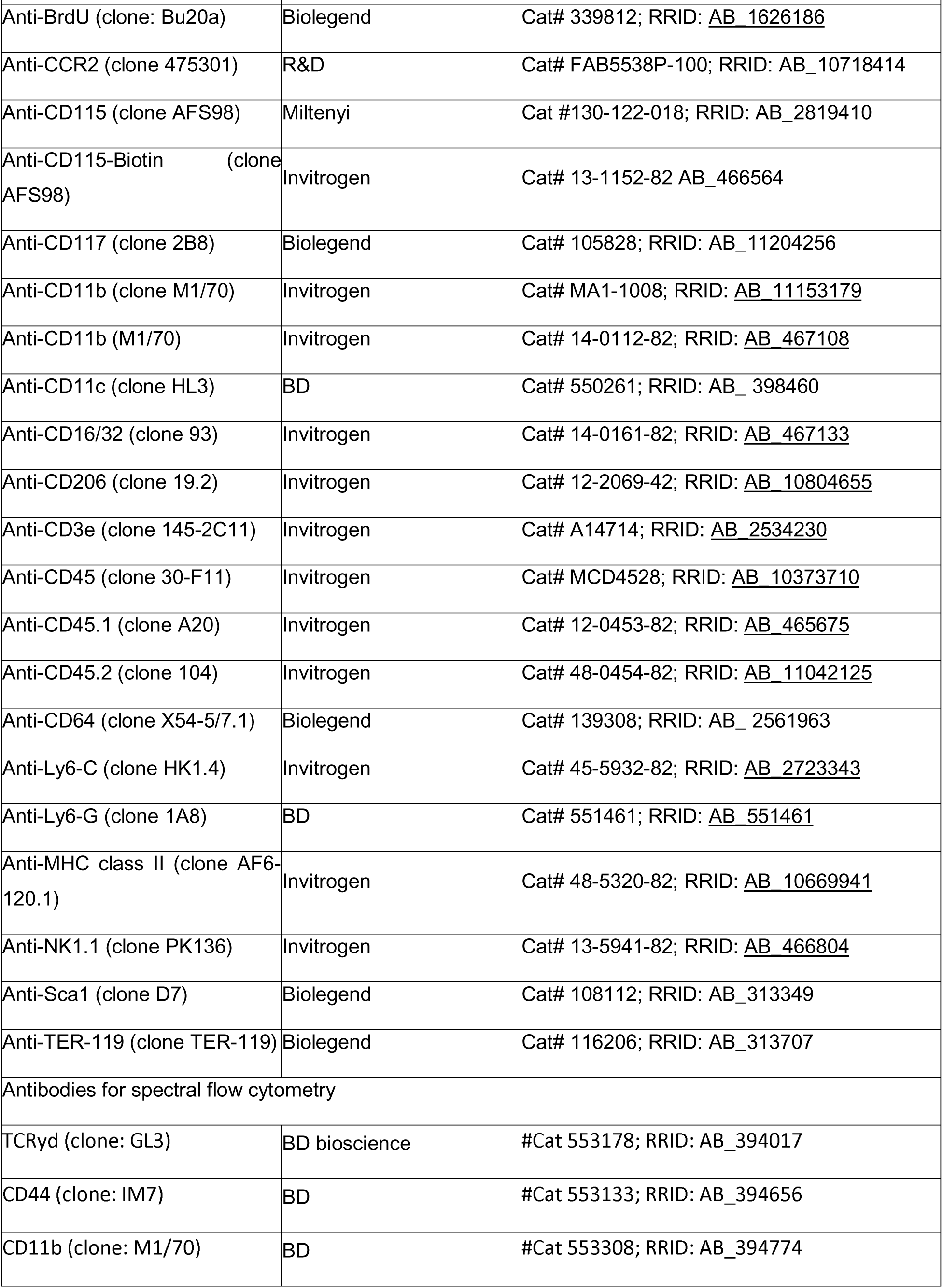
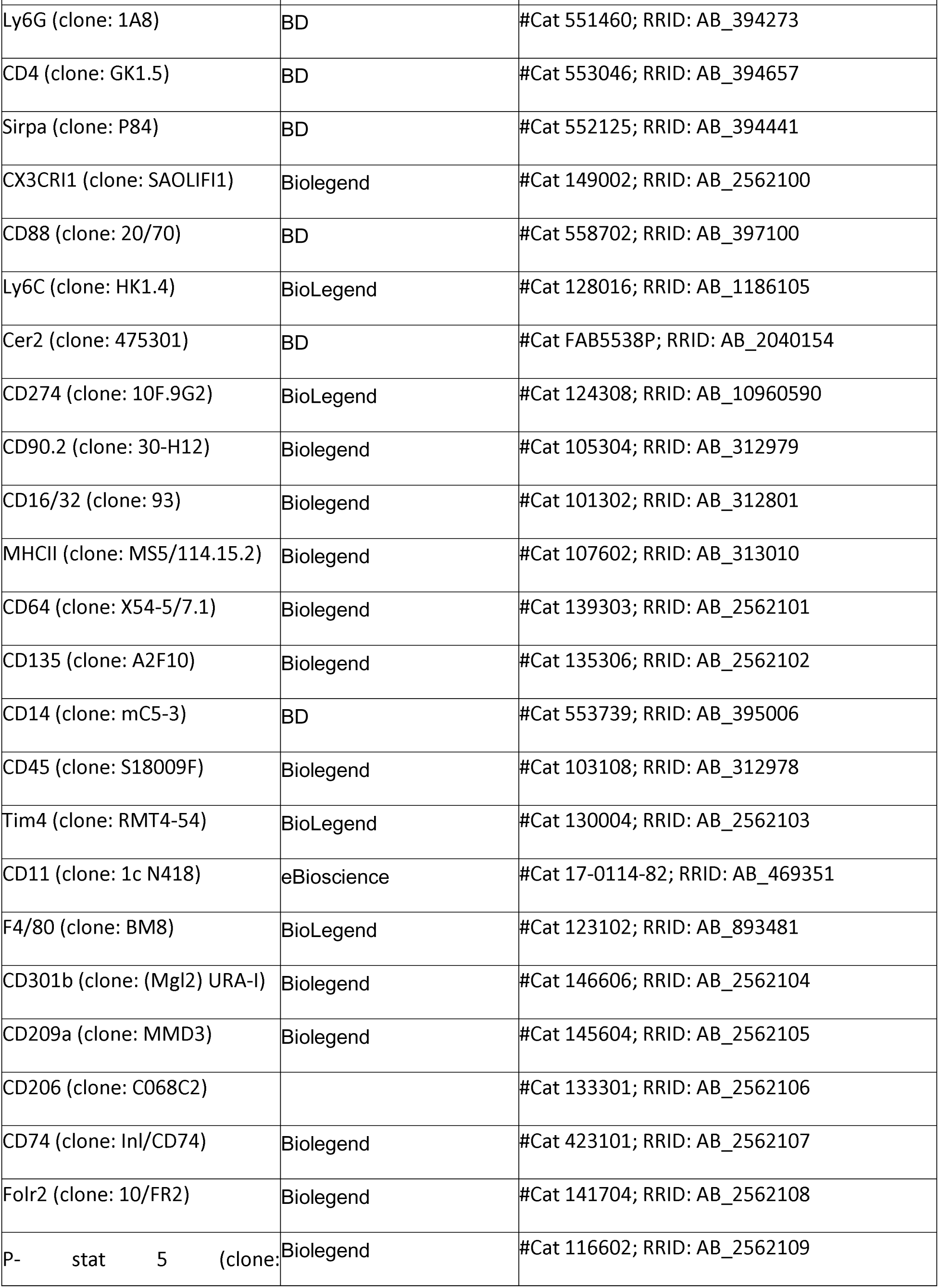
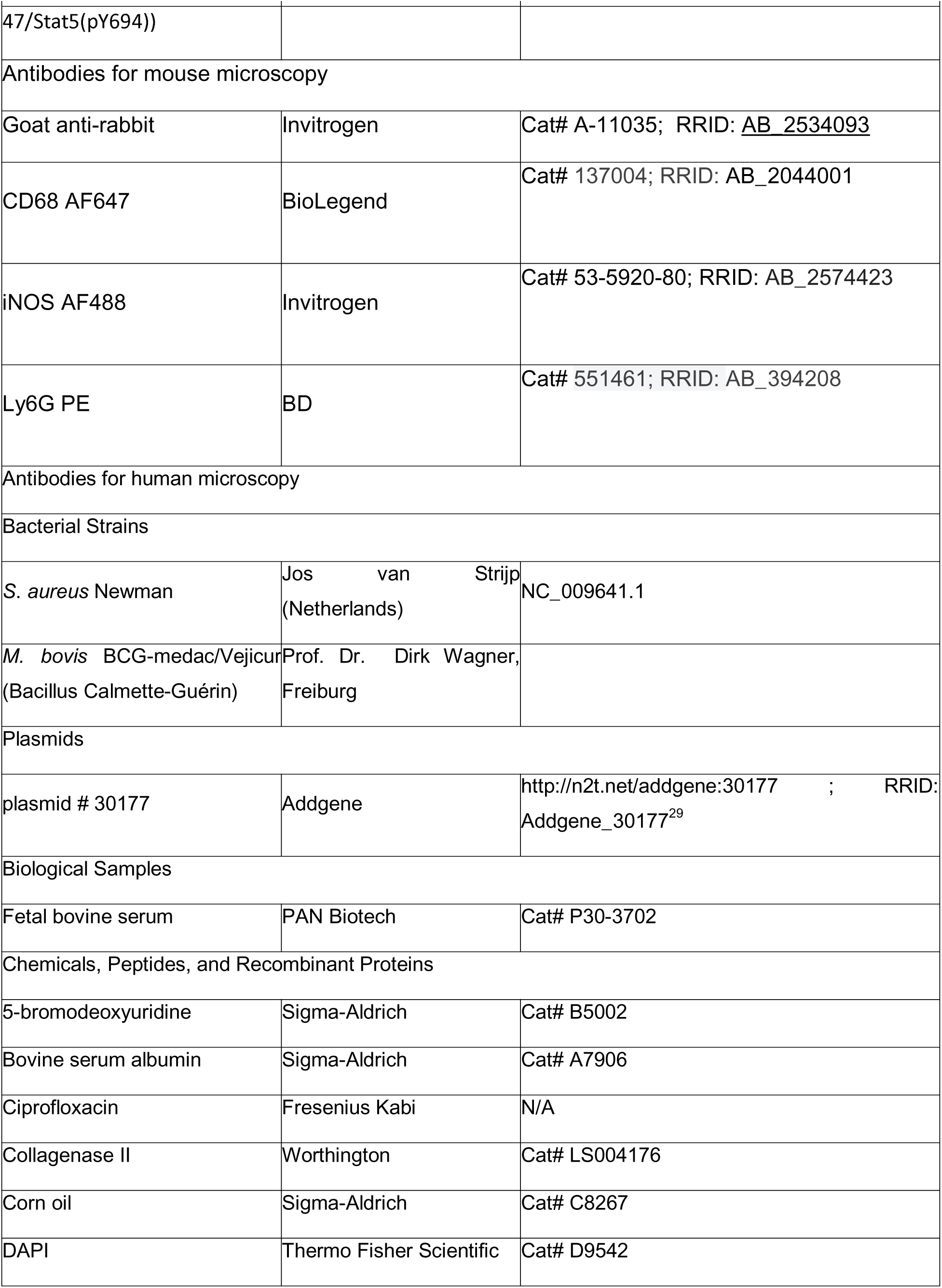
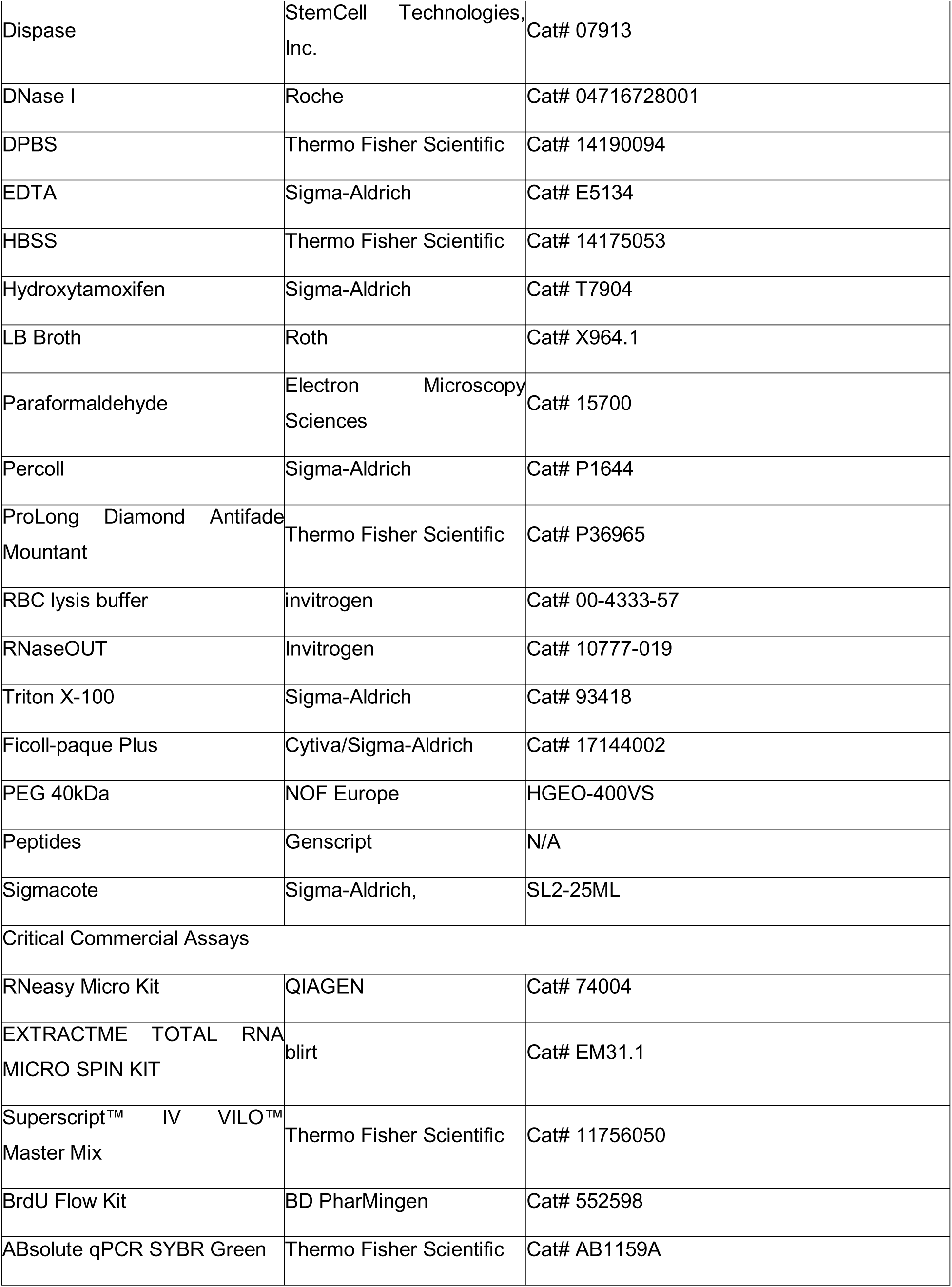
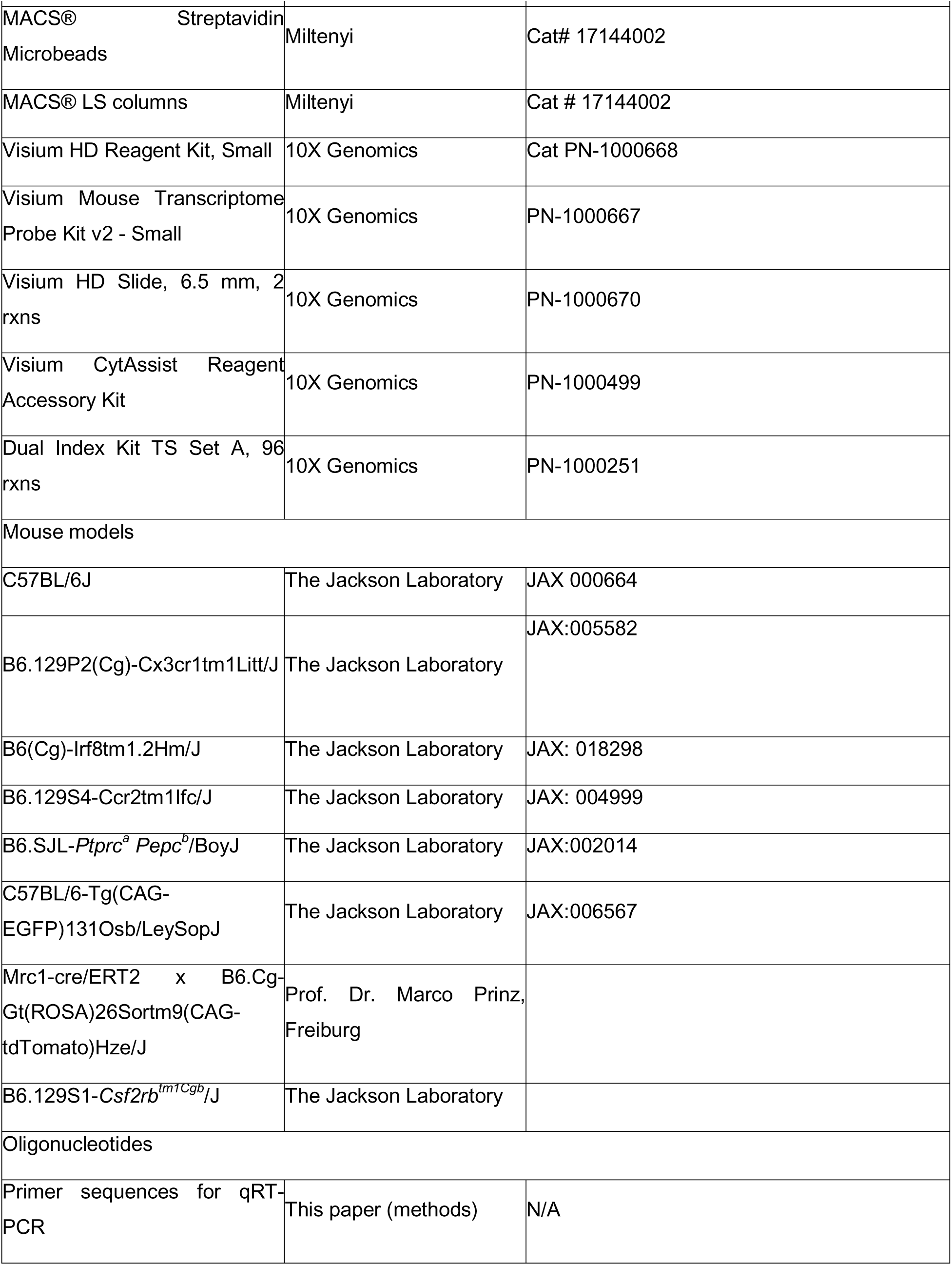
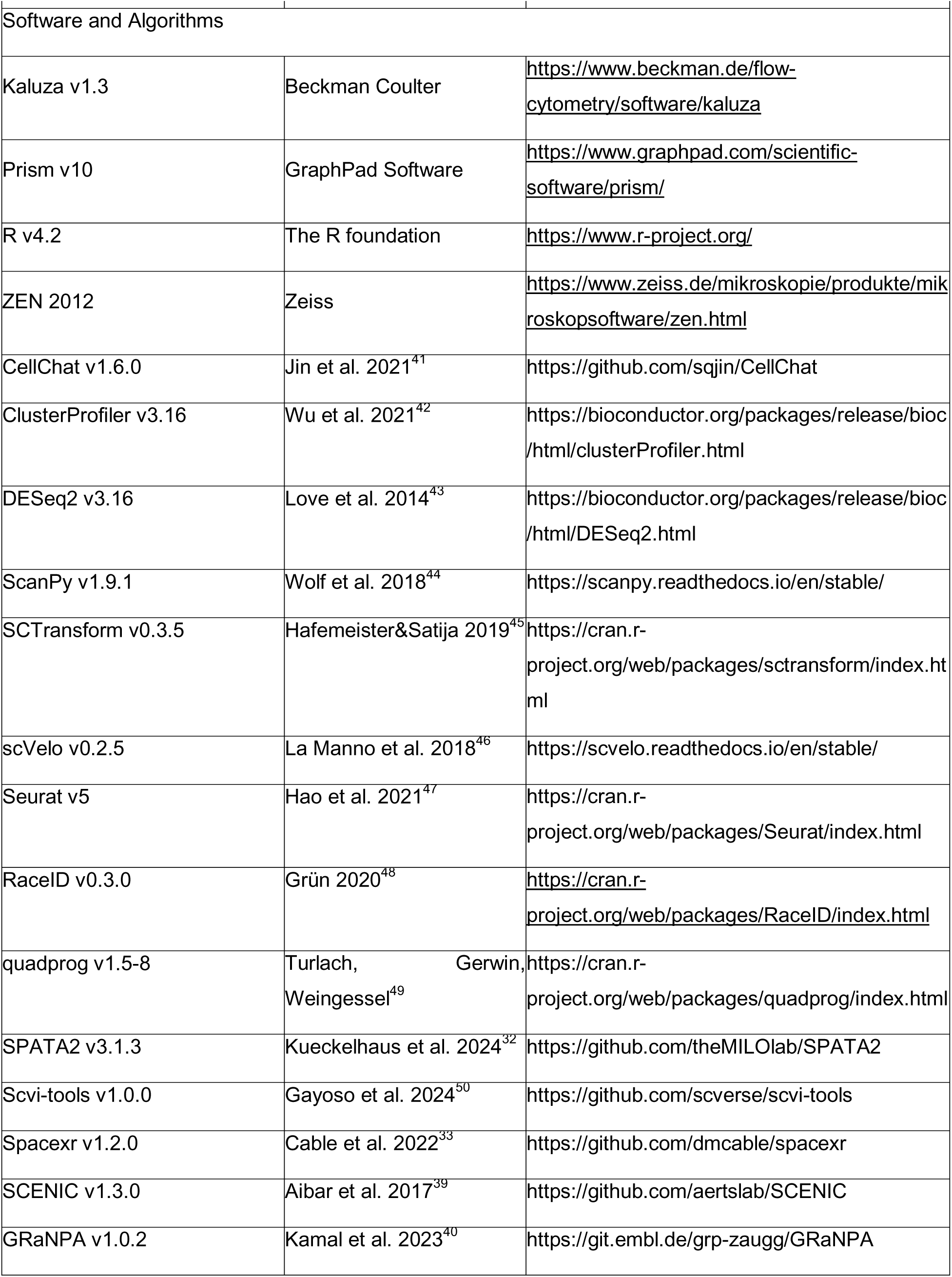

## Results

### Resident macs compensate for the lack of monocytes to maintain density and subset composition

*Irf8* mediates proper myeloid differentiation in the bone marrow and thus Mc formation. In line with this and previous findings^2,51,52^*, Irf8^-/-^* mice displayed an almost complete lack of circulating Mc, with an accumulation of committed Mc progenitors in the bone marrow that prematurely express CCR2 (figs. 1A and S1.1A&C-E). In contrast, flow cytometry revealed similar numbers of dermal macs in the adult skin in *Irf8^-/-^* and wild type (WT) mice (defined by CD64^hi^, CD45^+^, CD11b^+^, CD11c^-^ and CD3^-^, figs. 1B and S1.1B). In WT mice, a high number of dermal macs was replaced by Mc-derived macs in the first postnatal week, as reflected by Ly6C positivity^1^ and partly MHC-II expression, whereas these populations were less abundant in *Irf8^-/-^* mice (figs. 1C and S1.1B). The influx of Ly6C+ cells persisted, albeit at a reduced rate, into adulthood in WT mice (fig. 1C). In *Cx3cr1^+/gfp^ Irf8^-/-^* mice, Ly6C^+^ macs were significantly reduced, similar to CX_3_CR1^int^ macs, which were previously shown to be the immediate progeny of Mc^2^ (fig. S1.1F). In accordance with a partial role for recently immigrated macs^13,15^, *S. aureus*-infected *Irf8^-/-^* mice exhibited a slightly reduced expansion of dermal macs, and reduced numbers of Ly6C^+^, i.e. Mc-derived, macs (figs. S1.1G-H). However, clearance was only slightly decreased (fig. S1.1I), indicating a mostly functional resident dermal mac compartment in the absence of Mc-derived macs. Overall, BrdU incorporation revealed higher proliferation rates of dermal macs in *Irf8* deficiency (fig. S1.1J).

To address the conundrum of stable mac density despite lacking Mc in *Irf8* deficiency, we performed single-cell RNA sequencing (scRNAseq) of ∼18,000 sorted dermal macs from WT and *Irf8^-/-^* mice. Unsupervised clustering using Seurat^47^ identified 9 clusters in the combined analysis (figs. 1D-E and S1.2A), which we annotated according to their presumed identity based on the analysis of core marker genes (figs. 1F-G and S1.2B-C&E), gene set enrichment analysis (GSEA) on gene ontology (GO, fig. S1.2D) and module scores of gene sets associated to antigen processing and presentation, inflammasome activity and tissue remodeling (figs. 1H and S1.2F). In short, Clusters 0 and 1, termed “homeostatic macs,” resembled resident macs with suppressed immune functions, cytokine production, and locomotion. Cluster 2, “innate– inflammatory macs,” showed high chemokine and cytokine expression and gene sets linked to acute inflammation and pattern recognition. Cluster 3, “adaptive–inflammatory macs,” expressed MHC-class II assembly genes, *Il1b*, and *Csf2rb*, with GO terms enriched for antigen presentation and adaptive immune cross-talk. This cluster was reduced by ∼30% in *Irf8^-/-^* mice. Clusters 2 and 3 displayed modest proliferation (*Mki67,* fig. S1.2E). Cluster 4, “regulatory macs,” expressed *Jun*, *Fos*, *Maf*, and *Egr1*, associated with alternative activation, and showed enriched GO terms for phagocytosis and tissue development. Cluster 5, “tissue-shaping macs,” expressed *Ctsk*, indicative of tissue remodeling activity. Cluster 6, “Mc-like macs,” nearly absent in *Irf8^-/-^* mice, expressed *Ly6c2* and *Ccr2*, with genes linked to immunity, phagocytosis, and locomotion. Cluster 7, “recruiting macs,” expressed *Ccl4* and showed enrichment for innate immune responses. Cluster 8, “sNaM,” expressed *Cx3cr1*, *Cd9*, and *Axl*, aligning with sensory nerve-associated macs^2^. Table 1 summarizes the defining features of all clusters.

**Table 1.**
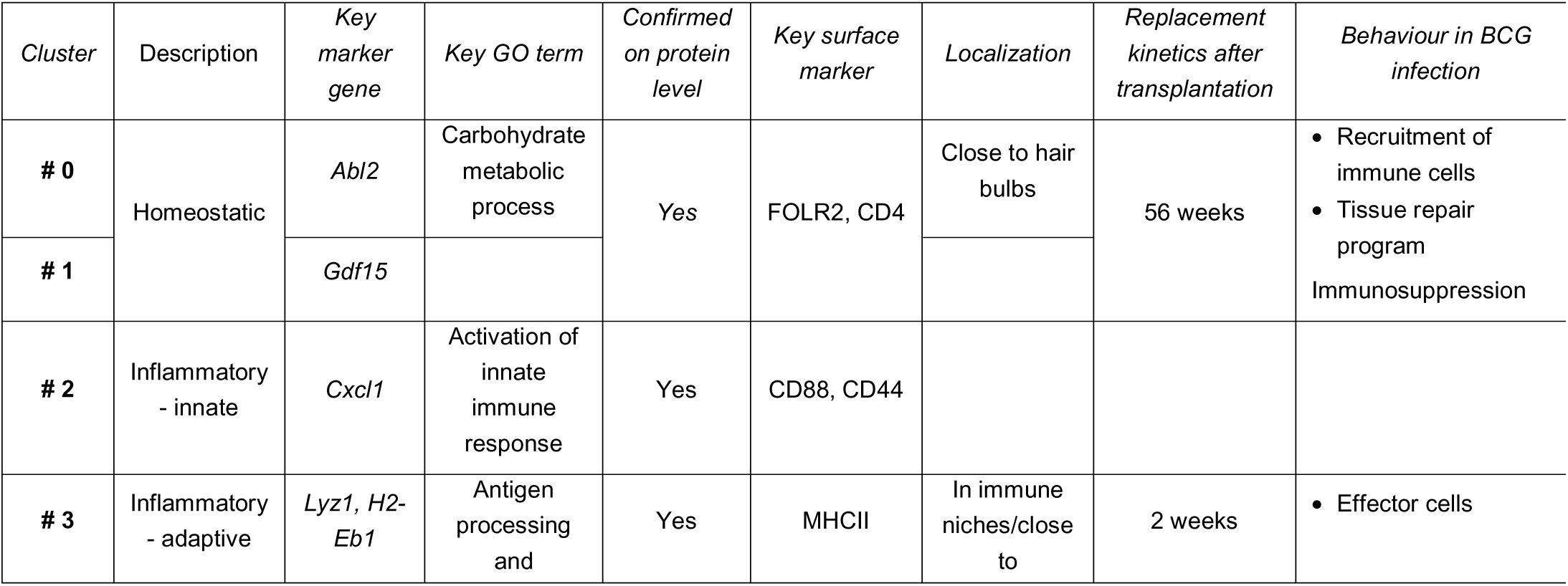
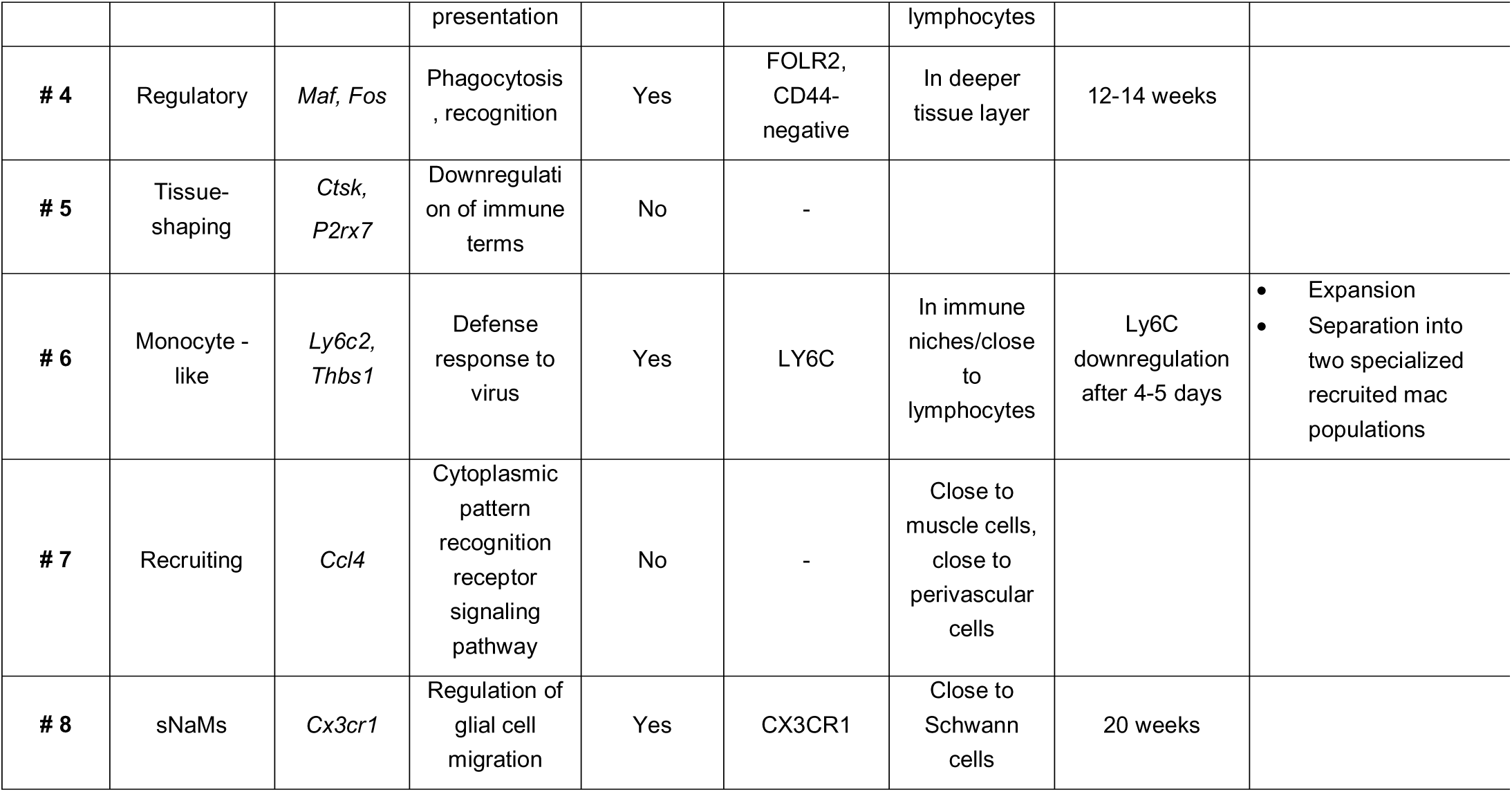

Of note, comparison of the total mac compartment of WT and *Irf8^-/-^* mice via pseudobulk analysis revealed 129 differentially expressed genes (table S3). Examples include reduced expression of MHC class II-related genes, and genes associated to a mc-phenotype including a virtual absence of *Ly6c2* expression in *Irf8^-/-^* mice (fig. 1I). Module scores of gene sets showed only minor deficiencies, mostly limited to antigen processing and presentation (fig. S1.2F), supporting a mostly intact and functional dermal mac compartment in the homeostatic dermis of *Irf8^-/-^* mice.

We delineated the transcriptional program employed by precursor populations differentiating into specific mac subsets utilizing the scVelo/VeloVI algorithm, which infers differentiation trajectories based on the kinetics of the mRNA lifecycle^46,50^. We identified two distinct differentiation trajectories of dermal macs (fig. 1J). One trajectory started with Mc-like and adaptive – inflammatory macs and gave rise to innate – inflammatory macs, while another started with regulatory macs and gave rise to homeostatic macs, as displayed by root cells and end points (fig. 1J-K). Partition-based graph abstraction (PAGA), which estimates the connectivity of complex manifold partitions^53^, confirmed the reduction to two main trajectories, an inflammatory trajectory and a regulatory trajectory (fig. 1L). Signaling receptors among the top 100 dynamically expressed “driver” genes in the inflammatory trajectory included *Fgfr1*, *Il1r* and *Csf2rb*. Additionally, *C5ar1* and *Notch2* were dynamically expressed along the regulatory trajectory (fig. 1M). Both delineated trajectories, as well as the pseudotemporal ordering, remained broadly intact in *Irf8* deficiency (figs. 1J&L), hinting at compensatory mechanisms for the lack of Mc influx in the maintenance of downstream clusters. Interestingly, *Csf1r* was expressed at highest levels along the regulatory trajectory, while *Csf2rb* was dynamically expressed along the inflammatory trajectory (figs. S1.2G).

### Communication and spatial distribution of macrophage subsets

To confirm the RNA expression-based dermal mac subset composition utilizing protein expression profiles, we employed high-parametric multi-color flow cytometry. Dimensionality reduction and clustering based on a subset of markers (displayed in fig. S2A) confirmed the overall subset architecture, including the identification of Mc-like, inflammatory, regulatory, and homeostatic macs as well as sNaM (figs. 2A-B). *Irf8^-/-^* mice recapitulated the scRNAseq dataset marked by a lack of Mc-like macs, while the other subsets remained intact (figs. 2A-B). Substantiating a role for GM-CSF in homeostatic mac differentiation, *Csf2rb*^-/-^ mice had lower numbers of homeostatic macs, but increased numbers of inflammatory macs (figs. 2A-B). Moreover, pSTAT5 staining in homeostatic macs after GM-CSF restimulation *ex vivo* revealed a slight shift in pSTAT5 expression, more pronounced in inflammatory macs, indicative of active GM-CSF signal transmission during homeostasis (figs. 2C and S2C).

The CellChat algorithm^41^, a tool that predicts intercellular communication networks, identified 23 significant interaction networks among WT clusters. Informative examples include TNF and IL10 signaling, which originated from most clusters and were directed in large parts towards Mc-like macs, while regulatory macs were completely exempt from TNF-crosstalk. CCL-crosstalk was mostly directed towards clusters 2, 3, 6 and 8 (fig. 2D and S2D).

**Figure 2:**
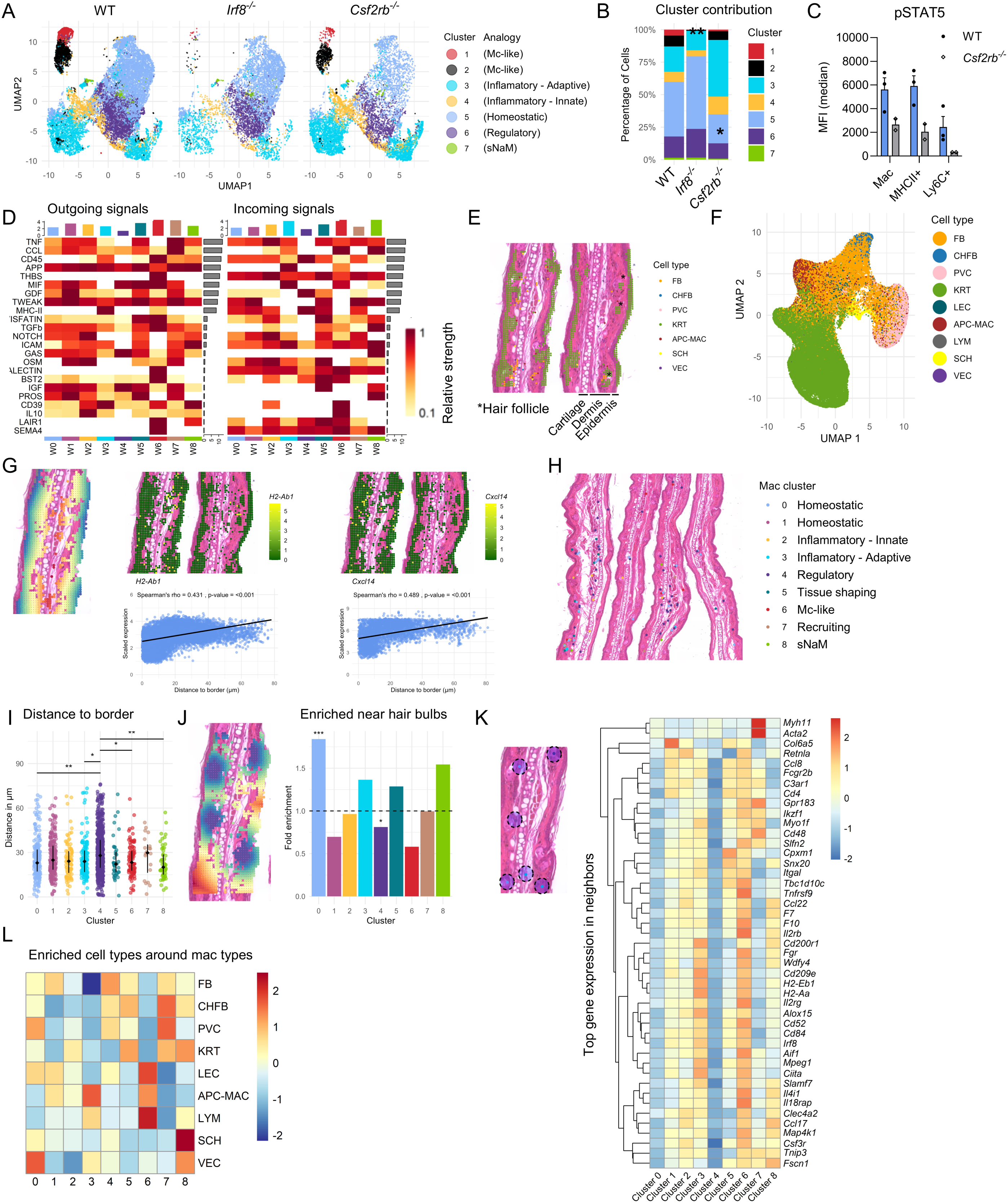
A: UMAP representation of spectral flow cytometry analysis of dermal macs of WT, *Irf8*^-/-^ and *Csf2rb*^-/-^. Cells from 3 mice are pooled per condition. Analogy to scRNAseq clusters displayed on the right. B: Cluster composition across the conditions described above. The asterisks indicate significantly reduced cluster sizes. C: pSTAT5 staining measured by spectral flow cytometry after *ex vivo* restimulation of dermal macs with 20 ng/ml GM-CSF for 20 min. D: Interactions between clusters estimated by the CellChat algorithm and ordered according to relative interaction strength. Outgoing signaling left and incoming signaling right, only WT is displayed, clustering as displayed in figure 1D. E: Cropped histology section of spatial RNAseq dataset with spots annotated by decomposition. Spotsize 8 µm. Abbreviations see below. F: UMAP plot colored according to main cell identities in all spots from 8 tissue sections from 4 separate ears. G: Relative distance of each spot to the tissue border (left) and scaled expression values for three exemplary genes with expression correlated to tissue depth. H: Cropped histology section displaying localization of mac clusters. I: Distance from tissue border calculated per mac cluster. J: Distance from hair bulb visualized on cropped histology section (blue:close, red:far), enrichment of mac clusters in direct vicinity (8µm) of hair bulbs.K: Scheme of neighbor analysis including a median of 8 spots around each mac spot (blue circle, left). Heatmap displaying top genes expressed in neighbor areas per cluster (right). L: Niche assay. Enrichment of cell types in the 20 neighbors of each spot assigned to a celltype (left) or in the 20 neighbors of each spot assigned to a mac cluster Statistics are in 2B: ANOVA pairwise comparison. 2I: Kruskal-Wallis followed by individual Dunn’s test. 2J: Fisher’s exact test with BH-correction. The cell types are abbreviated as: FB: Fibroblasts; CHFB: Chondrofibroblasts; PVC: Perivascular Cells; KRT: Keratinocytes; LEC: Lymphatic Endothelial Cells; APC-MAC: Antigen-Presenting Cells and mac; LYM: Lymphocytes; SCH: Schwann Cells; VEC: Vascular Endothelial Cells;

Next, we performed spatial RNA sequencing on the VisiumHD platform and annotated broad cell types by decomposition using a published scRNAseq dataset^34^ merged to our mac dataset. Choosing a spatial resolution of 8µm to recapitulate a roughly single-cell-level resolution, we obtained 315k spots on eight ear sections. Of those, 192k spots passed quality filtering and 115k were assigned to a cell type (figs. 2E-F &S2E). 7900 spots were identified as antigen presenting cells or mac (APC-MAC) as the most likely cell type. In a second decomposition step, we assigned the most likely mac cluster to each of the APC-MAC spots using our scRNAseq data of WT mac, with 3078 spots assigned to one of the mac clusters (fig. 2H and S2E). With a spatial resolution of 8x8 µm, each putative mac spot contained about 100-200 different gene transcripts. The limited sequencing depth makes it difficult to clearly separate the individual mac clusters in a dimensionality reduction diagram. Therefore, the assignments reflect only the most likely mac subgroup. We correlated gene expression to the distance from the epidermal border, and found that MHCII-associated genes and chemokines involved in mac recruitment (*Cxcl14* and *Ccl8,* not shown)^54,55^ were increasingly expressed in the deeper dermis (fig. 2G). Additionally, cluster 4 (regulatory macs) was found to be located in deeper tissue areas as compared to other clusters (fig. 2I). We manually identified all hair bulbs and found cluster 0 to be significantly enriched in the direct (8 µm) surrounding of hair bulbs (fig. 2J). Next, we defined areas around each mac spot including a mean of 6 spots in the direct surrounding. We identified a distinctive gene expression profile in the neighborhood of recruiting macs (cluster 7) with higher expression of marker genes of muscle cells (*Acta2*, *Myh11*). In the neighborhood of homeostatic macs (cluster 1), we found genes expressed in fibroblasts (*Col6a5*). *Ccl17* and *Ccl22* were highest in the neighborhood of Mc-like macs (cluster 6), while overall immune genes were enriched in the environment of innate-inflammatory and Mc-like macs (fig. 2K). When a broader neighborhood (20 spots) was assessed, we found enrichment of vascular endothelial cells near homeostatic macs (cluster 0), of fibroblasts near homeostatic macs (cluster 1), of perivascular cells near recruiting macs (cluster 7), and of lymphocytes around inflammatory-adaptive and mc-like macs (clusters 3 and 6). Furthermore, Schwann cells were highly enriched near sNaMs (cluster 8, fig. 2L). In summary, despite methodological limitations associated to limited sequencing depth, these associations provide an additional layer in the functional characterization of dermal mac subsets.

### Monocytes selectively replace a subset of dermal macrophages

As outlined before, infiltrating Mc and regulatory macs gave rise to resident macs with relatively low transcriptional activity, which may best be described as tonic base-line activaton. To explore turnover kinetics, we performed BM transplantation (BMTX) with focal ear shielding during irradiation, which protected resident macs (Res-mac) from irradiation damage while enabling complete BM chimerism. In contrast, conventional head shielding was associated with partial BM protection and incomplete chimerism (figs. 3A-B and S3A)^2,56^. Rapid engraftment in the Mc lineage within 5 days ensured that all incoming cells were marked by the CD45.1 congenic marker (figs. 3C and S3F). Ear-shielded mice did not show neutrophil influx into the skin, measurable loss of macs, or increased expression of *Bax* or inflammatory cytokines in dermal macs (figs. S3B-E). Analysis of dermal mac chimerism over time indicated continuous replacement by BM-derived macs (BM-mac, fig. 3D-E) and revealed that shielding protected ∼30% of dermal macs from exchange in addition to 14% of radioresistant macs (figs. 3F and S3F). Interestingly, there was almost no additional exchange between 16 and 24 weeks after BMTX, suggesting a saturation in exchange dynamics (fig. 3F). Thus, a substantial proportion of dermal macs was self-sustained and not replaced by BM-mac in adulthood. Ly6C downregulation occurred in dependence of the exchange kinetic (fig. 3G).

**Figure 3:**
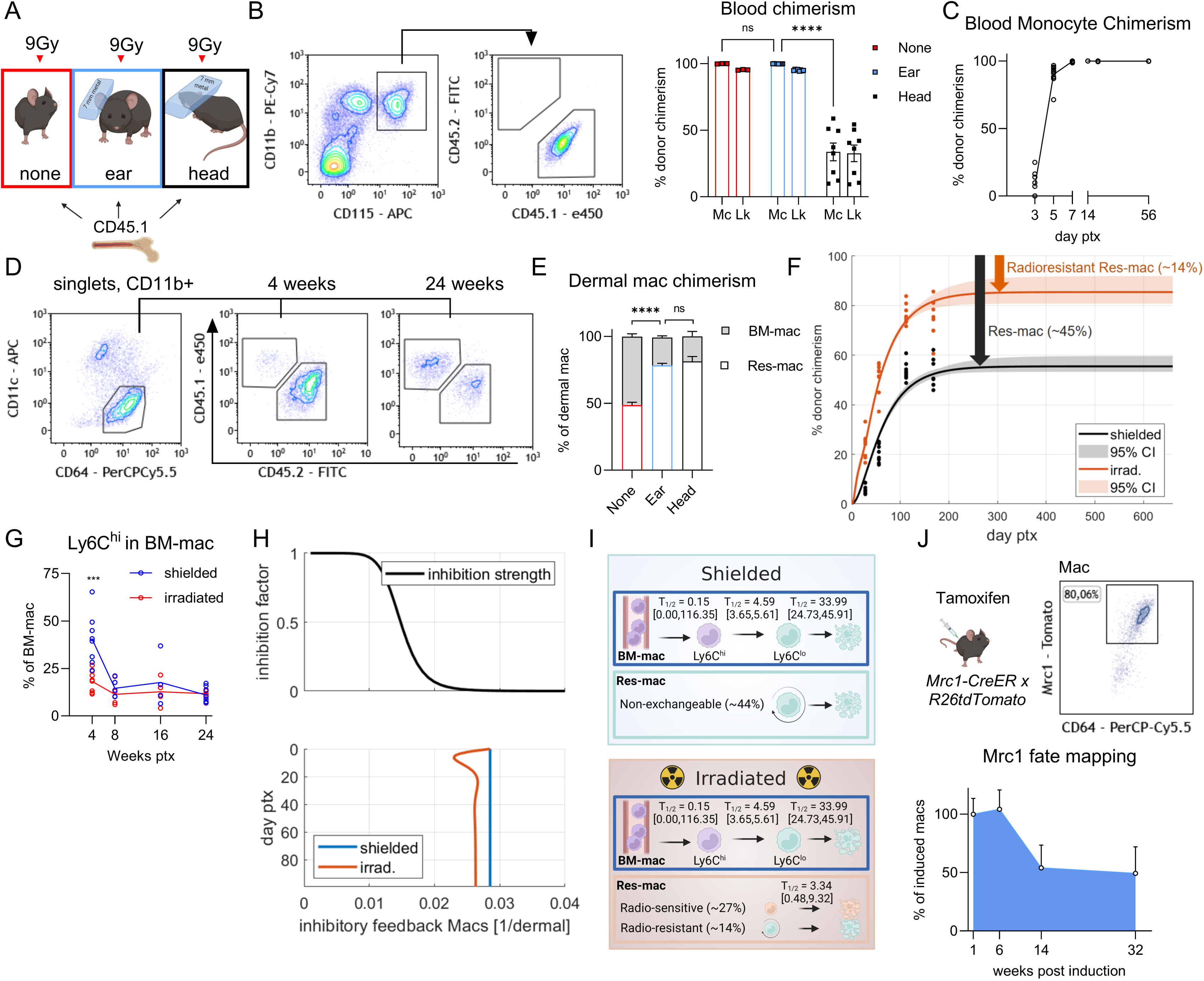
A: Scheme of different shielding approaches for bone marrow transplantation. B: Chimerism of blood Mc and blood leukocytes at 8 weeks after BMTX. Representative FACS plot for Mc on the left (shielded), quantification on the right. Mc = monocytes, Lk = leukocytes. N = 8-9 mice. C: Kinetic of Mc chimerism after BMTX with ear shielding. n= 8-11 mice. D, E: Flow cytometry analysis of dermal mac chimerism after BMTX with different shielding approaches or no shielding. D, Representative FACS plots for the ear shielding condition, E, quantification. n= 8-9 mice. F: Agreement between the kinetic model and experimentally measured dermal mac exchange (dots) for the shielded and irradiated condition and model prediction for the long-term exchange. The shaded regions indicate 95% CI of the model prediction. G: Ly6C expression of dermal macs after BMTX as percentage of BM-mac. n= 4-7 mice/timepoint. H: Estimated feedback strength for inhibiting mac recruitment into the dermis. Upper panel: Inhibition strength on recruitment on y axis. Lower panel: Time after BMTX on y axis. Both x-axes: proportion of the inhibitory resident mac pool. I: Kinetic modelling with estimated half-lives of tissue immigration, Ly6C downregulation and turnover rate of the exchangeable subset of dermal macs in the shielded (top) and irradiated (bottom) condition. J: Scheme and representative FACS plot of *Mrc1-CreER x R26tdTomato* mouse model (upper), quantification of dermal mac exchange in *Mrc1-CreER x R26tdTomato* mice (lower). n=4-5 mice per time point. Statistics: 3B, Two-way ANOVA; 3E, One-way ANOVA; 3G, student’s t-test including Holm-Šídák-correction for multiple comparisons.

Dynamic modelling of mac exchange kinetics using the Data2Dynamics toolbox^36^ supported the notion that ∼45% of dermal macs were exempt from renewal by BM-derived cells in the shielded dermis (figs. 3F and S3G). It further revealed the existence of a strong negative feedback mechanism that restricted immigration of abundant circulating precursors, yet enabled a rapid increase in mac number during emergency situations, such as loss of dermal macs by tissue irradiation, by reduction of the feedback strength (figs. 3H and S3H). Kinetic modelling suggested that Ly6C downregulation occurred with a half-life of ∼4.6 days after infiltration (CI 3.65-5.61 days), while the exchangeable mac compartment had a half-life of ∼34 days (CI 24.73-45.91 days, fig. 3I). This supported a simplified two-subset model, i.e. BM-mac, which were continuously replaced by circulating Mc, and non-exchangeable resident macs (Res-mac) (fig. 3I). Fate mapping using *Mrc1^CreERT2^*:*R26-tdTomato* mice^27^ confirmed an exemption of ∼40% of dermal macs from exchange by Mc-derived macs during 32 weeks (figs. 3J).

Taken together, we concluded that dermal macs are tightly maintained at specific densities via a negative feedback loop, and that a large subset of dermal macs is exempt from exchange by Mc.

### Transcription factor networks govern plasticity in resident macrophages

In line with a lack of circulating Mc, only 5% of dermal macs were exchanged within 8 weeks after transplantation of *Irf8^-/-^* bone marrow, as compared to 25% after transplantation of WT bone marrow (fig. S4A). Notably, blood Mc chimerism was near complete in either condition (fig. S4B). We wondered how WT Res-mac adapted to different levels of cellular influx by BM-mac. When transplanting *Irf8^-/-^* BM into WT mice, we found that WT Res-mac numerically adapted to the lack of incoming Mc, thus were able to self-sustain independently of BM-mac resupply, in line with constant total numbers of dermal macs in *Irf8^-/-^*mice (figs. 4A-B). Next, we compared Res-mac from WT mice transplanted with WT BM (WT Res-mac (WT BM)) with Res-mac from WT mice transplanted with Irf8-/- BM (WT Res-mac (*Irf8*^-/-^ BM)), i.e. which lacked supply by incoming Mc. For this purpose, we sorted BM-mac and Res-mac from WT mice transplanted with WT BM as well as WT Res-mac from mice transplanted with *Irf8*^-/-^ BM and performed bulk RNA sequencing. Thus, only cells from WT mice are included in this analysis. We found significant differences between the transcriptional profiles of BM-mac and Res-mac (figs. 4C and S4C&E). In BM-mac, a majority of the significantly upregulated 466 genes, as well as many of the enriched GO terms, were related to inflammation and immunity (fig. 4C). When aggregated expression counts of all upregulated genes in BM-mac or Res-mac were mapped to the steady-state scRNAseq data set, we found that BM-mac contributed mainly to Mc-like and inflammatory-adaptive macs, while Res-mac resembled regulatory and homeostatic macs (fig. 4D). Strikingly, a large set of genes potentially associated to differentiation, proliferation or density adaptation, in addition to some inflammatory genes, were upregulated in Res-mac (*Irf8*^-/-^ BM) compared to the more physiological Res-mac (WT BM). This revealed a profound plasticity in Res-mac to adapt to the environment and compensate for the absence of sufficient BM-mac (fig. 4E-F and S4D). Seeding BMDMs at different densities in 3D hydrogel cultures confirmed, in an exemplary manner, that expression of *Socs3* and *Irf4*, two of the upregulated genes in WT Res-mac (*Irf8*^-/-^ BM), was inversely associated with cellular density (fig. 4G), potentially indicating similar, robust mechanisms involved in density control *in* and *ex vivo*.

**Figure 4:**
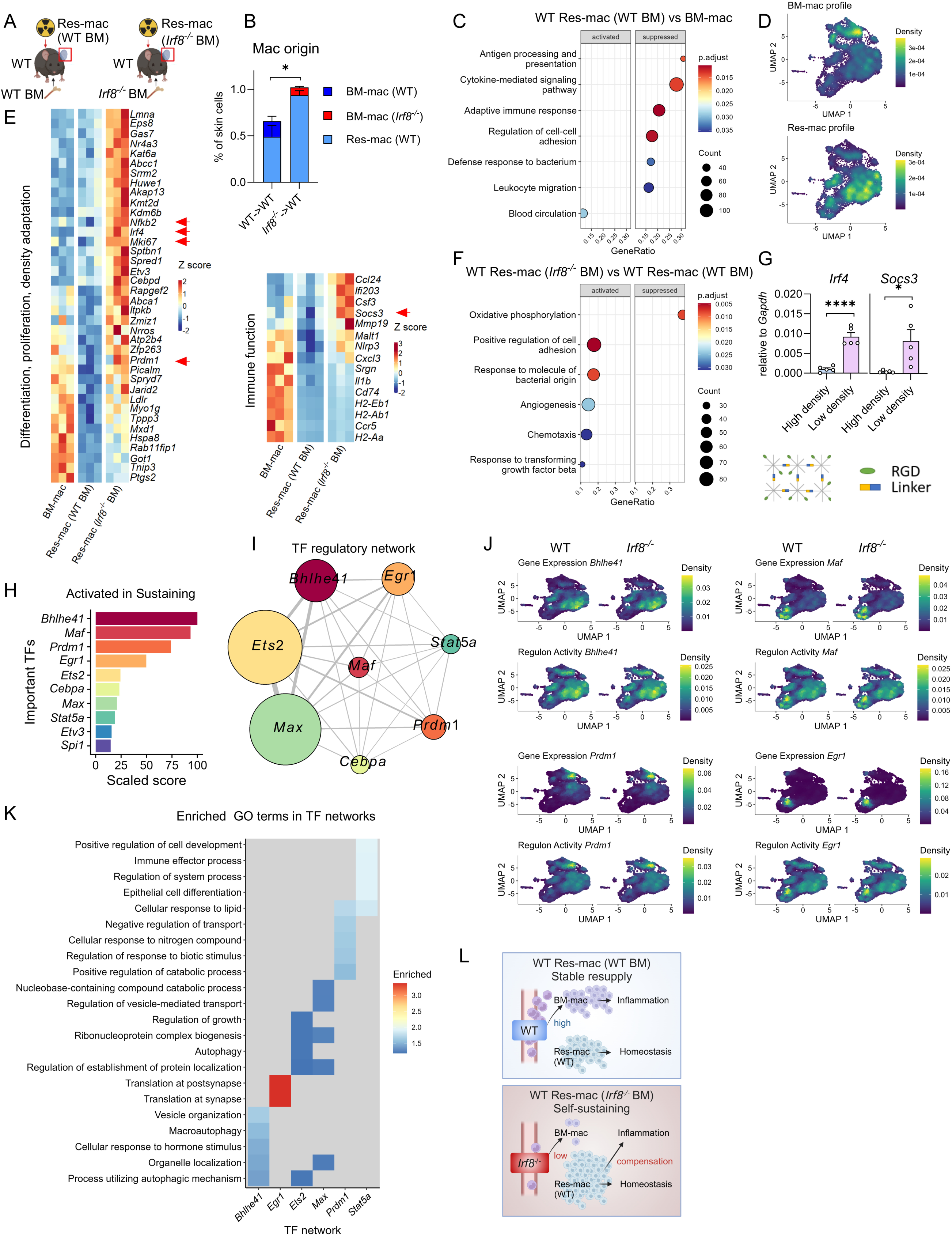
A: Scheme conceptualizing the two WT Res-mac conditions with different BM input. Analysis after 8 weeks. B: Mac origin in different BMTX conditions 8 weeks after BMTX by flow cytometry. Significance indicated for Res-mac densities. n=4-5 mice. WT BM-mac, WT Res-mac (WT BM), and WT Res-mac (*Irf8*^-/-^ BM) are used for bulkRNAseq. C: GSEA comparing GO terms upregulated or downregulated in WT Res-mac (WT BM) compared to WT BM-mac. Count: Number of genes present in gene set. GeneRatio: Ratio of overlapping genes to total genes in ranked input list. D: Aggregated gene score of all genes upregulated in BM-mac (above) or Res-mac (below) applied to the WT scRNAseq data set. E: Heatmaps displaying scaled expression of exemplary genes that are significantly differentially regulated in WT Res-mac (WT BM), and WT Res-mac (*Irf8*^-/-^ BM), related to differentiation, proliferation or density control (left) or to immune functions (right). F: GSEA comparing GO terms upregulated or downregulated in WT Res-mac (*Irf8*^-/-^ BM) compared to WT Res-mac (WT BM). G: Scheme of a PEG hydrogel culture (below) and qPCR results of BMDM seeded into hydrogel culture at high density (2000 cells/µl) or low density (660 cells/µl) after 5 days in culture. n=5 mice, 2 independent experiments. H: Top 10 transcription factor regulons upregulated in the “Sustaining” condition analyzed by GRaNPA (Gene Regulatory Network Prediction Analysis). The x-axis represents the scaled score, indicating the relative regulatory influence of each TF regulon. I: Gene regulatory network of important TFs (generated by SCENIC and assessed by GRaNPA) and their target genes in the scRNAseq dataset (as in fig. 1D). Size of dot: Represents the number of target genes regulated by each TF (regulon size). Connecting line: Indicates the amount of overlap in downstream genes (regulons) between two respective TFs. Line thickness: A thicker line corresponds to a greater overlap of target genes. J: UMAP visualisation of the expression levels of the 3 top transcription factors (top) and their inferred regulon activity (bottom) in the scRNAseq dataset, as analysed using SCENIC and GRaNPA (as in fig. 1D). K: Top enriched Gene Ontology (GO) terms for gene sets regulated by the top transcription factor regulons in the scRNAseq dataset (as in fig. 1D). This figure highlights the biological processes associated with each transcription factor regulon in the dataset. L: Scheme conceptualizing WT Res-mac compensation in the WT Res-mac (*Irf8*^-/-^ BM) condition compared to WT Res-mac (WT BM). Statistics: 4B, G, student’s t-test.

To gain a deeper mechanistic understanding of the observed gene expression patterns, we further investigated the transcription factors (TFs). We inferred a gene regulatory network from scRNA-seq using SCENIC^39^ and applied GRaNPA, a machine learning algorithm that identifies TFs within a network that predict a differential expression response between two conditions ^39^. For the DEG in the Res-mac comparison, the top 10 TFs, predictive of the adaptive density regulation, included *Bhlhe40*, *Maf*, and *Prdm1* (fig. 4H, S4F-G). The first two are known key differentiation regulators in macs^54,55^, while *Prdm1* has been described to control cell differentiation in cooperation with *Irf4*^56^. The top ten TFs where highly interconnected within the regulatory network (fig. 4I). *Maf* and *Prdm1* and the aggregated counts of the TF target genes were gradually expressed in the two root clusters of differentiation for and the *Bhlhe40* network in homeostatic macs (fig. 4J). Enriched GO terms within each gene set associated to any of the TFs included homeostatic functions involved in cell maintenance in addition to immune functions (fig. 4L).

Overall, BM-mac were transcriptionally primed towards immunity and Res-mac towards tissue homeostasis, but Res-mac were capable to compensate quantitatively and qualitatively for the lack of BM-mac through the coordinated activation of a TF network that was also engaged to maintain cellular homeostasis in steady state dermal macs (fig. 4L).

### Sequential scRNAseq unveils subset-specific macrophage replacement kinetics

In order to correlate Mc exchange with dermal mac subsets, we performed scRNAseq of macs at 4 and 16 weeks after bone marrow transplantation with focal ear shielding (fig. 5A). Of note, clustering was performed independently of the clustering in figure 1, to avoid artifacts in the steady-state data introduced by the irradiation. Strikingly, we found that all clusters received input by bone marrow-derived cells over time (fig. 5B-C). We used a two-state dynamic model to infer exchange kinetics by donor-derived cells for each cluster individually and to estimate the half-lives of the exchanges. Exchange rates were highly variable between different clusters, with only minimal BM-derived contribution to cluster T2 (with a half-life of >56 weeks), while cluster T4 was already largely exchanged after 4 weeks with a half-life of 2 weeks (fig. 5D). Core marker gene expressions, aggregated functional scores and gene set enrichment on GO terms recapitulated the heterogeneity among the different clusters (figs. 5E-F and S5A-C). Matching the data to steady-state scRNAseq, quadratic programming confirmed the close transcriptional similarity between the respective clusters of the two datasets (fig. S5D). Cluster T2 had the slowest turnover and lowest expression of immune-associated genes and thus resembled homeostatic macs, while cluster T4 resembled adaptive-inflammatory macs, and cluster T5/T6 regulatory macs. The correlation of distinctive transcriptional profiles (reflecting functional specialization) with characteristic turnover kinetics confirmed time dependency, where clusters with rapid turnover exhibited either an inflammatory or a regulatory profile, while long-term persistence in the tissue led to a more stable homeostatic phenotype with suppressed reactivity (fig. S5E). RNA velocity confirmed the previously described bidirectional resident mac maintenance, with differentiation starting from cluster T4 and even more from the clusters T5 and T6 (figs. 5G and S5F).

**Figure 5:**
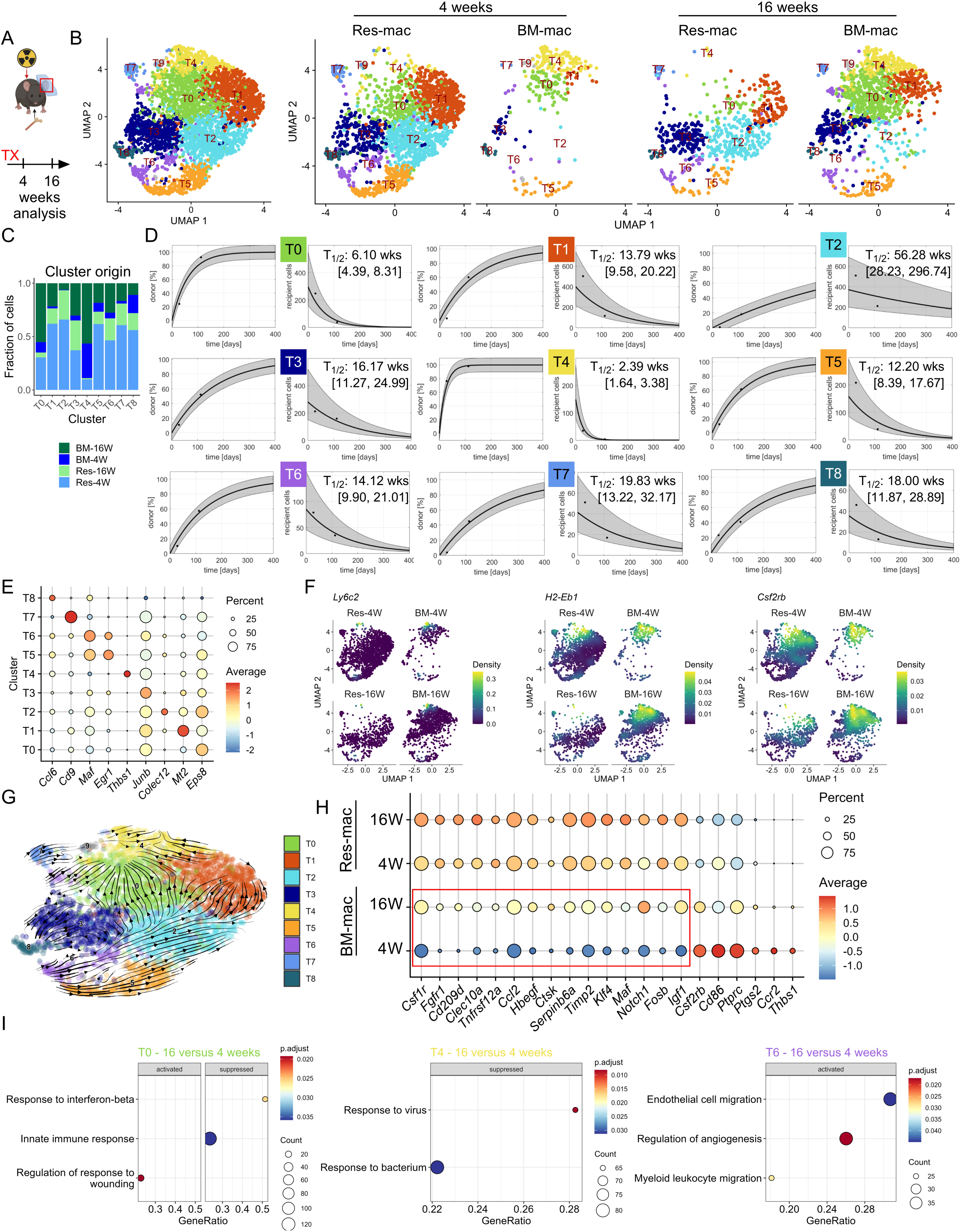
A: Scheme of experimental setup. B: UMAP representation based on gene expression profiling of sorted dermal macs highlighting the identified clusters using Seurat. Left, merged data, right, separated by experimental condition. Clusters have been obtained independently from clusters in figure 1D. C: Quantification of origin for each cluster. D: Estimated exchange kinetics of distinct clusters with the Data2Dynamics toolbox. For each indicated cluster, percentage of donor chimerism on the left, total number of recipient cells on the right. The calculated half-life is indicated in weeks, confidence interval in square brackets. E: Dot plot showing key marker genes differentially expressed in clusters 0-8. Color represents the average expression and dot size the fraction of cells expressing the gene. F: Key markers displayed as scaled expression on UMAP representation. G: RNA velocity displaying differentiation trajectories overlayed on UMAP representation with clustering. H: Dotplot displaying scaled expression per condition of selected DEGs focusing on adaptation between 4 to 16 weeks in BM-mac. I: Shortlisted enriched GO terms of clusters adapting between 4 and 16 weeks in the BM-mac conditions.

To identify tissue cues shaping BM-mac into specialized subsets, we focused on significant DEG in the total BM-mac population after 4 or 16 weeks of tissue adaptation. Of note, each of these populations contained a continuum of cells with tissue residency between 0 and 4 or 0 and 16 weeks, leading to underestimation of differences between the two conditions. The analysis revealed the upregulation of growth factor receptors *Csf1r* and *Fgfr1*, genes associated to tissue contact such as *Cd209d* and *Clec10a* as well as a set of TFs associated to a regulatory mac profile or density regulation (*Maf* and *Fosb*) at 16 weeks post transplantation. At the same time, immune-associated genes were downregulated (fig. 5H). GO-term based GSEA revealed upregulation of tissue remodeling terms and down-regulation of immune-associated terms in selected clusters (fig. 5I).

Taken together, distinct mac subsets were maintained at a stable density with individual exchange rates by Mc, directly translating into immune-related transcriptional profiles. As cells of different origin cluster together, we conclude that BM-mac retain the plastic potential to develop in all transcriptionally distinct, resident dermal mac subsets and that turnover characteristics depend on specific niche conditions rather than origin.

### BM-mac control mycobacteria in the dermis

Since *Irf8^-/-^* mice showed normal mac heterogeneity in homeostasis, and mounted reasonable immunity to staphylococcal skin infection (figs. S1.1G-H), we next wondered about the mechanistic basis of failing antimycobacterial immunity, which is the signature phenotypic trait in human *Irf8* deficiency. Accordingly, we dissected the dermal mac response after intradermal challenge with BCG. Tissue confocal microscopy revealed a high density of macs in WT mice after infection, with few macs harboring most of the bacteria, and high levels of iNOS expression. In contrast, in *Irf8^-/-^* mice, high numbers of neutrophils and debris covered the site of infection, while macs were sparse and iNOS expression was almost absent (figs. 6A and S6A). Flow cytometry confirmed a substantial increase in the dermal mac density during the peak of infection after one week in the WT, which was completely absent in *Irf8^-/-^*mice (figs. 6B-C and S6B). Although mycobacterial infection lowered the threshold for BM egress of Mc and their dedicated progenitors^28^, circulating Mc remained absent in infected *Irf8^-/-^* mice (fig. S6C). Instead, neutrophil numbers were increased in *Irf8^-/-^* mice (fig. 6D). Using fluorescent BCG, we identified a subset of macs that predominantly took up bacteria (fig. 6E). In *Irf8^-/-^*mice, neutrophils partially compensated for the lacking bacterial uptake by macs (fig. 6F), while *Irf8^-/-^* macs were fully able to phagocytose *in vitro* (fig. 6G).

**Figure 6:**
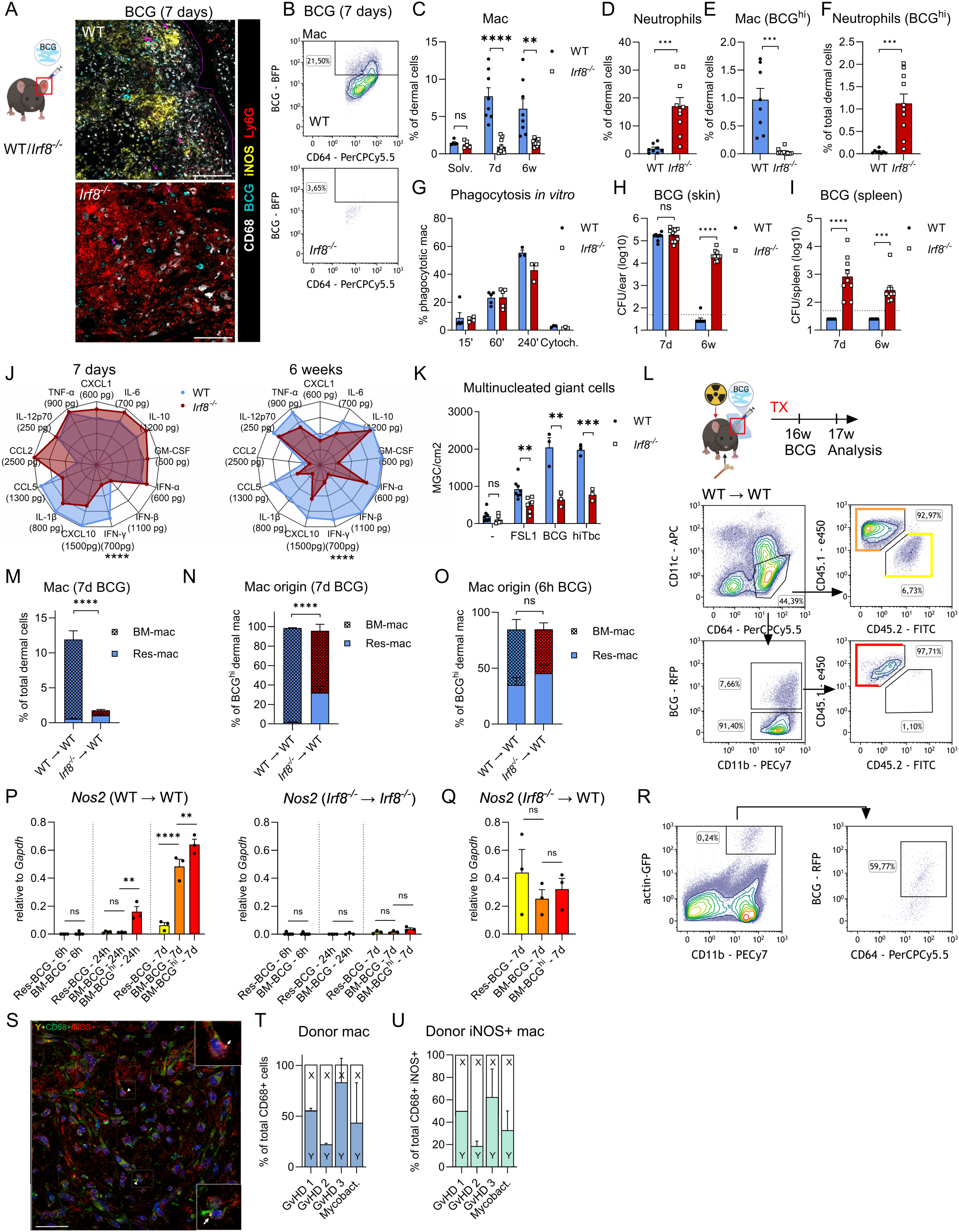
A: Scheme and confocal microscopy of cryosections of dermis 7d after intradermal BCG infection of WT and *Irf8*^-/-^ mice. Violet dashed line: Granuloma border. Violet arrows: Mac with high uptake of BCG. Representative image of 3 mice. White = CD68, light blue = BCG, yellow = iNOS, red = Ly6G. Scale bar = 100 µm. B/C/D: Representative flow cytometry plots (B) and quantification of dermal macs (C) or neutrophils (D) after 7d of intradermal BCG infection of WT and *Irf8*^-/-^ mice. n = 8-10 mice/infected condition. E/F: Subset of macs (E) or Neutrophils (F) that have taken up high numbers of BCG (BCG^hi^) determined by flow cytometry and fluorescent BCG. n=8 mice. G: Phagocytosis of fluorescent BCG by BMDM of WT or *Irf8^-/-^*-mice. n= 3 independent experiments. Cytoch: Cytochalasin B. H, I: BCG CFU counts of lysed dermal tissue (H) or spleen (I) after intradermal BCG infection of WT and *Irf8*^-/-^ mice. The dashed line indicates the detection limit. Samples without any detection were set to half the detection limit for statistics. n = 8-10 mice/condition. J: Bead-based Immunoassay (LegendPlex™) results of dermal tissue 7 days and 6 weeks after intradermal BCG infection of WT and *Irf8*^-/-^ mice. WT: n = 5 mice. *Irf8*^-/-^: n = 9-10 mice. K: *In vitro* formation of multinucleated giant cells from BMDMs after stimulation with the TLR2 ligand FSL1 (20 ng/ml) or with live (MOI 20) or heat-inactivated (10 µg/ml) mycobacteria for 6 days. n = 3 independent experiments. L: Scheme of experimental setup (above) and representative FACS plots (below) of dermal macs 7 days after intradermal BCG infection (upper panels), displaying the subset with high BCG uptake (lower panels). M/N/O: Quantification of BM-mac/Res-mac as % of total dermal cells (M) and as % taking up high amounts of BCG 7 days (N) or 6 h (O) after intradermal BCG infection. n = 8 mice (7 days), n = 5-7 mice (6h). P/Q: Relative *Nos2* expression measured by qPCR of sorted dermal mac at different time points post infection, 16 weeks post transplantation with indicated genotypes. n = 3-6 mice/condition. R: Exemplary FACS plots of dermal mac 24 h after Mc transfer and intradermal BCG infection. Representative of two mice. S: Exemplary patient histology after hematopoietic stem cell transplantation of female patients and a male donor who developed inflammatory skin condition (GvHD or mycobacterial infection) indicating BM-derived mac (using staining of Y chromosome) producing iNOS (white arrows). Scale bar 50 µm. T: Percentages of BM-mac (male donor, Y) of total CD68+ cells from patient samples. U: Quantification of BM-mac (male donor, Y) of total CD68+iNOS+ cells from patient samples. Statistics: Students t-test with Holm-Šídák-correction, and for figs. 6P-Q by two-way ANOVA followed by Tukey’s multiple comparison test. Comparisons in fig. 6G were non-significant.

Infected *Irf8^-/-^* mice were unable to clear the infection within 6 weeks, a time point when infection was fully cleared in the WT (fig. 6H). Furthermore, *Irf8^-/-^* mice showed BCG dissemination to the spleen (fig. 6I). A bead-based immunoassay on tissue lysates revealed that the impaired mac response was associated with an altered cytokine profile one week after infection, most notably with reduced amounts of IFNγ. At the same time, CCL2 was three-fold increased in *Irf8* deficiency, reflecting frustrated tissue recruitment of Mc. The cytokine profile after 6 weeks in the WT was marked by high levels of IFNγ, and GM-CSF, which were both reduced in *Irf8^-/-^* mice (fig. 6J). In addition, IFNβ expression, which has been associated with wound healing in injured tissue^57^, was elevated in the WT. Multinucleated giant cell (MGC) development of BMDM is a key feature of host-mycobacteria interaction, where reprogramming of lipid metabolism transforms cells into a pathogen-permissive state via an iNOS-driven process^20,28^. While we were unable to identify MGCs *in vivo* in this infection model, we explored MGC forming capacity *in vitro* as a cell-intrinsic antimycobacterial property, and found it to be partially impaired in *Irf8^-/-^*macs (fig. 6K and S6D).

Intradermal infection after BMTX revealed that the increase in mac numbers resulted exclusively from recruited BM-mac, whereas the number of Res-mac remained stable (figs. 6L-M and S6E). Intriguingly, after 7 days, fluorescent BCG was almost exclusively taken up by BM-mac in the WT. However, when *Irf8^-/-^* BM was transplanted into WT recipients, more Res-mac took up bacteria, indicating another layer of functional adaptation of Res-mac (fig. 6N). Notably, very early in infection (i.e. 6h p.i.), Res-mac and BM-mac equally contributed to mycobacterial uptake (fig. 6O). Transcription of *Nos2* (nitric oxide synthase 2), a key response to mycobacteria^28^, occurred nearly exclusively in BM-mac in WT mice, while it remained almost absent in Res-mac and *Irf8^-/-^* BM-mac (fig. 6P). This distribution of labor was confirmed using *Mrc1^CreERT2^*:*R26-tdTomato* mice infected with BCG (fig. S6F). *Nos2* expression was induced earlier and at higher levels in the population which ingested BCG (fig. 6P). WT mice transplanted with *Irf8^-/-^* BM revealed two striking findings: First, Res-mac were, again, able to partly compensate for lack of Nos2 expression in *Irf8*^-/-^ BM-mac. Second, *Irf8*^-/-^ BM-mac were able to express *Nos2* in a WT environment, which is in line with a previously postulated IRF8-IFNγ-iNOS axis^58^ where WT-derived IFNγ induces *Nos2* in *Irf8*^-/-^ BM-mac (fig. 6Q). *Il1b* expression was higher in BM-mac both in WT and *Irf8^-/-^* mice (fig. S6G).

To corroborate that Mc have a potential to immediately differentiate into an antimycobacterial tissue mac subset, we performed adoptive intravenous Mc transfer (WT Mc into *Irf8*^-/-^ recipients). Subsequent analysis confirmed that Mc rapidly infiltrated the site of infection and ingested large amounts of bacteria (fig. 6R). Transferring these findings to humans, we obtained skin biopsies of patients who previously underwent sex-mismatched BMTX where BM-mac were identifiable via Y chromosome staining. Three patients underwent skin biopsies for graft-versus-host disease and one for a chronic mycobacterial infection^59^. We confirmed a mixed chimerism for dermal macs and identified that both BM-mac and Res-mac expressed iNOS (figs. 6S-U).

### Specialized BM-mac are recruited to the site of infection

We next performed bulk RNA sequencing of Res-mac, BM-mac and BM-mac with high BCG content (BCG^hi^ mac) sorted from WT mice 16 weeks after BMTX and 7 days after BCG infection along with uninfected BM-mac and Res-mac 16 weeks after BMTX (sorting strategy in fig. 6L). Principal component analysis surprisingly displayed only subtle transcriptional changes in Res-mac despite 7 days exposure to a highly inflammatory microenvironment (fig. 7A). In contrast, BM-mac displayed a substantially different gene expression profile during infection, with a switch in gene expression associated to defense response to bacteria, including downregulation of IFNs and upregulation of *Tnf*, *Tlr4* and *Nod2*. In addition, we observed substantially altered gene expression profiles associated to chemokine and cytokine signaling, in response to hypoxia, as well as a switch in cholesterol metabolism with upregulation of *Abcg1* and *Ldlr* (figs. 7B-C and S7A-B). The latter two are important metabolic adaptive processes in the context of mycobacterial infections and skin infections^28,60^. Accordingly, many genes putatively related to the antimycobacterial defense, e.g. *Nos2*, *Il12* and *Arg1*^61,62^ were most abundantly expressed in BM-mac (figs. 7D-E). Among the selective genes significantly upregulated in the Res-mac in BCG infection, we found *Enpp2* which is involved in tissue remodeling and wound healing^63,64^ (fig. 7D). These findings further established a clear distribution of labor in dermal macs in the context of mycobacterial infections: a large fraction of urgently recruited BM-mac mount a strong inflammatory response, whereas Res-mac prevent systemic spread and tune up a selected set of genes guiding tissue remodeling from early on in infection.

**Figure 7:**
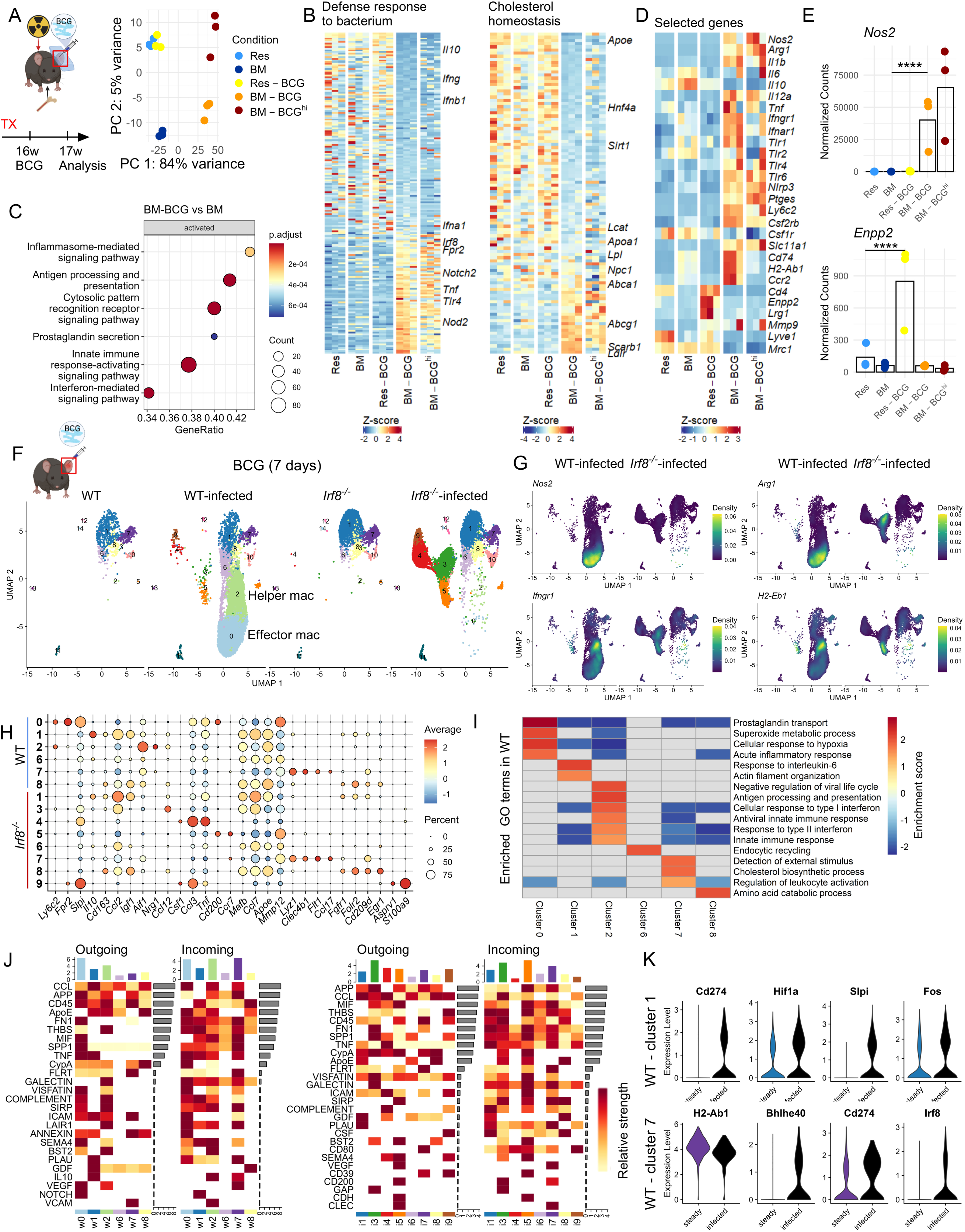
A: Scheme and PCA plot of bulkRNAseq of indicated cell types and conditions after 16 weeks of BMTX and 7 days after intradermal BCG infection. n = 3 mice/condition. Res: Res-mac; BM: BM-mac. B: Heatmaps displaying genes associated to two significantly regulated GO terms. C: Shortlisted enriched GO terms upregulated in BM-mac in BCG infection versus BM-mac in steady state. Count: Number of genes present in gene set. GeneRatio: Ratio of overlapping genes to total genes in ranked input list. D: Heatmaps displaying exemplary genes shortlisted based on presumed relevance in mycobacterial infections. E: Normalized counts of selected genes in bulkRNAseq. Significance determined in DEseq2 for indicated comparisons. F: Scheme and UMAP representation of scRNAseq analysis of sorted dermal macs from WT and *Irf8^-/-^*-mice with or without infection. n = 3 mice per condition. G: Scaled expression of key genes on UMAP representation of infected conditions. H: Dotplot displaying scaled marker gene expression of selected clusters across infected conditions of WT and *Irf8^-^*^/-^ mice. I: Shortlisted enriched GO terms in the main mac clusters in WT after infection compared to all other clusters. J: Cellchat analysis, heatmap displaying most likely signal communications between mac clusters in infection. Data from WT mice. K: Scaled expression of selected DEGs in resident clusters in steady state and infection.

Next, we performed scRNAseq of 21,000 sorted dermal macs of WT and *Irf8^-/-^* mice after 7 days of BCG infection or in steady state. The clusters identified in the uninfected condition remained broadly intact with infection (fig. 7F). In addition, two large clusters were recruited in the WT, marked by high levels of *Ly6c2* and *Csf2rb* expression, identifying them as BM-mac. These expressed profoundly distinct profiles from the Res-mac (figs. 7F-H). Cluster 0, named effector-macs expressed high levels of *Nos2* and *Arg1*, mostly concomitantly in the same cell (fig. S7C), along with *Fpr2* and other antimycobacterial effector components. Cluster 2, named helper-macs expressed high levels of *Ifngr1*, MHCII-associated genes, along with *Nrg1* and activation marker *Aif1* (figs. 7F-H). Accordingly, GSEA revealed enrichment of GO terms associated with acute inflammatory response, along with fatty acid metabolism and response to hypoxia in cluster 0 and reactivity to interferons and activation of antigen presentation in cluster 2 (fig. 7I). In *Irf8^-/-^*mice, several clusters (3, 4, 5, 9), which newly appeared in infection, expressed an atypical profile of reduced inflammatory markers, but high levels of genes associated with leukocyte recruitment such as *Ccl2*, *Ccl4*, but also *Csf1* and *Tnf* (figs. 7F-H).

Cellchat algorithm revealed globally expressed *Ccl* genes controlling mostly recruited macs. On the other hand, several pathways were directed from the BM-mac clusters 0 and 2 and acted on most clusters, such as fibronectin (*Fn1*), thrombospondin (*Thbs*), macrophage migration inhibitory factor (*Mif*), *Tnf* and galectin. In *Irf8^-/-^*dermal macs, inter-cluster communication was similarly diverse (fig. 7J, right) with higher expression of *Ccl* genes (fig. 7H). Steady-state macs remained largely intact after 7 days of infection (clusters 1, 7, 8, 10) and differential cluster-wise gene expression between steady-state and infection revealed only few changes, consistent with our previous findings of largely unaffected Res-mac transcriptomes during infection. *Hif1a* and *Slpi* were among the few DEGs in cluster 1 and 7. Furthermore downregulation of MHCII-associated genes and expression of *Cd274* was in line with immunosuppressive functions^65^, while upregulation of transcription factors *Fos* and *Bhlhe40*, reflected adoption of altered mac differentiation induced by an inflammatory environment.

Taken together, mechanisms that compensate for mac heterogeneity in *Irf8* deficiency during homeostasis failed to enable the dermis in coping with mycobacterial infections. Different subsets of recruited mac collaborated to mount an efficient defense, while atypically programmed recruited cells mainly expressed genes aiming at additional leukocyte recruitment in *Irf8* deficiency. A combination of impaired IFNγ action on mac and failed Mc recruitment may thus underlie the clinical presentation in *Irf8* deficiency.

## Discussion

In the skin, as in other barrier tissues, cells derived postnatally from definitive hematopoiesis are established to rapidly replace macs of primitive origin^12,66,67^. However, mechanisms regulating mac renewal, e.g by incoming Mc differentiating at site or by selfrenewal, remained largely elusive when we embarked on this current project. Here, we substantially expand on the complexity of the dermal mac network, which ensures high flexibility for divergent needs inherent to a constantly challenged tissue. Interestingly, we show that the dermis does not require Mc input to exhibit a multifaceted repertoire of macs throughout life. The transcriptionally heterogeneous dermal mac population comprised at least nine discriminable subsets with distinct functionality and renewal patterns. In homeostasis, interrelated resident mac subsets with high individual plasticity preserve density and function independently of Mc influx in all but one subset. It had already been suggested that a fraction of resident macs persists in the dermis for long time periods^68,69^. Yet our kinetic modelling adds precision to this understanding in that around 40% of dermal macs were not exchanged until late in adulthood, and that replacement was controlled by a tightly regulated negative feedback mechanism, preventing the constant influx of Mc. Interestingly, we found highly variable exchange kinetics between mac subsets, although all of them received input from bone marrow-derived progenitors to some extent. Moreover, using time-resolved scRNAseq after BMTX, we were able to explore how ontogeny and tissue residency define function and to identify potentially involved pathways. Our data indicated that functional diversification of dermal macs occurs largely independently of origin, but is substantially impacted by time of exposure to the microenvironment, i.e. by turnover kinetics.

Barrier tissues are challenged by constant exposure to the microbiota and other environmental stressors. Yet, regulatory mechanisms preventing permanent mac activation are incompletely understood^12,66^. Given the slow turnover kinetics of the homeostatic macs identified here (clusters 0 and 1), which account for approximately 35% of total dermal macs, it is tempting to speculate that they represent a differentiation endpoint resulting from converging trajectories of dermal macs with diverse fates. Furthermore, most dermal macs were polarized away from immunoreactivity, potentially explaining how the tissue restrains immune activation in light of a constant microbial challenge. In addition, upon bacterial challenge, these macs upregulate immunosuppressive genes and display a previously unrecognized plasticity to newly adopt immune reactivity. A homeostatic and long-term persisting mac population, tasked with tissue maintenance but not immune activation, has also been described in the intestine^7^, suggesting conserved mechanisms for mac maintenance and distribution of labor in barrier tissues. Our data expand previous reports showing that Mc-derived macs were more reactive towards pathogens and mount more robust inflammatory responses^9,70,71^ by exhibiting specialization within BM-mac subsets during infection. It seems important that “emergency” traits in macs are overridden by a tissue specific imprint only after long-term persistence in the organ, leading to the acquisition of another extreme phenotype as depicted in cluster T2 (fig. 6) or homeostatic macs (fig. 1). These long-lived macs show rather modest transcription of immunological genes and a half-life of more than 50 weeks. Accordingly, Mc-derived macs can contribute to, but are not required for, all facets of the functional mac mosaic through microanatomical adaptation over long time periods. Thus we expand the established concept that the tissue environment directs mac identity^72^.

Mycobacterial skin infections represent a localized, but chronic infectious stress that allowed us to study the inflammatory response of individual mac subsets. The hallmark infection in patients with *Irf8* deficiency is systemic BCGitis after vaccination^21^, which was mimicked in *Irf8*-deficient mice. Thus, our data confirmed the indispensable role of recently immigrated macs in the uptake of and inflammatory response to low pathogenic mycobacteria, whereas Res-mac appeared overwhelmed within a short time post infection. Anti-mycobacterial immunity is centered on the granuloma, a complex structure with temporal dynamics involving constant death of macs and recruitment of additional innate and adaptive immune cells, for instance of Mc^25,73^, but also profound tissue remodeling, driven by macs and non-immune cells^74,75^. In agreement with a previous study^76^, a lack of replenishing macs promoted infection via increased necrosis. Our results suggest that different mac subsets contribute to inflammation (BM-mac) and tissue remodeling (Res-mac) from early on in BCG infection, with cell-intrinsic programs and functions dependent on prior tissue exposure time.

It seems important to briefly address the methodological difficulties in the functional characterization of mac subsets *in vivo* even when the latest multidimensional technologies are used. High resolution spatial sequencing may serve as a particular example for this. The necessary methodological compromise between the spatial resolution closest to single-cell level and sufficient sequencing depth leads to the challenge that putative single mac spots (8x8 µm) contained only about 100-200 different gene transcripts each. As a consequence of this limited sequencing depth the separation of individual mac clusters in a dimensionality reduction diagram is impaired. Therefore, the assignment by spatial sequencing reflects only the most likely mac subgroup. However, we are confident that integration of multidimensional data with local and systemic perturbations of mac turnover and activation enables us to propose a well-reasoned model of a functional dermal mac mosaic comprising at least nine dermal mac subsets, with each subset receiving a specific cellular input.

Mc dominate in stress situations, while resident mac display plasticity to partially adapt to changing requirements. We propose a model in which tissue macs are not subdivided based on their origin, but where all subsets receive input from self-renewal of a priori resident and from recently incoming macs. Given the evolutionary importance of a functional mac network in barrier tissues, Mc replacement is largely redundant in dermal mac homeostasis, and mac subsets are mostly able to self-sustain and differentiate into other subsets. This allows for maximal temporal and spatial flexibility of dermal innate immunity in a tissue that constantly confronted with multiple stressors and seemingly conflicting tasks, like combatting infections^9,13,77^, tissue repair after trauma^78^ and maintaining sensitivity^79^.

## Acknowledgement

We thank Anita Imm, Reem Alsumati and the staff of the ZTZ “lighthouse” core facility for their excellent technical assistance.

Funding was given

| To | By | Project number |
| --- | --- | --- |
| Lighthouse<br>Core<br>Facility | Medical Faculty, University of Freiburg | 2021/A2-Fol; 2021/B3-Fol |
|  | DFG | 450392965 |
| AL&FL | IMM-PACT stipend (DFG) | 413517907 |
| FL | Spemann Graduate School for Biology and<br>Medicine (Ministry for Science, Research<br>and Arts of the State of Baden-Wuerttemberg) |  |
| RS | IMMediate stipend (DFG) |  |
|  | BMBF | 01EO2103 |
|  | Else Kröner Fresenius/Fritz Thyssen<br>Foundation, DFG | SFB-1479 – Project ID: 441891347 |
| JK | EQUIP (Medical Faculty, University of<br>Freiburg) |  |
|  | DFG | DFG 491676693 TRR 359 – PILOT |
| BB | ERC (European Union's Horizon 2020 research and innovation program) | 882424 |
|  | Swiss National Science Foundation | 733 310030_170320, 310030_188450, CRSII5_183478 |
| MP | Sobek Foundation, Ernst-Jung Foundation, Klaus Faber Foundation, Novo Nordisk Prize |  |
|  | DFG | CRC/TRR167 ProjectID259373024 “NeuroMac”, SFB 992 Project ID192904750, SFB 1160, SFB 1479 Project ID441891347, TRR 359 ProjectID491676693, Gottfried Wilhelm Leibniz Prize |
|  | DFG (Germany's Excellence Strategy) | CIBSS – EXC-2189 – Project ID390939984 |
| Sa | DFG | 491676693 TRR 359 - PILOT |
| PH | DFG | 283781347, 254895677, 446317895-SPP EXIT, ID 259373024 - TRR 167 NeuroMac, and 491676693 TRR 359 - PILOT |

## Author contributions

PH, Sa and FL conceptualized the study and wrote the manuscript.

FL performed the majority of experiments under the supervision of PH and the analysis of single-cell, bulk and spatial RNA sequencing data under the supervision of Sa.

RS performed the wet-lab spatial RNA sequencing experiment and helped with the analysis under the supervision of MP.

RS and OS (under the supervision of MP) and Sa performed parts of the wet-lab scRNAseq experiments.

SS performed the experiments involving hydrogels, bead immunoassays and substantially helped with experiments. This work is part of his MD thesis.

JN performed microscopy on tissue sections, created fluorescent BCG and helped with experiments.

EV performed the SCENIC and GRaNPA analysis under the supervision of JZ.

FW and ET performed the spectral flow cytometry under the supervision of BB.

JK performed the experiments involving head shielding and helped with experiments.

AL helped with experiments and provided relevant intellectual input.

AH, CD and NG helped with experiments.

EK helped with the analysis of the scRNA-seq data.

CK performed the modeling involving the D2D tool.

KM performed the microscopy on human tissue under the supervision of KE.

RZ and AM provided patient tissue samples.

All Coauthors have read and approved the final version of the manuscript.

## Figure legends

<u>Figure S1.1</u>

A: Representative flow cytometry plots and gating strategy for whole blood cells after red blood cell lysis.

B: Representative flow cytometry plots and gating strategy for total dermal cells after tissue digestion. *Irf8^-/^*^-^ mac expression of MHCII and LY6C on the right, gate coloring as in fig. 1C.

C: Representative flow cytometry plots and gating strategy for bone marrow.

D: Flow cytometry quantification of bone marrow Mc progenitors in WT and *Irf8^-/-^*mice. n = 3 mice for LSK cells and 8-9 mice for the other populations. LSK: hematopoietic stem cell; MDP: monocyte-dendritic cell progenitor; cMoP: common monocyte progenitor; BMc: Bone marrow monocyte.

E: CCR2 expression in bone marrow Mc progenitors in WT and *Irf8^-/-^*mice. Exemplary flow cytometry plot on the right. n= 6 mice.

F: Mac subsets according to CX_3_CR1 expression in dermal macs assessed in *Cx3cr1*^+/gfp^ reporter mice or *Cx3cr1* ^+/gfp^ reporter mice crossed to the *Irf8^-/-^* background. n = 5-6 mice.

G: Flow cytometry quantification of mac density during intradermal *S. aureus* infection. n = 6-7 mice per infected condition.

H: Ly6C expression in dermal macs 5 d after *S. aureus* infection. n = 6-7 mice.

I: CFU counts of lyzed tissue during intradermal *S. aureus* infection. n = 3-7 mice per condition and timepoint.

J: BrdU incorporation into dermal macs after 48 h of *S. aureus* infection or uninfected skin of the same mouse.

All statistics are calculated by students t-test including Holm-Šídák-correction for multiple t-tests in one graph.

<u>Figure S1.2</u>

A: UMAP representation showing the cluster distribution as in figure 1D separated per replicate.

B: Dot plot showing key marker genes differentially expressed in clusters 0-8 in *Irf8^-/-^*. Color represents the mean expression of the gene (“Average”) in the respective cluster and dot size (“Percent”) represents the fraction of cells in the cluster expressing the gene.

C: Density plots displaying scaled expression levels of exemplary genes on UMAP representation. Blue:low, yellow:high.

D: Shortlisted GO terms enriched in the WT clusters compared to all other clusters. Color displays normalized enrichment score.

E: Violinplot displaying scaled *Mki67* expression across clusters.

F: Violin plots showing module scores of gene sets associated with the indicated function. Wilcoxon test compares between genotypes. Box plots depict median with IQR. The gene sets are listed in supplemental table S9.

G: UMAP representation showing scaled *Csf1r* and *Csf2rb* expression (right plot) and velocity (i.e. up- or downregulation, left plot).

<u>Figure S2</u>

A: Heatmap displaying normalized expression levels of the cluster-defining markers for individual replicates.

B: Dotplot displaying expression levels of the genes for the cluster defining markers used for Fig. 2A in the scRNAseq dataset.

C: Histoplot displaying pSTAT5 levels in MHCII+ dermal macs after 20 min ex vivo GM-CSF restimulation, each replicate is displayed with individual legend (related to fig. 2C).

D: Relative contribution of specific CCL-CCR interaction dyads to the overall CCL-CCR interaction network (fig. 2D).

E: Visualized flow diagram of decomposition and analysis algorithm of the spatial dataset.

<u>Figure S3</u>

A: Photographs displaying the different shielding approaches.

B: Representative FACS plots and gating strategy for blood after bone marrow transplantation.

C, D: Neutrophil (C, Np) and mac (D) density 24 h after irradiation, quantified by FACS. n = 6-7 animals/condition.

E: Relative gene expression values in sorted dermal macs 24 h after irradiation. n = 6-8 animals/condition.

F: Schematic, representative FACS plots and gating strategy for skin after bone marrow transplantation.

G: Complete overview of the data used for kinetic modeling. The solid lines show the fitted dynamics of the kinetic model. The bands show the fitted error model that quantifies the uncertainty of individual data points.

H: Effect of the postulated inhibitory feedback on immigration of circulating Mc. Top row: Dynamics without feedback, Middle row: The most likely feedback strength on incoming Mc as estimated from our data. Bottom row: complete suppression of immigration that would be seen for the maximal feedback strength. In all three scenarios, BM-mac do not reach 100% due to Res-mac.

Statistics in S4C-E are calculated by one-way ANOVA followed by Tukey’s multiple comparison test. Where no statistic is indicated, comparisons were non-significant.

<u>Figure S4</u>

A/B: Dermal macs (A) and Mc (B) chimerism of *Irf8*^-/-^ versus WT transplantation conditions 8 weeks after transplantation.

C/D: Volcano plots of differentially expressed genes between WT Res-mac (WT BM) and WT BM-mac (C) and of WT Res-mac (*Irf8*^-/-^ BM) versus WT Res-mac (WT BM) (D). Cutoffs for logFC are set to 0.5 and for p-value to 0.05. Dots beyond these cutoffs are highlighted in red.

E: Heatmap displaying scaled gene expression levels of WT BM-mac, WT Res-mac (WT BM) and WT Res-mac (*Irf8*^-/-^ BM) of genes associated to the indicted GO term that had significantly different expression levels between BM-mac and Res-mac (WT BM).

F: Output of GRaNPA showing the density distribution of R2R^2R2 values from three random forest runs for the WT mac GRN predicting differential expression in the empty versus full niche. This highlights the predictive power of the GRN for gene expression changes under varying niche states.

G: Output of GRaNPA showing true versus predicted log2 fold-changes for the mac expression in the sustaining vs resupply condition. Predictions are derived from the WT mac SCENIC-GRN. The plot reflects the performance of GRaNPA in predicting gene expression changes between conditions from the GRN.

Statistics: S4A,B, one-way-ANOVA.

<u>Figure S5</u>

A: Shortlisted GO terms enriched in the respective clusters compared to all other clusters. Same terms as in figure S1.2C

B: Violin plots showing module scores of gene sets associated with the indicated function. Wilcoxon test compares relevant clusters. Box plots depict median with IQR. The gene sets are listed in table S9.

C: Density plots displaying scaled expression levels of exemplary genes.

D: Violin plots showing quadratic programming weight of clusters 1,3,4,6,7 and 8 of steady state scRNAseq data calculated for clusters T0-T8 of BMTX scRNAseq data to display similarity between the respective clusters.

E: Closest correlation between cluster identities with steady-state cluster identities based on marker gene expression and quadratic programming (fig. S5.1A)

F: RNA velocities derived from the dynamical model projected into the pseudotime UMAP representation (as in fig. 5G) separated according to condition.

<u>Figure S6</u>

A: Single stainings correlating to fig. 6A

B: FACS plots and gating strategy of skin 7 days post intradermal BCG infection.

C: Blood myeloid cell counts 7 days after intradermal BCG infection of WT and *Irf8*^-/-^ mice. n=5 mice. Np: Neutrophils

D: Brightfield microscopy pictures correlating to fig. 6K after Hemacolor® staining of adherent cells. Violet arrow: Exemplary MGC.

E: Representative FACS plots for the *Irf8*^-/-^ -> WT condition, correlating to fig. 6L

F: Relative *Nos2* expression measured by qPCR in sorted macs after 7 days of BCG infection in *Mrc1-CreERT2* reporter mice. n = 4 mice/condition

G: Relative *Il1b* expression measured by qPCR of sorted dermal macs at different time points post infection, 16 weeks post transplantation. n = 3-6 mice/condition.

H: Single colors of the exemplary histology image displayed in fig. 6S.

<u>Figure S7</u>

A: Volcano plots of different comparisons from bulkRNAseq after 16 weeks of BMTX and 7 days after intradermal BCG infection. Cutoffs for logFC are set to 0.5 and for p-value to 0.05. Dots beyond these cutoffs are highlighted in red.

B: Heatmaps displaying normalized RNA expression of selected GO terms significantly enriched in BM-mac after infection.

C: Concomitant expression of *Nos2* and *Arg1* in recruited macs in cluster 0, WT.

D: Shortlisted enriched GO terms in the main mac clusters in *Irf8^-/-^* after 7 days infection compared to all other clusters.

## References

1. Tamoutounour, S., Guilliams, M., Montanana Sanchis, F., Liu, H., Terhorst, D., Malosse, C., Pollet, E., Ardouin, L., Luche, H., Sanchez, C., et al. (2013). Origins and Functional Specialization of Macrophages and of Conventional and Monocyte-Derived Dendritic Cells in Mouse Skin. Immunity 39, 925–938. 10.1016/j.immuni.2013.10.004.

2. Kolter, J., Feuerstein, R., Zeis, P., Hagemeyer, N., Paterson, N., d’Errico, P., Baasch, S., Amann, L., Masuda, T., Lösslein, A., et al. (2019). A Subset of Skin Macrophages Contributes to the Surveillance and Regeneration of Local Nerves. Immunity 50, 1482–1497.e7. 10.1016/j.immuni.2019.05.009.

3. Schulz, C., Gomez Perdiguero, E., Chorro, L., Szabo-Rogers, H., Cagnard, N., Kierdorf, K., Prinz, M., Wu, B., Jacobsen, S.E.W., Pollard, J.W., et al. (2012). A lineage of myeloid cells independent of Myb and hematopoietic stem cells. Science 336, 86–90. 10.1126/science.1219179.

4. Gomez Perdiguero, E., Klapproth, K., Schulz, C., Busch, K., Azzoni, E., Crozet, L., Garner, H., Trouillet, C., de Bruijn, M.F., Geissmann, F., et al. (2015). Tissue-resident macrophages originate from yolk sac-derived erythro-myeloid progenitors. Nature 518, 547–551. 10.1038/nature13989.

5. Bain, C.C., Bravo-Blas, A., Scott, C.L., Perdiguero, E.G., Geissmann, F., Henri, S., Malissen, B., Osborne, L.C., Artis, D., and Mowat, A.M. (2014). Constant replenishment from circulating monocytes maintains the macrophage pool in the intestine of adult mice. Nat. Immunol. 15, 929–937. 10.1038/ni.2967.

6. Miyake, K., Ito, J., Takahashi, K., Nakabayashi, J., Brombacher, F., Shichino, S., Yoshikawa, S., Miyake, S., and Karasuyama, H. (2024). Single-cell transcriptomics identifies the differentiation trajectory from inflammatory monocytes to pro-resolving macrophages in a mouse skin allergy model. Nat. Commun. 15, 1666. 10.1038/s41467-024-46148-4.

7. De Schepper, S., Verheijden, S., Aguilera-Lizarraga, J., Viola, M.F., Boesmans, W., Stakenborg, N., Voytyuk, I., Schmidt, I., Boeckx, B., Dierckx de Casterlé, I., et al. (2018). Self-Maintaining Gut Macrophages Are Essential for Intestinal Homeostasis. Cell 175, 400–415.e13. 10.1016/j.cell.2018.07.048.

8. Shaw, T.N., Houston, S.A., Wemyss, K., Bridgeman, H.M., Barbera, T.A., Zangerle-Murray, T., Strangward, P., Ridley, A.J.L., Wang, P., Tamoutounour, S., et al. (2018). Tissue-resident macrophages in the intestine are long lived and defined by Tim-4 and CD4 expression. J. Exp. Med. 215, 1507–1518. 10.1084/jem.20180019.

9. Abtin, A., Jain, R., Mitchell, A.J., Roediger, B., Brzoska, A.J., Tikoo, S., Cheng, Q., Ng, L.G., Cavanagh, L.L., von Andrian, U.H., et al. (2014). Perivascular macrophages mediate neutrophil recruitment during bacterial skin infection. Nat. Immunol. 15, 45–53. 10.1038/ni.2769.

10. Barreiro, O., Cibrian, D., Clemente, C., Alvarez, D., Moreno, V., Valiente, Í., Bernad, A., Vestweber, D., Arroyo, A.G., Martín, P., et al. (2016). Pivotal role for skin transendothelial radio-resistant anti-inflammatory macrophages in tissue repair. eLife 5, e15251. 10.7554/eLife.15251.

11. Amit, I., Winter, D.R., and Jung, S. (2016). The role of the local environment and epigenetics in shaping macrophage identity and their effect on tissue homeostasis. Nat. Immunol. 17, 18–25. 10.1038/ni.3325.

12. Mowat, A.M., Scott, C.L., and Bain, C.C. (2017). Barrier-tissue macrophages: functional adaptation to environmental challenges. Nat. Med. 23, 1258–1270. 10.1038/nm.4430.

13. Feuerstein, R., Seidl, M., Prinz, M., and Henneke, P. (2015). MyD88 in Macrophages Is Critical for Abscess Resolution in Staphylococcal Skin Infection. J. Immunol. 194, 2735–2745. 10.4049/jimmunol.1402566.

14. Feuerstein, R., Forde, A.J., Lohrmann, F., Kolter, J., Ramirez, N.J., Zimmermann, J., Gomez de Agüero, M., and Henneke, P. (2020). Resident macrophages acquire innate immune memory in staphylococcal skin infection. eLife 9, e55602. 10.7554/eLife.55602.

15. Forde, A.J., Kolter, J., Zwicky, P., Baasch, S., Lohrmann, F., Eckert, M., Gres, V., Lagies, S., Gorka, O., Rambold, A.S., et al. (2023). Metabolic rewiring tunes dermal macrophages in staphylococcal skin infection. Sci. Immunol. 8, eadg3517. 10.1126/sciimmunol.adg3517.

16. Amorim, A., De Feo, D., Friebel, E., Ingelfinger, F., Anderfuhren, C.D., Krishnarajah, S., Andreadou, M., Welsh, C.A., Liu, Z., Ginhoux, F., et al. (2022). IFNγ and GM-CSF control complementary differentiation programs in the monocyte-to-phagocyte transition during neuroinflammation. Nat. Immunol. 23, 217–228. 10.1038/s41590-021-01117-7.

17. Dai, X.-M., Ryan, G.R., Hapel, A.J., Dominguez, M.G., Russell, R.G., Kapp, S., Sylvestre, V., and Stanley, E.R. (2002). Targeted disruption of the mouse colony-stimulating factor 1 receptor gene results in osteopetrosis, mononuclear phagocyte deficiency, increased primitive progenitor cell frequencies, and reproductive defects. Blood 99, 111–120. 10.1182/blood.v99.1.111.

18. Wiktor-Jedrzejczak, W., Ratajczak, M.Z., Ptasznik, A., Sell, K.W., Ahmed-Ansari, A., and Ostertag, W. (1992). CSF-1 deficiency in the op/op mouse has differential effects on macrophage populations and differentiation stages. Exp. Hematol. 20, 1004–1010.

19. Jiemy, W.F., van Sleen, Y., van der Geest, K.S., ten Berge, H.A., Abdulahad, W.H., Sandovici, M., Boots, A.M., Heeringa, P., and Brouwer, E. (2020). Distinct macrophage phenotypes skewed by local granulocyte macrophage colony-stimulating factor (GM-CSF) and macrophage colony-stimulating factor (M-CSF) are associated with tissue destruction and intimal hyperplasia in giant cell arteritis. Clin. Transl. Immunol. 9, e1164. 10.1002/cti2.1164.

20. Gharun, K., Senges, J., Seidl, M., Lösslein, A., Kolter, J., Lohrmann, F., Fliegauf, M., Elgizouli, M., Vavra, M., Schachtrup, K., et al. (2017). Mycobacteria exploit nitric oxide-induced transformation of macrophages into permissive giant cells. EMBO Rep. 18, 2144–2159. 10.15252/embr.201744121.

21. Hambleton, S., Salem, S., Bustamante, J., Bigley, V., Boisson-Dupuis, S., Azevedo, J., Fortin, A., Haniffa, M., Ceron-Gutierrez, L., Bacon, C.M., et al. (2011). IRF8 mutations and human dendritic-cell immunodeficiency. N. Engl. J. Med. 365, 127–138. 10.1056/NEJMoa1100066.

22. Kurotaki, D., Yamamoto, M., Nishiyama, A., Uno, K., Ban, T., Ichino, M., Sasaki, H., Matsunaga, S., Yoshinari, M., Ryo, A., et al. (2014). IRF8 inhibits C/EBPα activity to restrain mononuclear phagocyte progenitors from differentiating into neutrophils. Nat. Commun. 5. 10.1038/ncomms5978.

23. Kuntz, M., Kohlfürst, D.S., Feiterna-Sperling, C., Krüger, R., Baumann, U., Buchtala, L., Elling, R., Grote, V., Hübner, J., Hufnagel, M., et al. (2020). Risk Factors for Complicated Lymphadenitis Caused by Nontuberculous Mycobacteria in Children. Emerg. Infect. Dis. 26, 579–586. 10.3201/eid2603.191388.

24. Wu, U.-I., and Holland, S.M. (2015). Host susceptibility to non-tuberculous mycobacterial infections. Lancet Infect. Dis. 15, 968–980. 10.1016/S1473-3099(15)00089-4.

25. Pagán, A.J., and Ramakrishnan, L. (2018). The Formation and Function of Granulomas. Annu. Rev. Immunol. 36, 639–665. 10.1146/annurev-immunol-032712-100022.

26. Holtschke, T., Löhler, J., Kanno, Y., Fehr, T., Giese, N., Rosenbauer, F., Lou, J., Knobeloch, K.-P., Gabriele, L., Waring, J.F., et al. (1996). Immunodeficiency and chronic myelogenous leukemia-like syndrome in mice with a targeted mutation of the ICSBP gene. Cell 87, 307–317.

27. Masuda, T., Amann, L., Monaco, G., Sankowski, R., Staszewski, O., Krueger, M., Del Gaudio, F., He, L., Paterson, N., Nent, E., et al. (2022). Specification of CNS macrophage subsets occurs postnatally in defined niches. Nature 604, 740–748. 10.1038/s41586-022-04596-2.

28. Lösslein, A.K., Lohrmann, F., Scheuermann, L., Gharun, K., Neuber, J., Kolter, J., Forde, A.J., Kleimeyer, C., Poh, Y.Y., Mack, M., et al. (2021). Monocyte progenitors give rise to multinucleated giant cells. Nat. Commun. 12, 2027. 10.1038/s41467-021-22103-5.

29. Takaki, K., Davis, J.M., Winglee, K., and Ramakrishnan, L. (2013). Evaluation of the pathogenesis and treatment of Mycobacterium marinum infection in zebrafish. Nat. Protoc. 8, 1114–1124. 10.1038/nprot.2013.068.

30. Zhu, A., Ibrahim, J.G., and Love, M.I. (2019). Heavy-tailed prior distributions for sequence count data: removing the noise and preserving large differences. Bioinforma. Oxf. Engl. 35, 2084–2092. 10.1093/bioinformatics/bty895.

31. Herman, J.S., Sagar, and Grün, D. (2018). FateID infers cell fate bias in multipotent progenitors from single-cell RNA-seq data. Nat. Methods 15, 379–386. 10.1038/nmeth.4662.

32. Kueckelhaus, J., Frerich, S., Kada-Benotmane, J., Koupourtidou, C., Ninkovic, J., Dichgans, M., Beck, J., Schnell, O., and Heiland, D.H. (2024). Inferring histology-associated gene expression gradients in spatial transcriptomic studies. Nat. Commun. 15, 7280. 10.1038/s41467-024-50904-x.

33. Cable, D.M., Murray, E., Zou, L.S., Goeva, A., Macosko, E.Z., Chen, F., and Irizarry, R.A. (2022). Robust decomposition of cell type mixtures in spatial transcriptomics. Nat. Biotechnol. 40, 517–526. 10.1038/s41587-021-00830-w.

34. Anderson-Crannage, M., Ascensión, A.M., Ibanez-Solé, O., Zhu, H., Schaefer, E., Ottomanelli, D., Hochberg, B., Pan, J., Luo, W., Tian, M., et al. (2023). Inflammation-mediated fibroblast activation and immune dysregulation in collagen VII-deficient skin. Front. Immunol. 14, 1211505. 10.3389/fimmu.2023.1211505.

35. Lutolf, M.P., Lauer-Fields, J.L., Schmoekel, H.G., Metters, A.T., Weber, F.E., Fields, G.B., and Hubbell, J.A. (2003). Synthetic matrix metalloproteinase-sensitive hydrogels for the conduction of tissue regeneration: Engineering cell-invasion characteristics. Proc. Natl. Acad. Sci. 100, 5413–5418. 10.1073/pnas.0737381100.

36. Raue, A., Steiert, B., Schelker, M., Kreutz, C., Maiwald, T., Hass, H., Vanlier, J., Tönsing, C., Adlung, L., Engesser, R., et al. (2015). Data2Dynamics: a modeling environment tailored to parameter estimation in dynamical systems. Bioinformatics 31, 3558–3560. 10.1093/bioinformatics/btv405.

37. Raue, A., Schilling, M., Bachmann, J., Matteson, A., Schelke, M., Kaschek, D., Hug, S., Kreutz, C., Harms, B.D., Theis, F.J., et al. (2013). Lessons Learned from Quantitative Dynamical Modeling in Systems Biology. PLOS ONE 8, e74335. 10.1371/journal.pone.0074335.

38. Kreutz, C., Raue, A., Kaschek, D., and Timmer, J. (2013). Profile likelihood in systems biology. FEBS J. 280, 2564–2571. 10.1111/febs.12276.

39. Aibar, S., González-Blas, C.B., Moerman, T., Huynh-Thu, V.A., Imrichova, H., Hulselmans, G., Rambow, F., Marine, J.-C., Geurts, P., Aerts, J., et al. (2017). SCENIC: single-cell regulatory network inference and clustering. Nat. Methods 14, 1083–1086. 10.1038/nmeth.4463.

40. Kamal, A., Arnold, C., Claringbould, A., Moussa, R., Servaas, N.H., Kholmatov, M., Daga, N., Nogina, D., Mueller-Dott, S., Reyes-Palomares, A., et al. (2023). GRaNIE and GRaNPA: inference and evaluation of enhancer-mediated gene regulatory networks. Mol. Syst. Biol. 19, e11627. 10.15252/msb.202311627.

41. Jin, S., Guerrero-Juarez, C.F., Zhang, L., Chang, I., Ramos, R., Kuan, C.-H., Myung, P., Plikus, M.V., and Nie, Q. (2021). Inference and analysis of cell-cell communication using CellChat. Nat. Commun. 12, 1088. 10.1038/s41467-021-21246-9.

42. Wu, T., Hu, E., Xu, S., Chen, M., Guo, P., Dai, Z., Feng, T., Zhou, L., Tang, W., Zhan, L., et al. (2021). clusterProfiler 4.0: A universal enrichment tool for interpreting omics data. The Innovation 2, 100141. 10.1016/j.xinn.2021.100141.

43. Love, M.I., Huber, W., and Anders, S. (2014). Moderated estimation of fold change and dispersion for RNA-seq data with DESeq2. Genome Biol. 15, 550. 10.1186/s13059-014-0550-8.

44. Wolf, F.A., Angerer, P., and Theis, F.J. (2018). SCANPY: large-scale single-cell gene expression data analysis. Genome Biol. 19, 15. 10.1186/s13059-017-1382-0.

45. Hafemeister, C., and Satija, R. (2019). Normalization and variance stabilization of single-cell RNA-seq data using regularized negative binomial regression. Genome Biol. 20, 296. 10.1186/s13059-019-1874-1.

46. La Manno, G., Soldatov, R., Zeisel, A., Braun, E., Hochgerner, H., Petukhov, V., Lidschreiber, K., Kastriti, M.E., Lönnerberg, P., Furlan, A., et al. (2018). RNA velocity of single cells. Nature 560, 494–498. 10.1038/s41586-018-0414-6.

47. Hao, Y., Hao, S., Andersen-Nissen, E., Mauck, W.M., Zheng, S., Butler, A., Lee, M.J., Wilk, A.J., Darby, C., Zager, M., et al. (2021). Integrated analysis of multimodal single-cell data. Cell 184, 3573–3587.e29. 10.1016/j.cell.2021.04.048.

48. Grün, D. (2020). Revealing dynamics of gene expression variability in cell state space. Nat. Methods 17, 45–49. 10.1038/s41592-019-0632-3.

49. quadprog: Functions to solve Quadratic Programming Problems. | BibSonomy https://www.bibsonomy.org/publication/8b73f0cd8854d54f0d900b6f8d7a6059.

50. Gayoso, A., Weiler, P., Lotfollahi, M., Klein, D., Hong, J., Streets, A., Theis, F.J., and Yosef, N. (2024). Deep generative modeling of transcriptional dynamics for RNA velocity analysis in single cells. Nat. Methods 21, 50–59. 10.1038/s41592-023-01994-w.

51. Kierdorf, K., Erny, D., Goldmann, T., Sander, V., Schulz, C., Perdiguero, E.G., Wieghofer, P., Heinrich, A., Riemke, P., Hölscher, C., et al. (2013). Microglia emerge from erythromyeloid precursors via Pu.1- and Irf8-dependent pathways. Nat. Neurosci. 16, 273–280. 10.1038/nn.3318.

52. Hagemeyer, N., Kierdorf, K., Frenzel, K., Xue, J., Ringelhan, M., Abdullah, Z., Godin, I., Wieghofer, P., Costa Jordão, M.J., Ulas, T., et al. (2016). Transcriptome-based profiling of yolk sac-derived macrophages reveals a role for Irf8 in macrophage maturation. EMBO J. 35, 1730– 1744. 10.15252/embj.201693801.

53. Wolf, F.A., Hamey, F.K., Plass, M., Solana, J., Dahlin, J.S., Göttgens, B., Rajewsky, N., Simon, L., and Theis, F.J. (2019). PAGA: graph abstraction reconciles clustering with trajectory inference through a topology preserving map of single cells. Genome Biol. 20, 59. 10.1186/s13059-019-1663-x.

54. Gowhari Shabgah, A., Haleem Al-qaim, Z., Markov, A., Valerievich Yumashev, A., Ezzatifar, F., Ahmadi, M., Mohammad Gheibihayat, S., and Gholizadeh Navashenaq, J. (2021). Chemokine CXCL14; a double-edged sword in cancer development. Int. Immunopharmacol. 97, 107681. 10.1016/j.intimp.2021.107681.

55. Barrio-Alonso, C., Nieto-Valle, A., García-Martínez, E., Gutiérrez-Seijo, A., Parra-Blanco, V., Márquez-Rodas, I., Avilés-Izquierdo, J.A., Sánchez-Mateos, P., and Samaniego, R. (2024). Chemokine profiling of melanoma-macrophage crosstalk identifies CCL8 and CCL15 as prognostic factors in cutaneous melanoma. J. Pathol. 262, 495–504. 10.1002/path.6252.

56. Hickey, W.F., and Kimura, H. (1988). Perivascular Microglial Cells of the CNS Are Bone Marrow-Derived and Present Antigen in Vivo. Science. 10.1126/science.3276004.

57. Di Domizio, J., Belkhodja, C., Chenuet, P., Fries, A., Murray, T., Mondéjar, P.M., Demaria, O., Conrad, C., Homey, B., Werner, S., et al. (2020). The commensal skin microbiota triggers type I IFN-dependent innate repair responses in injured skin. Nat. Immunol. 21, 1034– 1045. 10.1038/s41590-020-0721-6.

58. Tripathi, A., Singh Rawat, B., Addya, S., Surjit, M., Tailor, P., Vrati, S., and Banerjee, A. (2021). Lack of Interferon (IFN) Regulatory Factor 8 Associated with Restricted IFN-γ Response Augmented Japanese Encephalitis Virus Replication in the Mouse Brain. J. Virol. 95, 10.1128/jvi.00406-21. 10.1128/jvi.00406-21.

59. Simonis, A., Fux, M., Nair, G., Mueller, N.J., Haralambieva, E., Pabst, T., Pachlopnik Schmid, J., Schmidt, A., Schanz, U., Manz, M.G., et al. (2018). Allogeneic hematopoietic cell transplantation in patients with GATA2 deficiency-a case report and comprehensive review of the literature. Ann. Hematol. 97, 1961–1973. 10.1007/s00277-018-3388-4.

60. André, A.C., Laborde, M., and Marteyn, B.S. (2022). The battle for oxygen during bacterial and fungal infections. Trends Microbiol. 30, 643–653. 10.1016/j.tim.2022.01.002.

61. Cooper, A.M., Magram, J., Ferrante, J., and Orme, I.M. (1997). Interleukin 12 (IL-12) Is Crucial to the Development of Protective Immunity in Mice Intravenously Infected with Mycobacterium tuberculosis. J. Exp. Med. 186, 39–45.

62. Mattila, J.T., Ojo, O.O., Kepka-Lenhart, D., Marino, S., Kim, J.H., Eum, S.Y., Via, L.E., Barry, C.E., Klein, E., Kirschner, D.E., et al. (2013). Microenvironments in tuberculous granulomas are delineated by distinct populations of macrophage subsets and expression of nitric oxide synthase and arginase isoforms. J. Immunol. Baltim. Md 1950 191, 773–784. 10.4049/jimmunol.1300113.

63. Camilli, C., Hoeh, A.E., De Rossi, G., Moss, S.E., and Greenwood, J. (2022). LRG1: an emerging player in disease pathogenesis. J. Biomed. Sci. 29, 6. 10.1186/s12929-022-00790-6.

64. Ojha, K.R., Padgham, S., Shin, S.Y., Woodman, C.R., and Trache, A. (2022). LPA effect on integrin recruitment and actin remodeling in aged vascular smooth muscle cells. Biophys. J. 121, 491a–492a. 10.1016/j.bpj.2021.11.316.

65. Dong, H., Zhu, G., Tamada, K., and Chen, L. (1999). B7-H1, a third member of the B7 family, co-stimulates T-cell proliferation and interleukin-10 secretion. Nat. Med. 5, 1365–1369. 10.1038/70932.

66. Lohrmann, F., Forde, A.J., Merck, P., and Henneke, P. (2021). Control of myeloid cell density in barrier tissues. FEBS J. 288, 405–426. 10.1111/febs.15436.

67. Sieweke, M.H., and Allen, J.E. (2013). Beyond stem cells: self-renewal of differentiated macrophages. Science 342, 1242974. 10.1126/science.1242974.

68. Chakarov, S., Lim, H.Y., Tan, L., Lim, S.Y., See, P., Lum, J., Zhang, X.-M., Foo, S., Nakamizo, S., Duan, K., et al. (2019). Two distinct interstitial macrophage populations coexist across tissues in specific subtissular niches. Science 363. 10.1126/science.aau0964.

69. Liu, Z., Gu, Y., Chakarov, S., Bleriot, C., Kwok, I., Chen, X., Shin, A., Huang, W., Dress, R.J., Dutertre, C.-A., et al. (2019). Fate Mapping via Ms4a3-Expression History Traces Monocyte-Derived Cells. Cell 178, 1509–1525.e19. 10.1016/j.cell.2019.08.009.

70. Reynolds, G., Vegh, P., Fletcher, J., Poyner, E.F.M., Stephenson, E., Goh, I., Botting, R.A., Huang, N., Olabi, B., Dubois, A., et al. (2021). Developmental cell programs are co-opted in inflammatory skin disease. Science 371, eaba6500. 10.1126/science.aba6500.

71. Sanin, D.E., Ge, Y., Marinkovic, E., Kabat, A.M., Castoldi, A., Caputa, G., Grzes, K.M., Curtis, J.D., Thompson, E.A., Willenborg, S., et al. (2022). A common framework of monocyte-derived macrophage activation. Sci. Immunol. 7, eabl7482. 10.1126/sciimmunol.abl7482.

72. Lavin, Y., Winter, D., Blecher-Gonen, R., David, E., Keren-Shaul, H., Merad, M., Jung, S., and Amit, I. (2014). Tissue-Resident Macrophage Enhancer Landscapes Are Shaped by the Local Microenvironment. Cell 159, 1312–1326. 10.1016/j.cell.2014.11.018.

73. Ehlers, S., and Schaible, U. (2013). The Granuloma in Tuberculosis: Dynamics of a Host–Pathogen Collusion. Front. Immunol. 3.

74. Warsinske, H.C., DiFazio, R.M., Linderman, J.J., Flynn, J.L., and Kirschner, D.E. (2017). Identifying mechanisms driving formation of granuloma-associated fibrosis during Mycobacterium tuberculosis infection. J. Theor. Biol. 429, 1–17. 10.1016/j.jtbi.2017.06.017.

75. Wynn, T.A., and Vannella, K.M. (2016). Macrophages in Tissue Repair, Regeneration, and Fibrosis. Immunity 44, 450–462. 10.1016/j.immuni.2016.02.015.

76. Pagán, A.J., Yang, C.-T., Cameron, J., Swaim, L.E., Ellett, F., Lieschke, G.J., and Ramakrishnan, L. (2015). Myeloid Growth Factors Promote Resistance to Mycobacterial Infection by Curtailing Granuloma Necrosis through Macrophage Replenishment. Cell Host Microbe 18, 15–26. 10.1016/j.chom.2015.06.008.

77. Bryden, S.R., Pingen, M., Lefteri, D.A., Miltenburg, J., Delang, L., Jacobs, S., Abdelnabi, R., Neyts, J., Pondeville, E., Major, J., et al. (2020). Pan-viral protection against arboviruses by activating skin macrophages at the inoculation site. Sci. Transl. Med. 12, eaax2421. 10.1126/scitranslmed.aax2421.

78. Hoeffel, G., Debroas, G., Roger, A., Rossignol, R., Gouilly, J., Laprie, C., Chasson, L., Barbon, P.-V., Balsamo, A., Reynders, A., et al. (2021). Sensory neuron-derived TAFA4 promotes macrophage tissue repair functions. Nature 594, 94–99. 10.1038/s41586-021-03563-7.

79. Tanaka, T., Okuda, H., Isonishi, A., Terada, Y., Kitabatake, M., Shinjo, T., Nishimura, K., Takemura, S., Furue, H., Ito, T., et al. (2023). Dermal macrophages set pain sensitivity by modulating the amount of tissue NGF through an SNX25-Nrf2 pathway. Nat. Immunol. 24, 439–451. 10.1038/s41590-022-01418-5.

