## Supplemental figures for "Tissue imprinting defines functional mosaic of dermal macrophages"

Fig. S1.1: supplementing figure 1

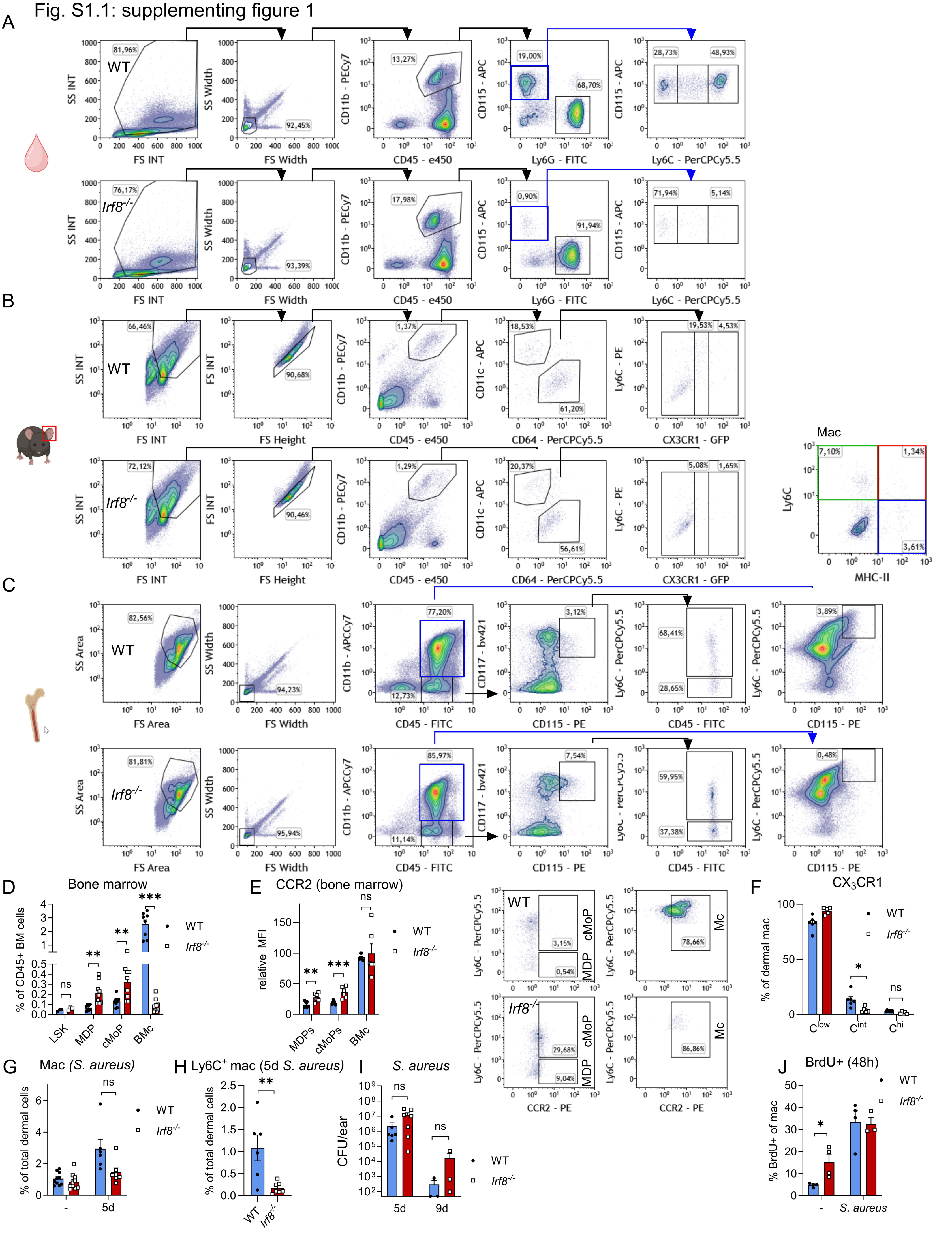

Fig. S1.2: supplementing figure 1

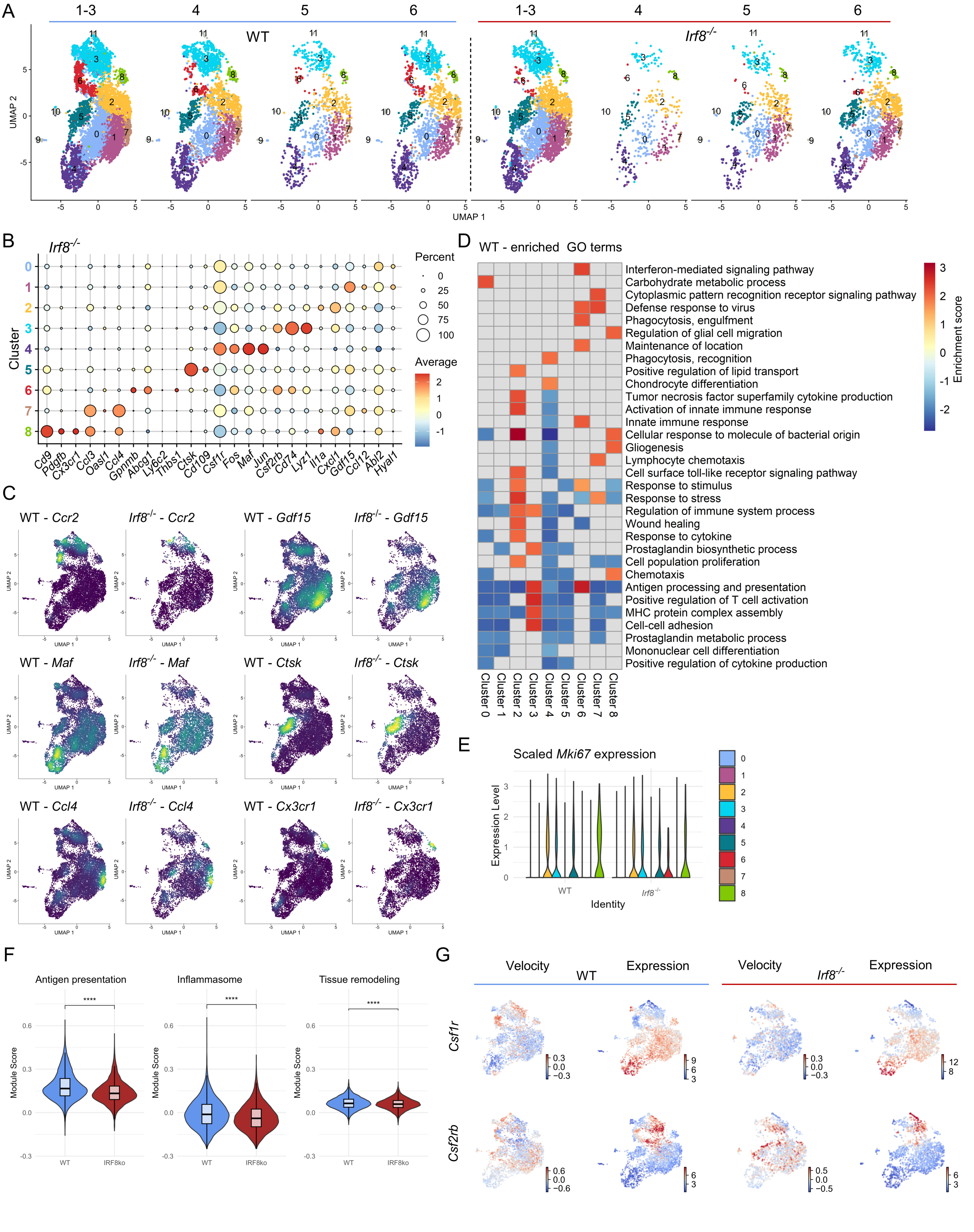

Fig. S2: supplementing figure 2

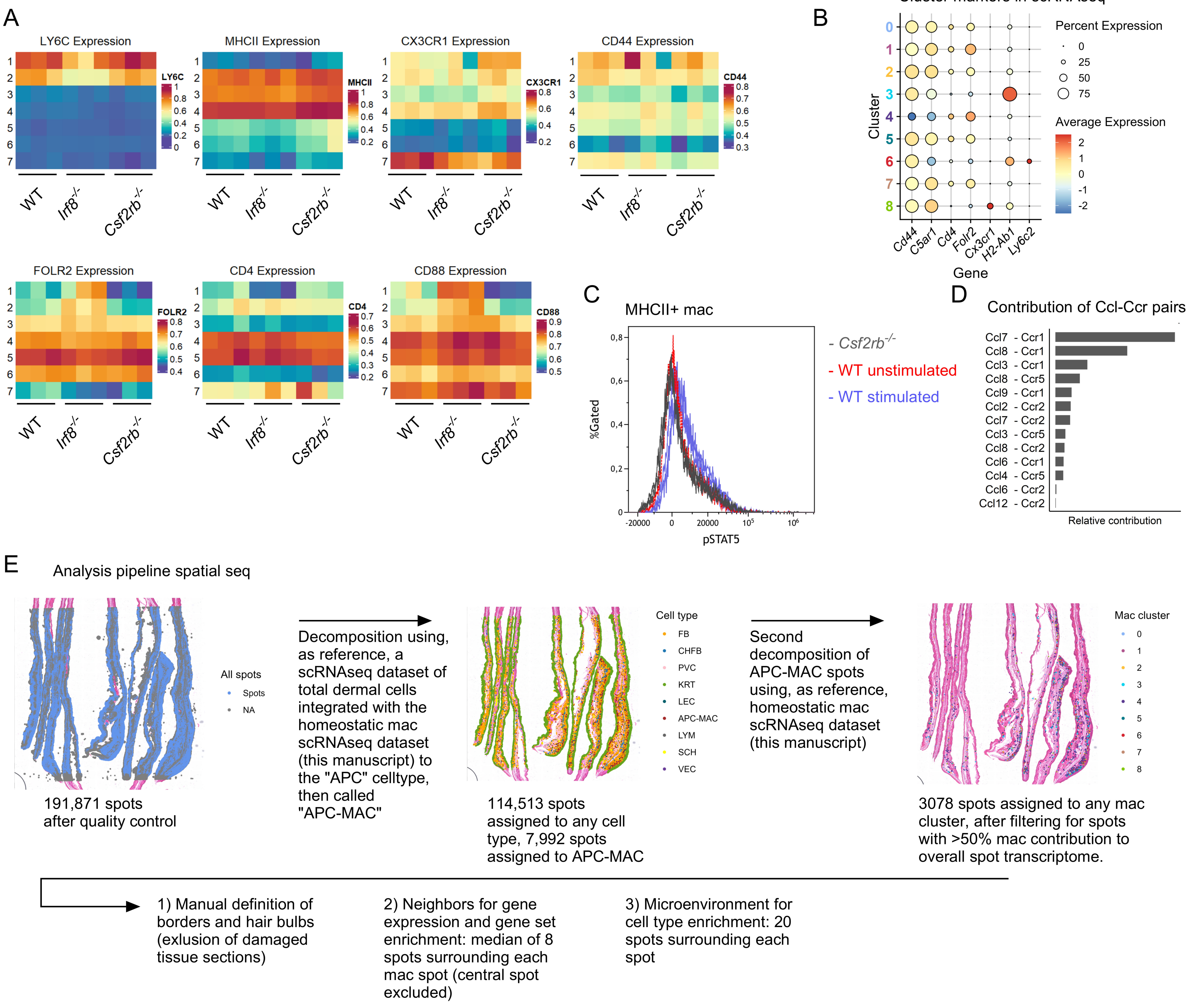

Fig. S3: supplementing fig. 3

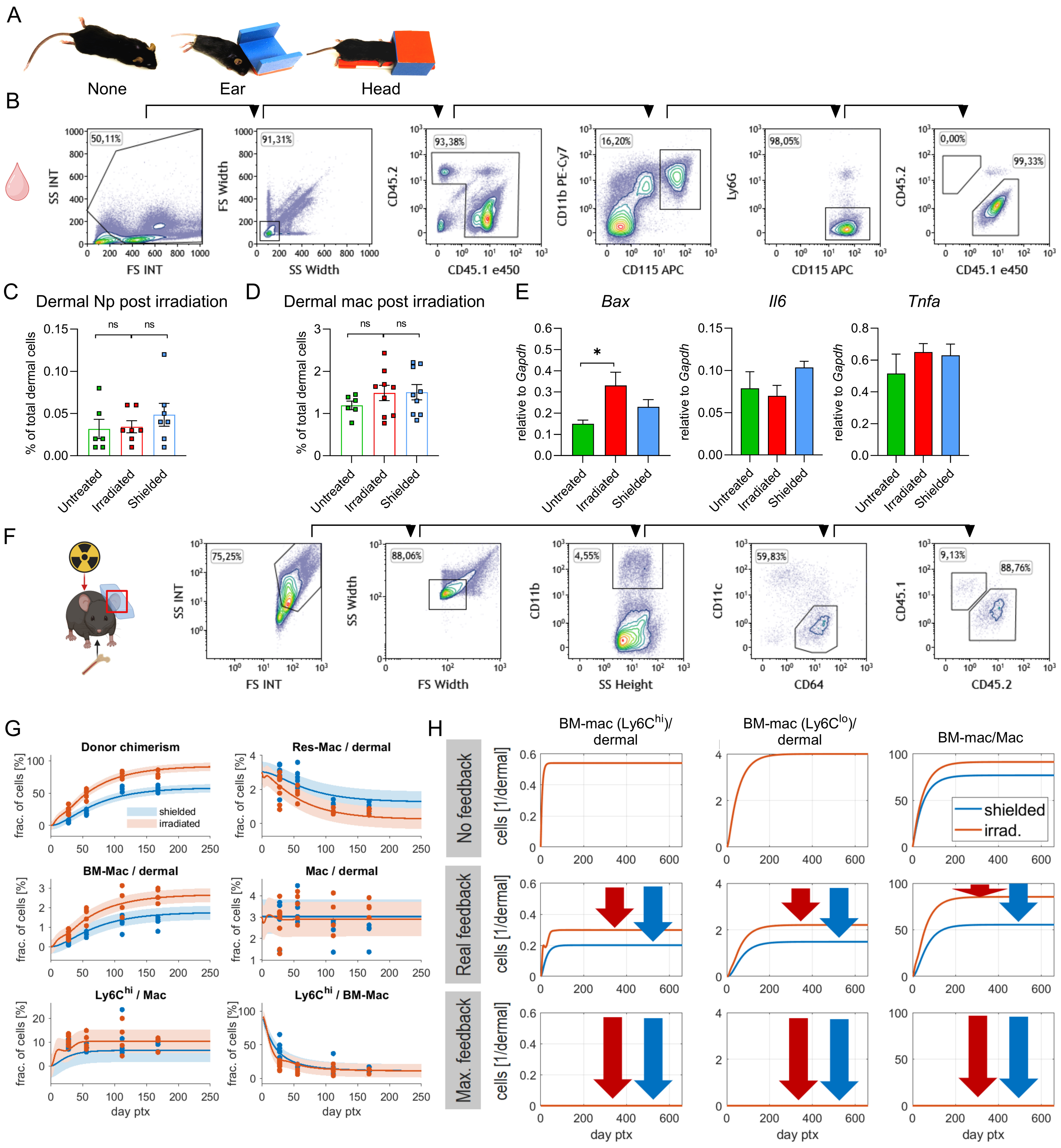

Fig. S4 supplementing figure 4

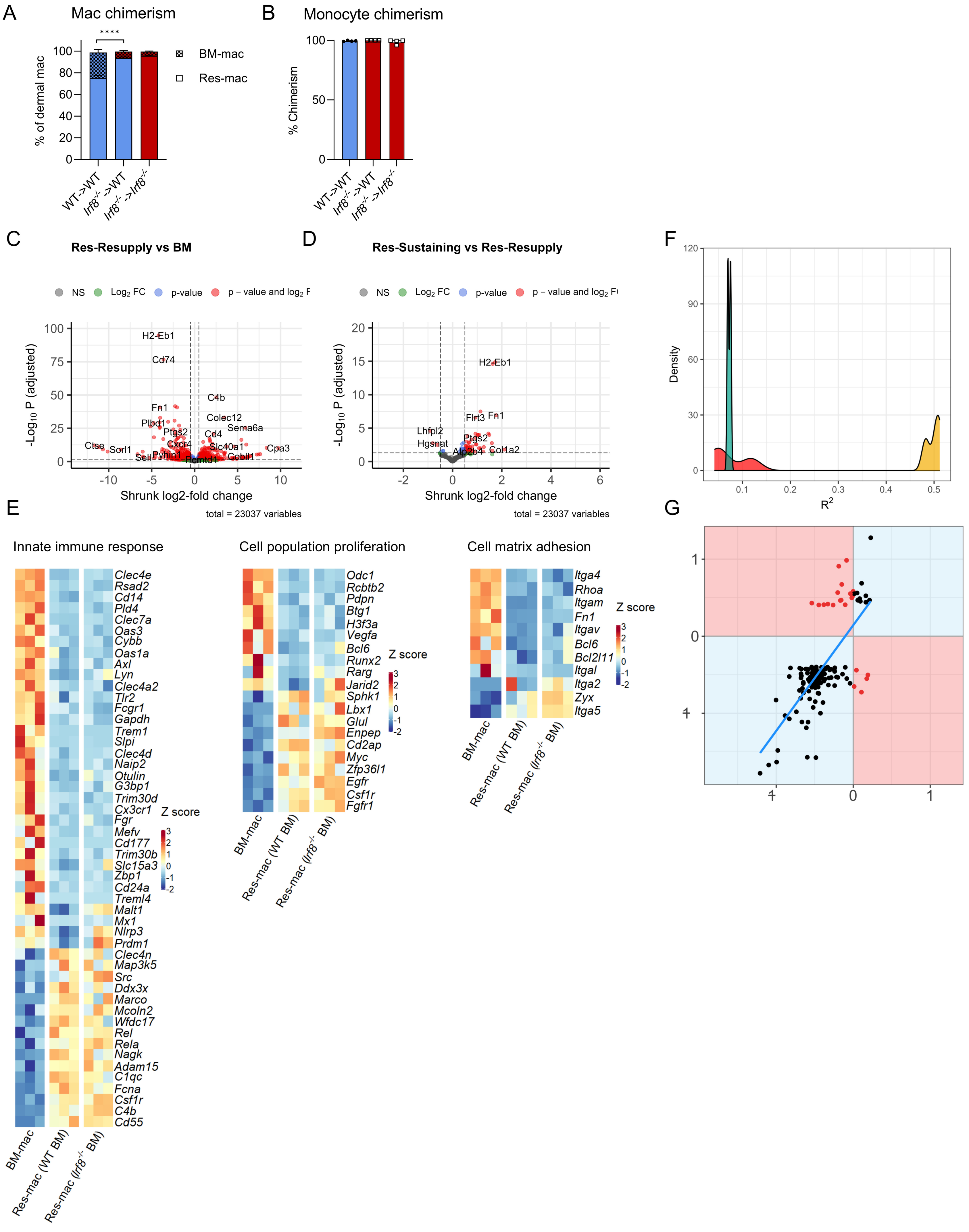

Fig. S5 supplementing fig. 5

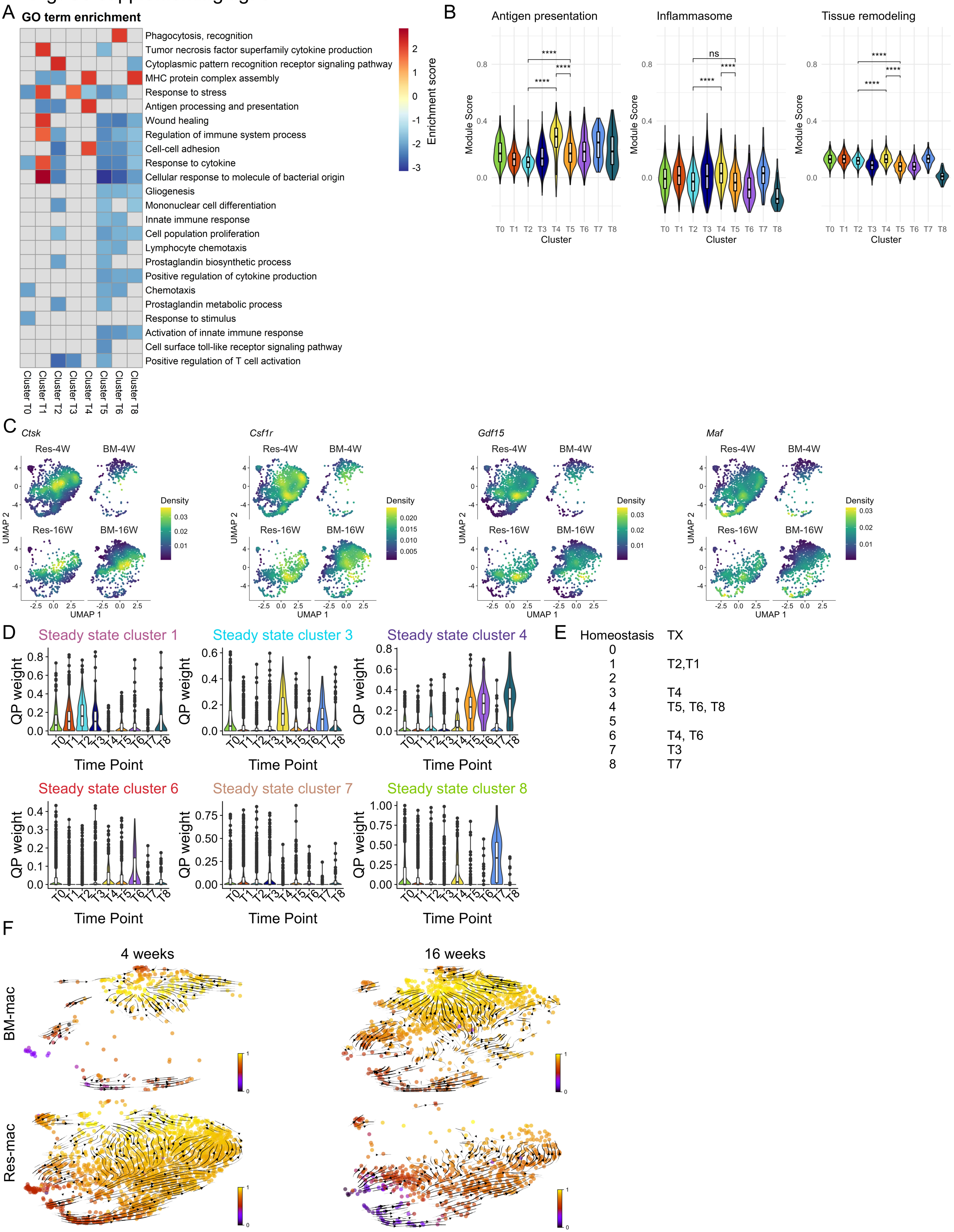

Fig. S6 supplementing figure 6

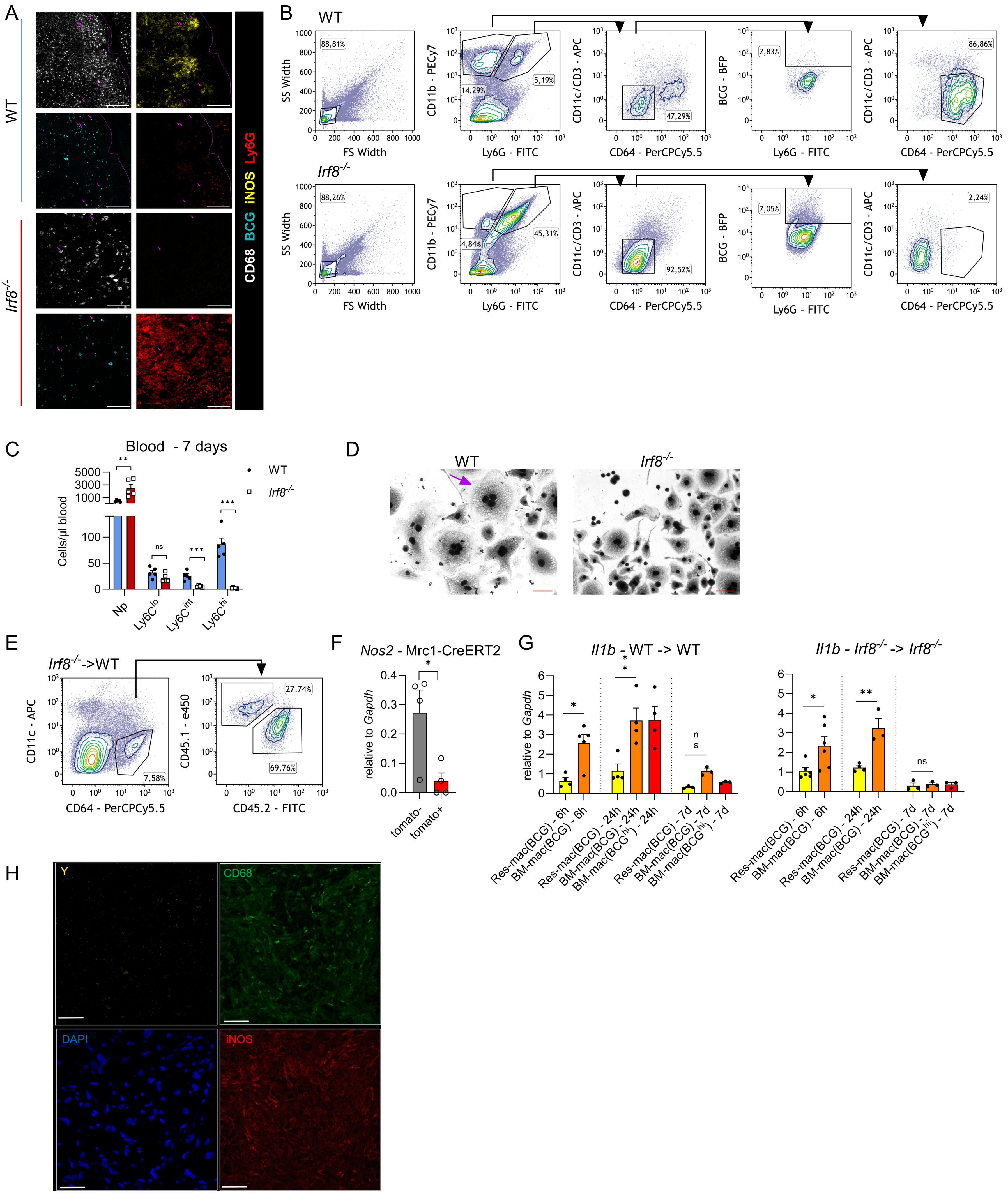

### Fig. S7: supplementing figure 7

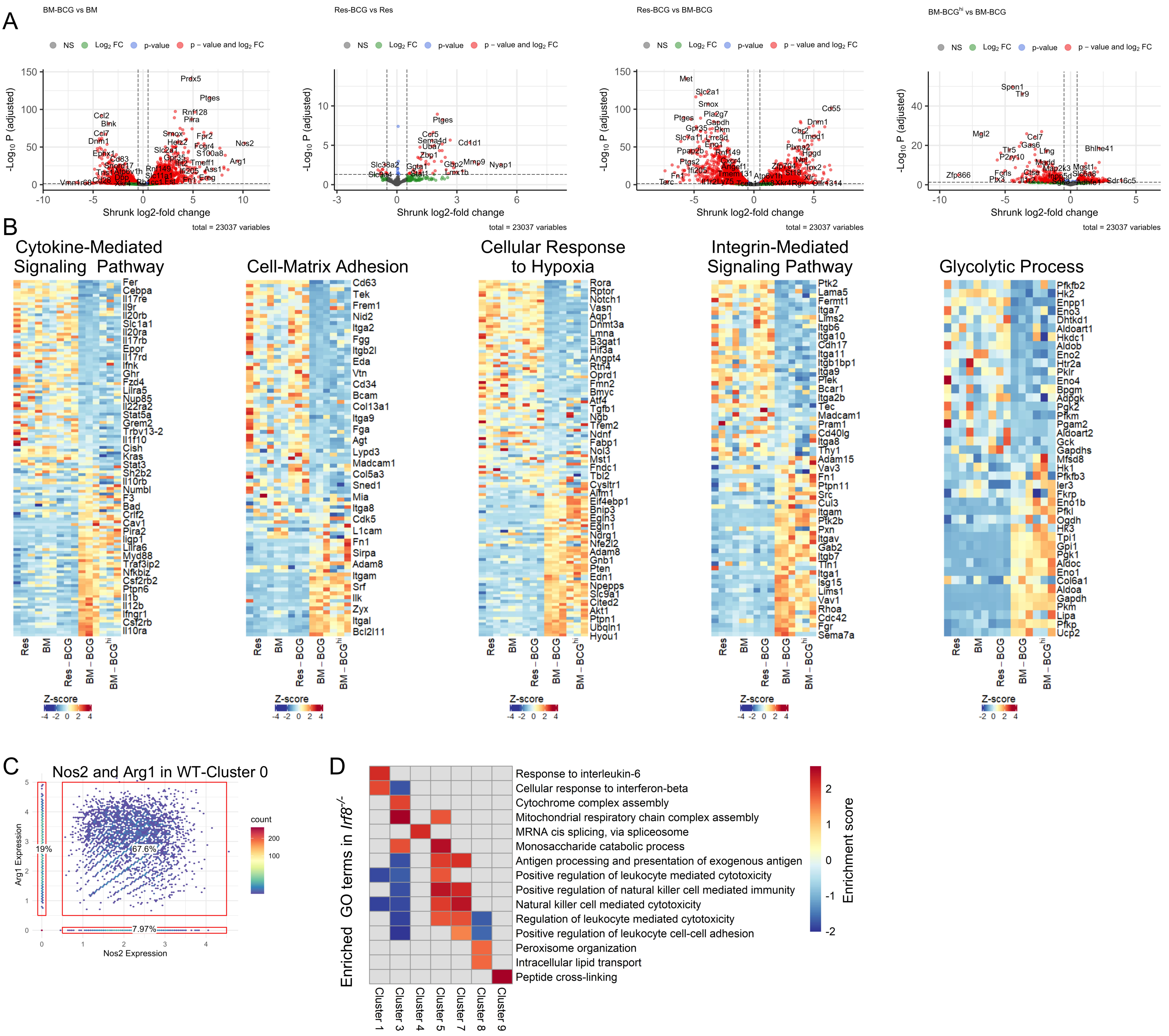
