## Supplemental tables for "Tissue imprinting defines functional mosaic of dermal macrophages": Data2Dynamics_modeling_report.pdf

### Data2Dynamics Software – Modeling Report

ckreutz

27-Jan-2023 11:00:21

**Website:** http://www.data2dynamics.org

**Key reference:**

- Data2Dynamics: a modeling environment tailored to parameter estimation in dynamical systems. A. Raue, B. Steiert, M. Schelker, C. Kreutz, T. Maiwald, H. Hass, J. Vanlier, C. Tönsing, L. Adlung, R. Engesser, W. Mader, T. Heinemann, J. Hasenauer, M. Schilling, T. Höfer, E. Klipp, F. Theis, U. Klingmüller, B. Schoeberl and J. Timmer. *Bioinformatics*, **31**(21), 3558-3560, 2015.
- Lessons learned from quantitative dynamical modeling in systems biology. A. Raue, M. Schilling, J. Bachmann, A. Matteson, M. Schelker, D. Kaschek, S. Hug, C. Kreutz, BD. Harms, F. Theis, U. Klingmüller, and J. Timmer. *PLOS ONE*, **8**(9), e74335, 2013.

#### Contents


**1 Model: HalfLifeModel 3**
1.1 Comments 3
1.2 Dynamic variables 3
1.3 Reactions 3
1.4 ODE system 4
1.5 Observables 4
1.6 Conditions 4
1.7 Experiment: Cluster_numbers_27Jan2023_ T0 5
1.7.1 Comments 5
1.7.2 Experiment specific conditions 5
1.7.3 Experimental data and model fit 5
1.8 Experiment: Cluster_numbers_27Jan2023_ T1 8
1.8.1 Comments 8
1.8.2 Experiment specific conditions 8
1.8.3 Experimental data and model fit 8
1.9 Experiment: Cluster_numbers_27Jan2023_ T2 11
1.9.1 Comments 11
1.9.2 Experiment specific conditions 11
1.9.3 Experimental data and model fit 11
1.10 Experiment: Cluster_numbers_27Jan2023_ T3 14
1.10.1 Comments 14
1.10.2 Experiment specific conditions 14
1.10.3 Experimental data and model fit 14
1.11 Experiment: Cluster_numbers_27Jan2023_ T4 17
1.11.1 Comments 17
1.11.2 Experiment specific conditions 17

1.11.3 Experimental data and model fit . . . 17
1.12 Experiment: Cluster_numbers_27Jan2023_ T5 . . . 20
1.12.1 Comments . . . 20
1.12.2 Experiment specific conditions . . . 20
1.12.3 Experimental data and model fit . . . 20
1.13 Experiment: Cluster_numbers_27Jan2023_ T6 . . . 23
1.13.1 Comments . . . 23
1.13.2 Experiment specific conditions . . . 23
1.13.3 Experimental data and model fit . . . 23
1.14 Experiment: Cluster_numbers_27Jan2023_ T7 . . . 26
1.14.1 Comments . . . 26
1.14.2 Experiment specific conditions . . . 26
1.14.3 Experimental data and model fit . . . 26
1.15 Experiment: Cluster_numbers_27Jan2023_ T8 . . . 29
1.15.1 Comments . . . 29
1.15.2 Experiment specific conditions . . . 29
1.15.3 Experimental data and model fit . . . 29

**2 Estimated model parameters** **32**

**3 Profile likelihood of model parameters** **33**

**4 Confidence intervals for the model parameters** **35**

### 1 Model: HalfLifeModel

#### 1.1 Comments

model .def file template

#### 1.2 Dynamic variables

The model contains 2 dynamic variables. The dynamics of those variables evolve according to a system of ordinary differential equations (ODE) as will be defined in the following. The following list indicates the unique variable names and their initial conditions.

- **Dynamic variable 1:** Donor

  $$[\mathrm{Donor}](\mathrm{t}=0) = \mathrm{init\_Donor}$$

- **Dynamic variable 2:** Recipient

  $$[\mathrm{Recipient}](\mathrm{t}=0) = \mathrm{init\_Recipient}$$

#### 1.3 Reactions

The model contains 3 reactions. Reactions define interactions between dynamics variables and build up the ODE systems. The following list indicates the reaction laws and their corresponding reaction rate equations. Promoting rate modifiers are indicated in black above the rate law arrow. Inhibitory rate modifiers are indicated in red below the rate law arrow. In the reaction rate equations dynamic and input variables are indicated by square brackets. The remaining variables are model parameters that remain constant over time.

- **Reaction 1:**

  $$\varnothing \longrightarrow \mathrm{Donor}$$

  $$v_1 = \mathrm{kin\_ClusterID}$$

- **Reaction 2:**

  $$\mathrm{Donor} \longrightarrow \varnothing$$

  $$v_2 = \frac{[\mathrm{Donor}] \cdot \ln(2)}{\mathrm{HalfLife\_ClusterID}}$$

- **Reaction 3:**

  $$\mathrm{Recipient} \longrightarrow \varnothing$$

  $$v_3 = \frac{[\mathrm{Recipient}] \cdot \ln(2)}{\mathrm{HalfLife\_ClusterID}}$$

#### 1.4 ODE system

The specified reation laws and rate equations $v$ determine an ODE system. The time evolution of the dynamical variables is calculated by solving this equation system.

$$\begin{aligned} \mathrm{d[Donor]/dt} &= +v_1 - v_2 \\ \mathrm{d[Recipient]/dt} &= -v_3 \end{aligned}$$

The ODE system was solved by a parallelized implementation of the CVODES algorithm [**?** ]. It also supplies the parameter sensitivities utilized for parameter estimation.

#### 1.5 Observables

The model contains 3 standard observables. Observables are calculated after the ODE system was solved and derived variables are calculated. Dynamic, input and derived variables are indicated by square brackets. The remaining variables are model parameters that remain constant over time. In additon to the equation for the observable, also their corresponding error model $\sigma$ is indicated.

- **Observable 1:** DonorChimerism

$$\begin{aligned} \mathrm{DonorChimerism(t)} &= \frac{100 \cdot \mathrm{[Donor]}}{\mathrm{[Donor]} + \mathrm{[Recipient]}} \\ \sigma\{\mathrm{DonorChimerism}\}(\mathrm{t}) &= \mathrm{SD2} \end{aligned}$$

- **Observable 2:** DonorCells

$$\begin{aligned} \mathrm{DonorCells(t)} &= \log_{10}(\mathrm{[Donor]} + 1) \\ \sigma\{\mathrm{DonorCells}\}(\mathrm{t}) &= \mathrm{SD} \end{aligned}$$

- **Observable 3:** RecipientCells

$$\begin{aligned} \mathrm{RecipientCells(t)} &= \log_{10}(\mathrm{[Recipient]} + 1) \\ \sigma\{\mathrm{RecipientCells}\}(\mathrm{t}) &= \mathrm{SD} \end{aligned}$$

#### 1.6 Conditions

Conditions modify the model according to replacement rules. New model parameters can be introduced or relations between existing model parameters can be implemented. The following list are default conditions that can be replace my experiment specific conditions defined seperately for each data set.

$$\begin{aligned} \mathrm{init\_Donor} &\rightarrow 0 \\ \mathrm{init\_Recipient} &\rightarrow \frac{\mathrm{HalfLife\_ClusterID} \cdot \mathrm{kin\_ClusterID}}{\ln(2)} \end{aligned}$$

| time [min] | DonorChimerism donor chimerism [%] | DonorCells cells [counts] | RecipientCells cells [counts] |
|---:|---:|---:|---:|
| 4 | 24.159 | 79 | 248 |
| 16 | 92.3868 | 449 | 37 |
| 52 | NaN | NaN | NaN |

**Table 1: Experimental data for the experiment Cluster_numbers_27Jan2023_ T0**

#### 1.7 Experiment: Cluster_numbers_27Jan2023_ T0

##### 1.7.1 Comments

data .def file template

##### 1.7.2 Experiment specific conditions

To evaluate the model for this experiment the following conditions are applied.

- **Local condition #1 (global condition #1):**

$$\mathrm{HalfLife\_ClusterID} \rightarrow \mathrm{HalfLife\_ClusterIDT0}$$

$$\mathrm{init\_Recipient} \rightarrow \frac{\mathrm{HalfLife\_ClusterIDT0} \cdot \mathrm{kin\_ClusterIDT0}}{\ln(2)}$$

$$\mathrm{kin\_ClusterID} \rightarrow \mathrm{kin\_ClusterIDT0}$$

##### 1.7.3 Experimental data and model fit

The model observables and the experimental data is shown in Figure 1. The agreement of the model observables and the experimental data, given in Table 1, yields a value of the objective function $\chi^2 = 9.1999$ for 4 data points in this data set. The trajectories of the input, dynamic and derived variables that correspond to the experimental conditions in this experiment are shown in Figure 2.

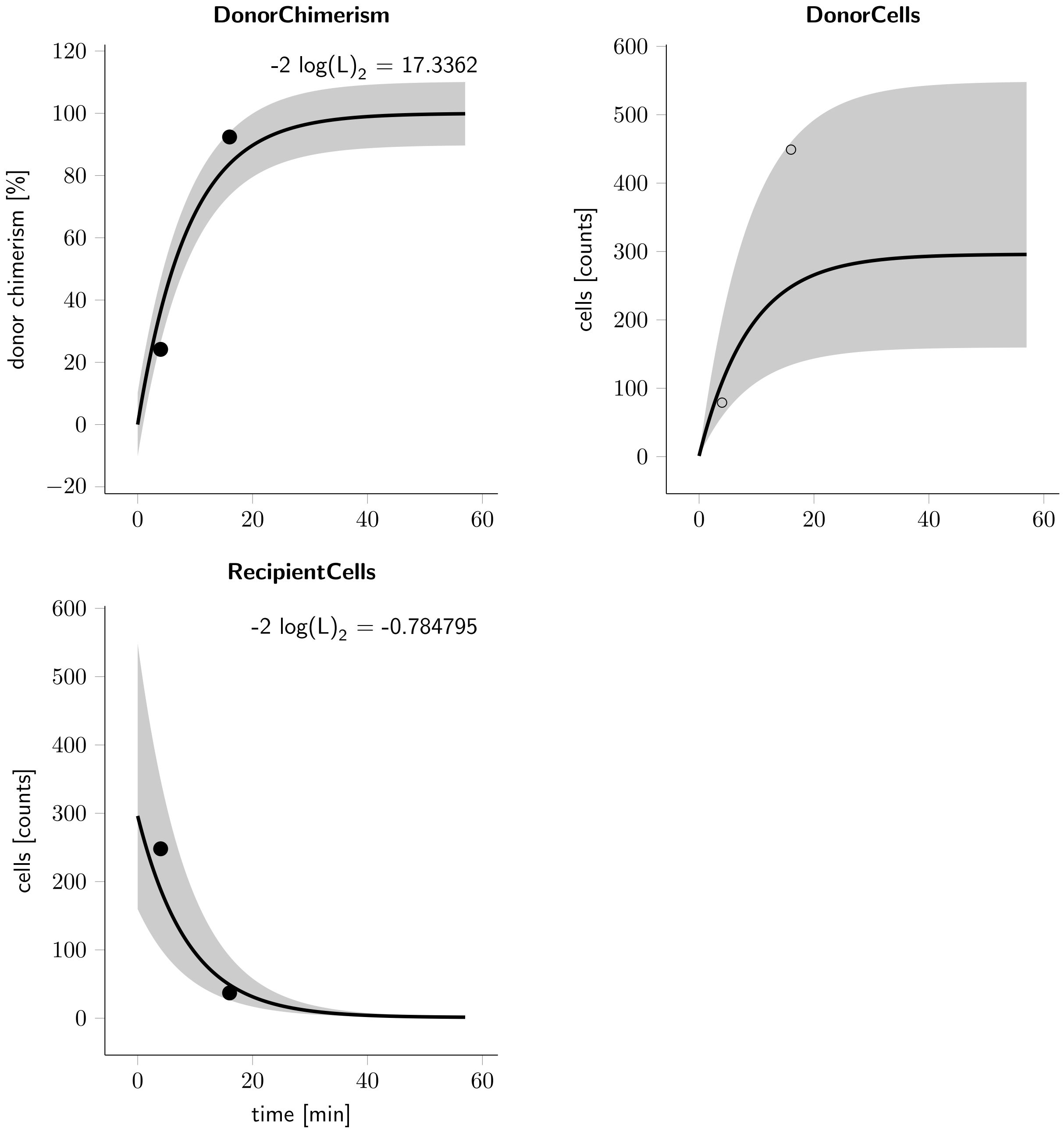


**Figure 1: Cluster_numbers_27Jan2023_ T0 observables and experimental data for the experiment.** The observables are displayed as solid lines. The error model that describes the measurement noise is indicated by shades.

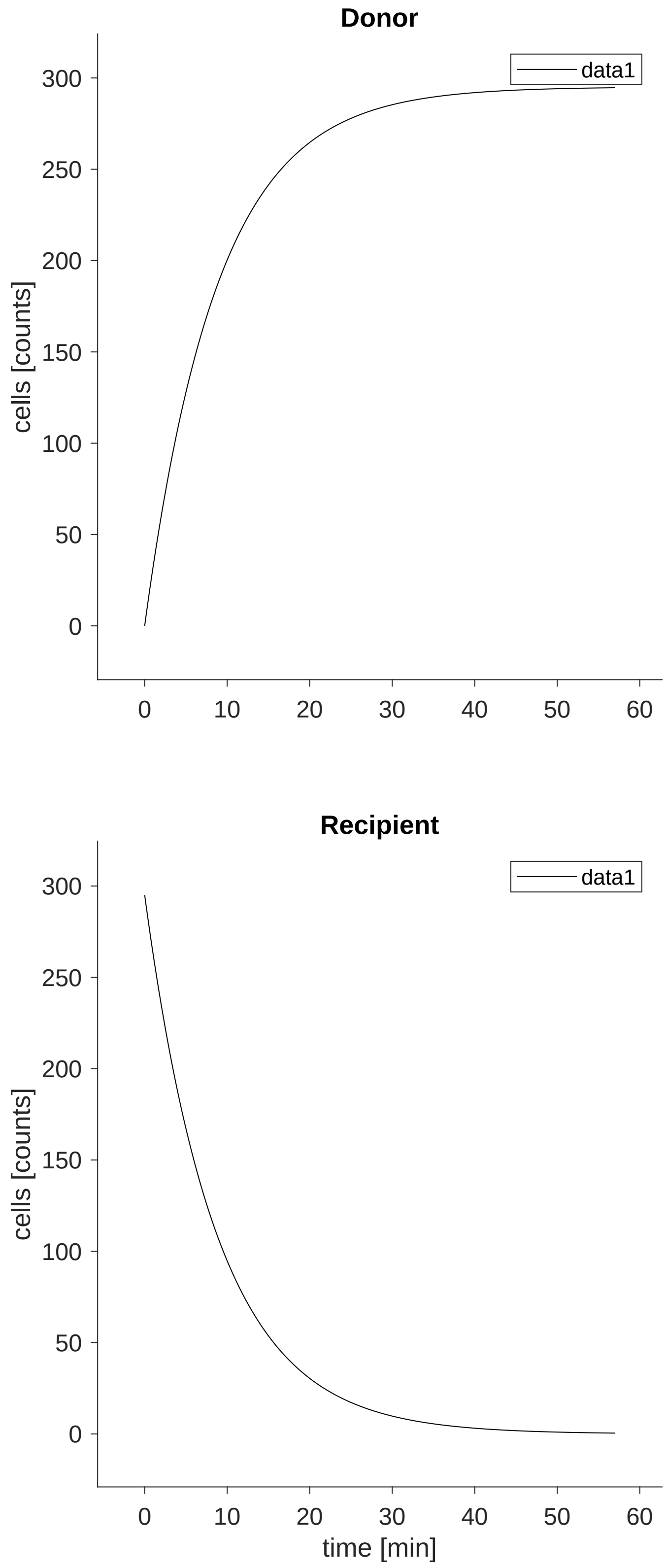




**Figure 2: Cluster_numbers_27Jan2023_ T0 trajectories of the input, dynamic and derived variables.** The dynamical behaviour is determined by the ODE system defined in Section
refHalfLifeModel$_{o}de$.

| time [min] | DonorChimerism donor chimerism [%] | DonorCells cells [counts] | RecipientCells cells [counts] |
|---|---|---|---|
| 4 | 2.90135 | 15 | 502 |
| 16 | 60.274 | 176 | 116 |
| 52 | NaN | NaN | NaN |

**Table 2: Experimental data for the experiment Cluster_numbers_27Jan2023_ T1**

#### 1.8 Experiment: Cluster_numbers_27Jan2023_ T1

##### 1.8.1 Comments

data .def file template

##### 1.8.2 Experiment specific conditions

To evaluate the model for this experiment the following conditions are applied.

- **Local condition #2 (global condition #2):**

$$\mathrm{HalfLife\_ClusterID} \rightarrow \mathrm{HalfLife\_ClusterIDT1}$$
$$\mathrm{init\_Recipient} \rightarrow \frac{\mathrm{HalfLife\_ClusterIDT1} \cdot \mathrm{kin\_ClusterIDT1}}{\ln(2)}$$
$$\mathrm{kin\_ClusterID} \rightarrow \mathrm{kin\_ClusterIDT1}$$

##### 1.8.3 Experimental data and model fit

The model observables and the experimental data is shown in Figure 3. The agreement of the model observables and the experimental data, given in Table 2, yields a value of the objective function $\chi^2 = 10.9802$ for 4 data points in this data set. The trajectories of the input, dynamic and derived variables that correspond to the experimental conditions in this experiment are shown in Figure 4.

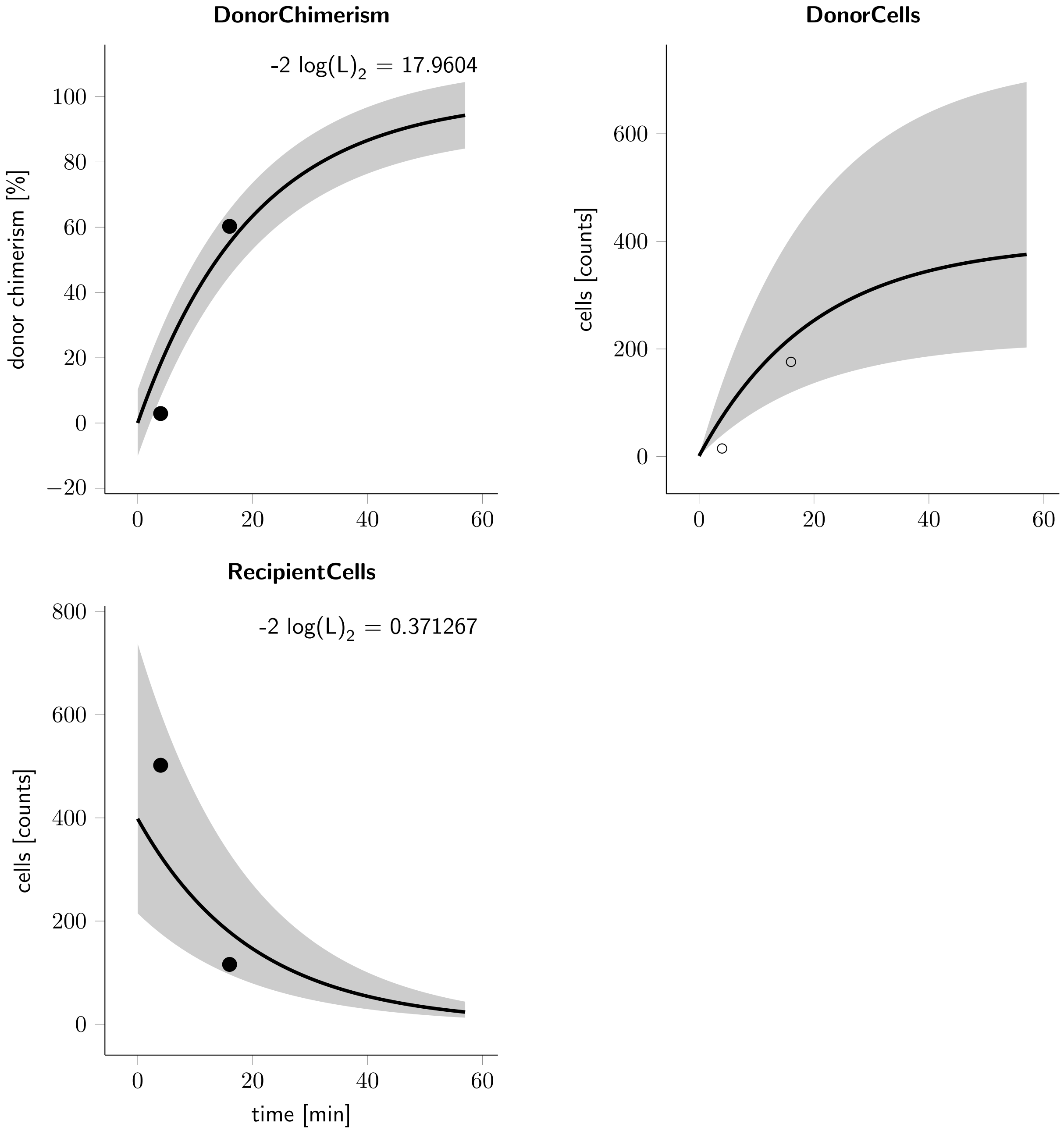


**Figure 3: Cluster_numbers_27Jan2023_ T1 observables and experimental data for the experiment.** The observables are displayed as solid lines. The error model that describes the measurement noise is indicated by shades.

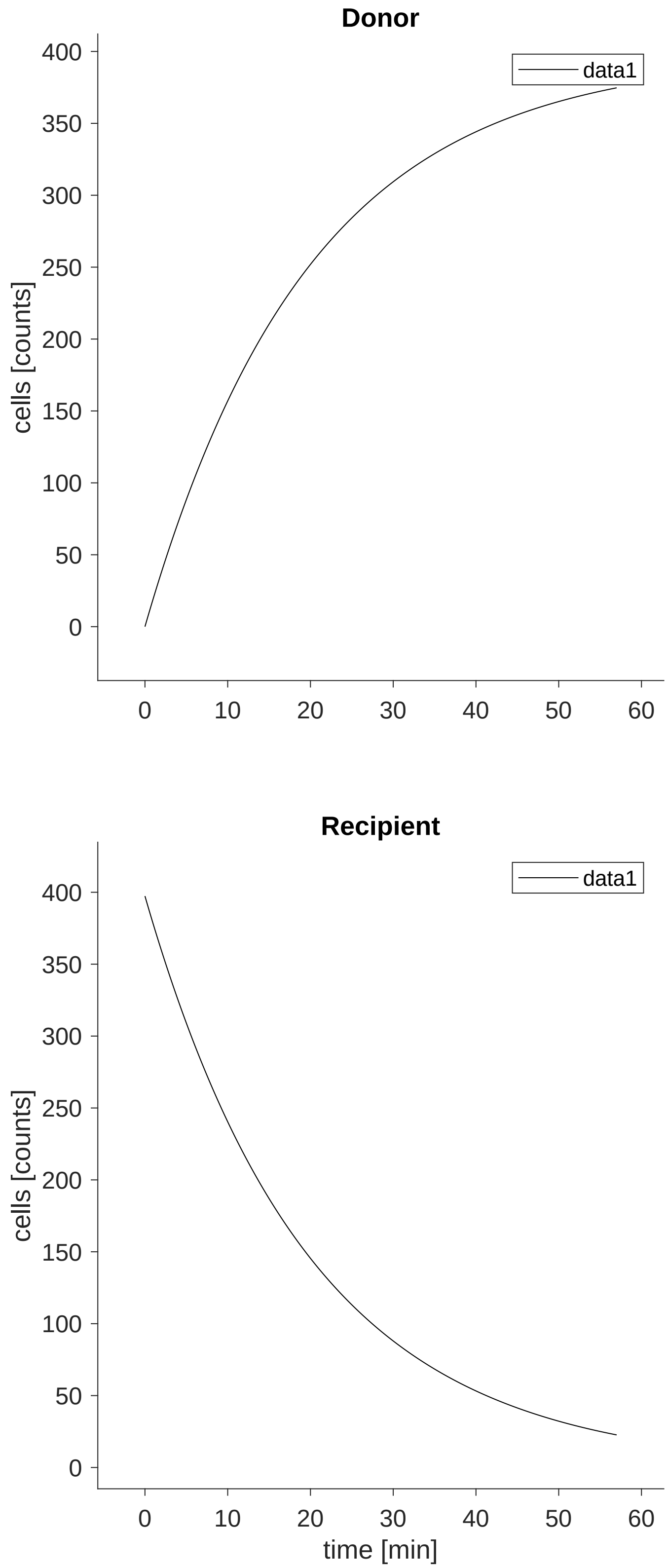


**Figure 4: Cluster_numbers_27Jan2023_ T1 trajectories of the input, dynamic and derived variables.** The dynamical behaviour is determined by the ODE system defined in Section
refHalfLifeModel$_{o}de$.

| time [min] | DonorChimerism donor chimerism [%] | DonorCells cells [counts] | RecipientCells cells [counts] |
|---|---|---|---|
| 4 | 0.779727 | 4 | 509 |
| 16 | 18.1467 | 47 | 212 |
| 52 | NaN | NaN | NaN |

**Table 3: Experimental data for the experiment Cluster_numbers_27Jan2023_ T2**

#### 1.9 Experiment: Cluster_numbers_27Jan2023_ T2

##### 1.9.1 Comments

data .def file template

##### 1.9.2 Experiment specific conditions

To evaluate the model for this experiment the following conditions are applied.

- **Local condition #3 (global condition #3):**

$$\mathrm{HalfLife\_ClusterID} \rightarrow \mathrm{HalfLife\_ClusterIDT2}$$
$$\mathrm{init\_Recipient} \rightarrow \frac{\mathrm{HalfLife\_ClusterIDT2} \cdot \mathrm{kin\_ClusterIDT2}}{\ln(2)}$$
$$\mathrm{kin\_ClusterID} \rightarrow \mathrm{kin\_ClusterIDT2}$$

##### 1.9.3 Experimental data and model fit

The model observables and the experimental data is shown in Figure 5. The agreement of the model observables and the experimental data, given in Table 3, yields a value of the objective function $\chi^2 = 5.72609$ for 4 data points in this data set. The trajectories of the input, dynamic and derived variables that correspond to the experimental conditions in this experiment are shown in Figure 6.

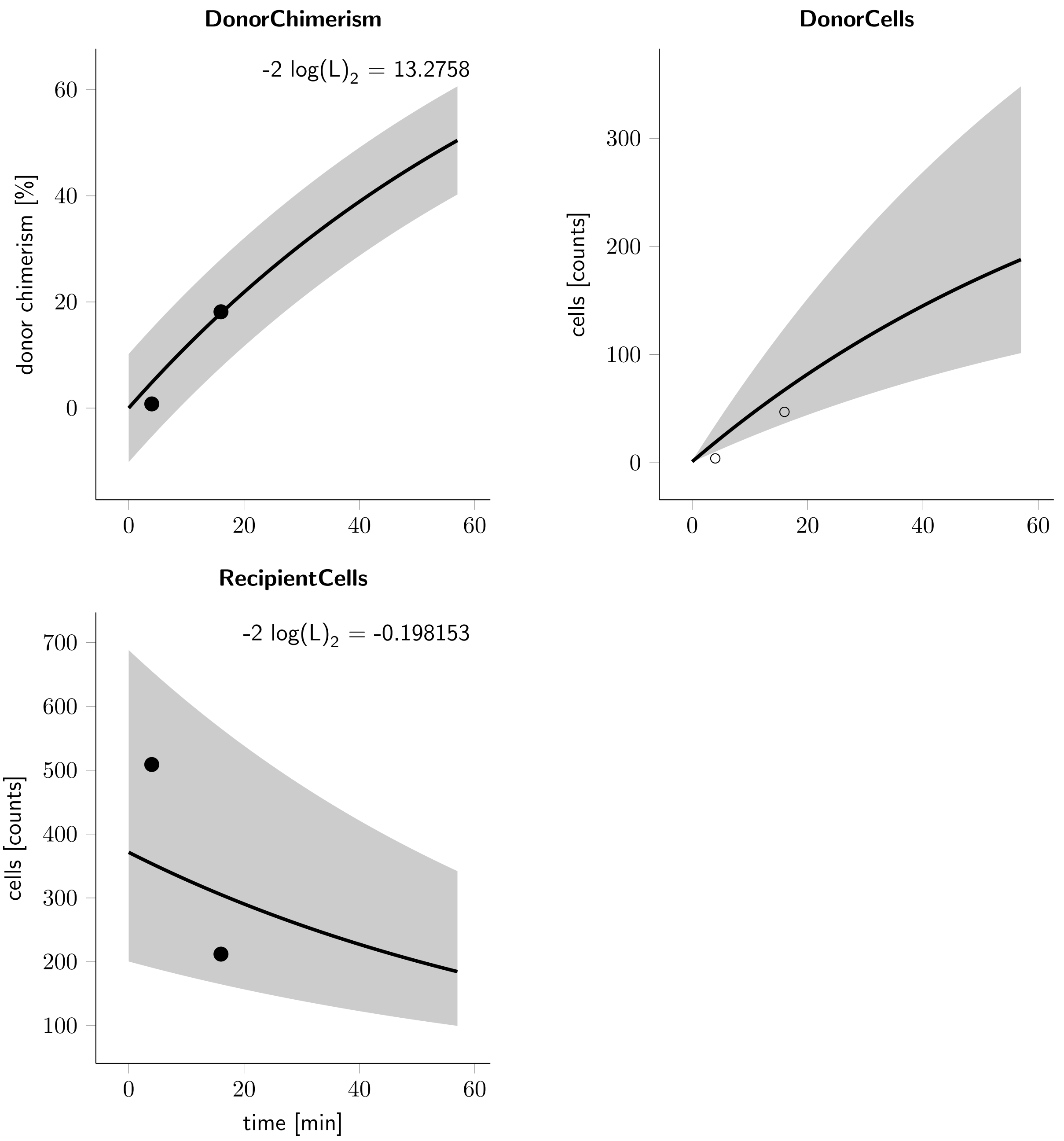


**Figure 5: Cluster_numbers_27Jan2023_ T2 observables and experimental data for the experiment.** The observables are displayed as solid lines. The error model that describes the measurement noise is indicated by shades.

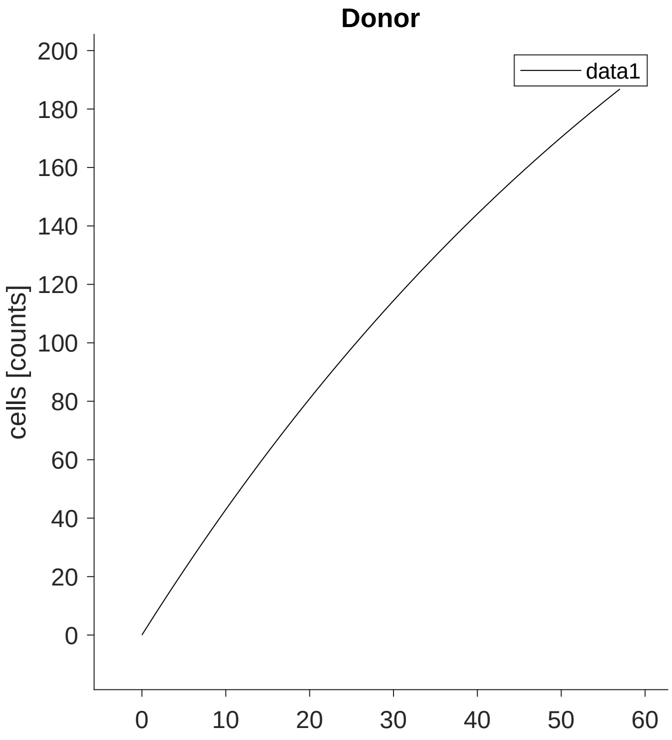


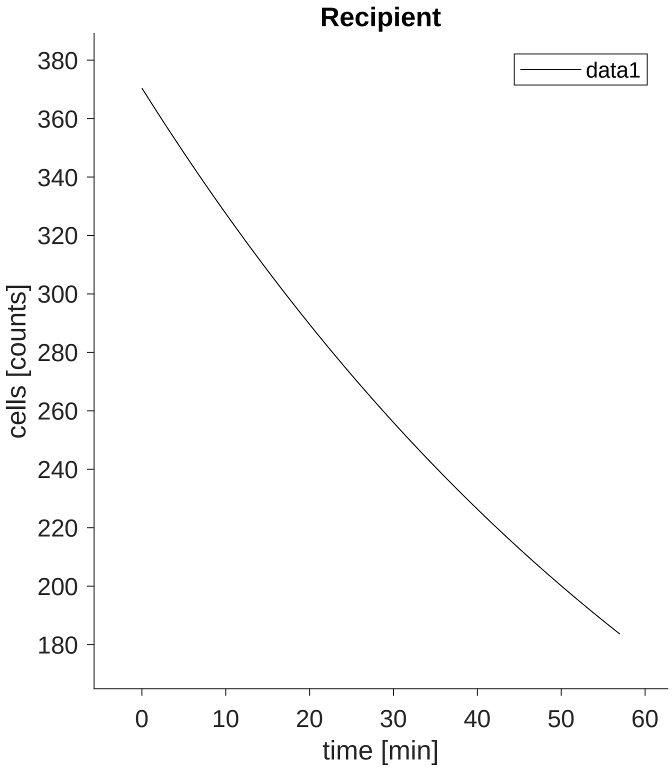




**Figure 6: Cluster_numbers_27Jan2023_ T2 trajectories of the input, dynamic and derived variables.** The dynamical behaviour is determined by the ODE system defined in Section
refHalfLifeModel$_ode$.

| time [min] | DonorChimerism donor chimerism [%] | DonorCells cells [counts] | RecipientCells cells [counts] |
|---|---|---|---|
| 4 | 10.9244 | 26 | 212 |
| 16 | 52.1084 | 173 | 159 |
| 52 | NaN | NaN | NaN |

**Table 4: Experimental data for the experiment Cluster_numbers_27Jan2023_ T3**

#### 1.10 Experiment: Cluster_numbers_27Jan2023_ T3

##### 1.10.1 Comments

data .def file template

##### 1.10.2 Experiment specific conditions

To evaluate the model for this experiment the following conditions are applied.

- **Local condition #4 (global condition #4):**

$$\text{HalfLife\_ClusterID} \rightarrow \text{HalfLife\_ClusterIDT3}$$
$$\text{init\_Recipient} \rightarrow \frac{\text{HalfLife\_ClusterIDT3} \cdot \text{kin\_ClusterIDT3}}{\ln(2)}$$
$$\text{kin\_ClusterID} \rightarrow \text{kin\_ClusterIDT3}$$

##### 1.10.3 Experimental data and model fit

The model observables and the experimental data is shown in Figure 7. The agreement of the model observables and the experimental data, given in Table 4, yields a value of the objective function $\chi^2 = 4.71667$ for 4 data points in this data set. The trajectories of the input, dynamic and derived variables that correspond to the experimental conditions in this experiment are shown in Figure 8.

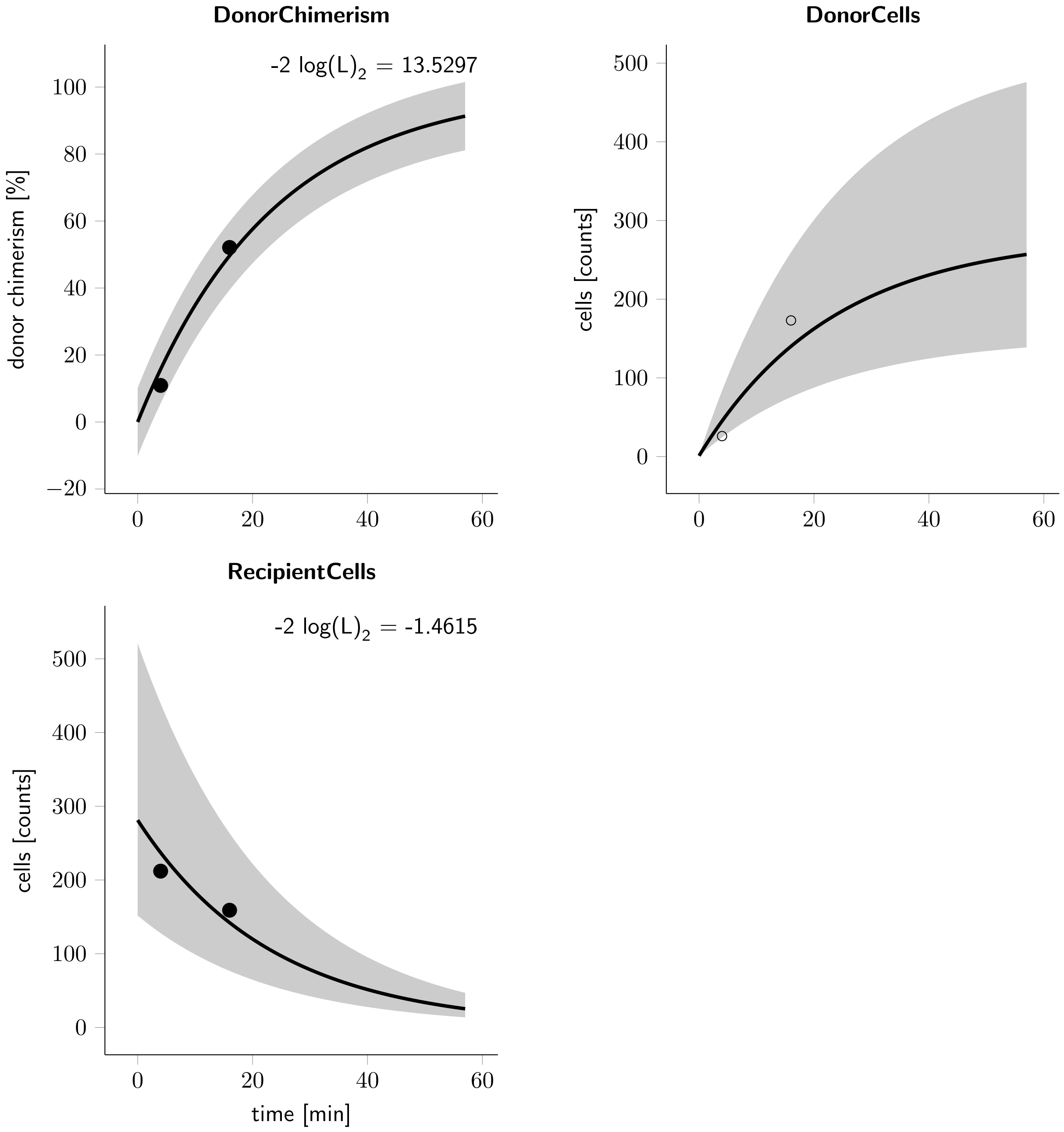


**Figure 7: Cluster_numbers_27Jan2023_ T3 observables and experimental data for the experiment.** The observables are displayed as solid lines. The error model that describes the measurement noise is indicated by shades.

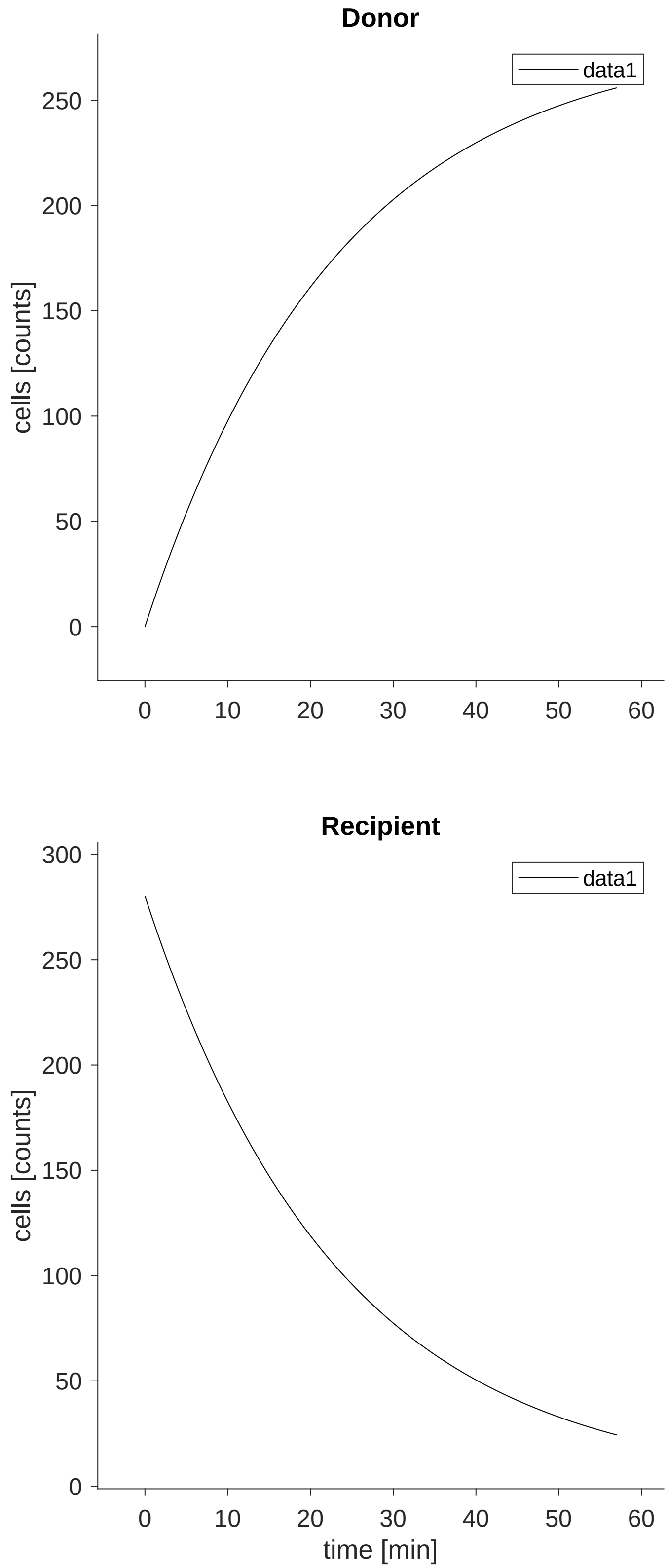




**Figure 8: Cluster_numbers_27Jan2023_ T3 trajectories of the input, dynamic and derived variables.** The dynamical behaviour is determined by the ODE system defined in Section refHalfLifeModel$_ode$.

| time [min] | DonorChimerism donor chimerism [%] | DonorCells cells [counts] | RecipientCells cells [counts] |
|---|---|---|---|
| 4 | 76.5101 | 114 | 35 |
| 16 | 98.0296 | 199 | 4 |
| 52 | NaN | NaN | NaN |

**Table 5: Experimental data for the experiment Cluster_numbers_27Jan2023_ T4**

#### 1.11 Experiment: Cluster_numbers_27Jan2023_ T4

##### 1.11.1 Comments

data .def file template

##### 1.11.2 Experiment specific conditions

To evaluate the model for this experiment the following conditions are applied.

- **Local condition #5 (global condition #5):**

$$\mathrm{HalfLife\_ClusterID} \rightarrow \mathrm{HalfLife\_ClusterIDT4}$$
$$\mathrm{init\_Recipient} \rightarrow \frac{\mathrm{HalfLife\_ClusterIDT4} \cdot \mathrm{kin\_ClusterIDT4}}{\ln(2)}$$
$$\mathrm{kin\_ClusterID} \rightarrow \mathrm{kin\_ClusterIDT4}$$

##### 1.11.3 Experimental data and model fit

The model observables and the experimental data is shown in Figure 9. The agreement of the model observables and the experimental data, given in Table 5, yields a value of the objective function $\chi^2 = 7.01642$ for 4 data points in this data set. The trajectories of the input, dynamic and derived variables that correspond to the experimental conditions in this experiment are shown in Figure 10.

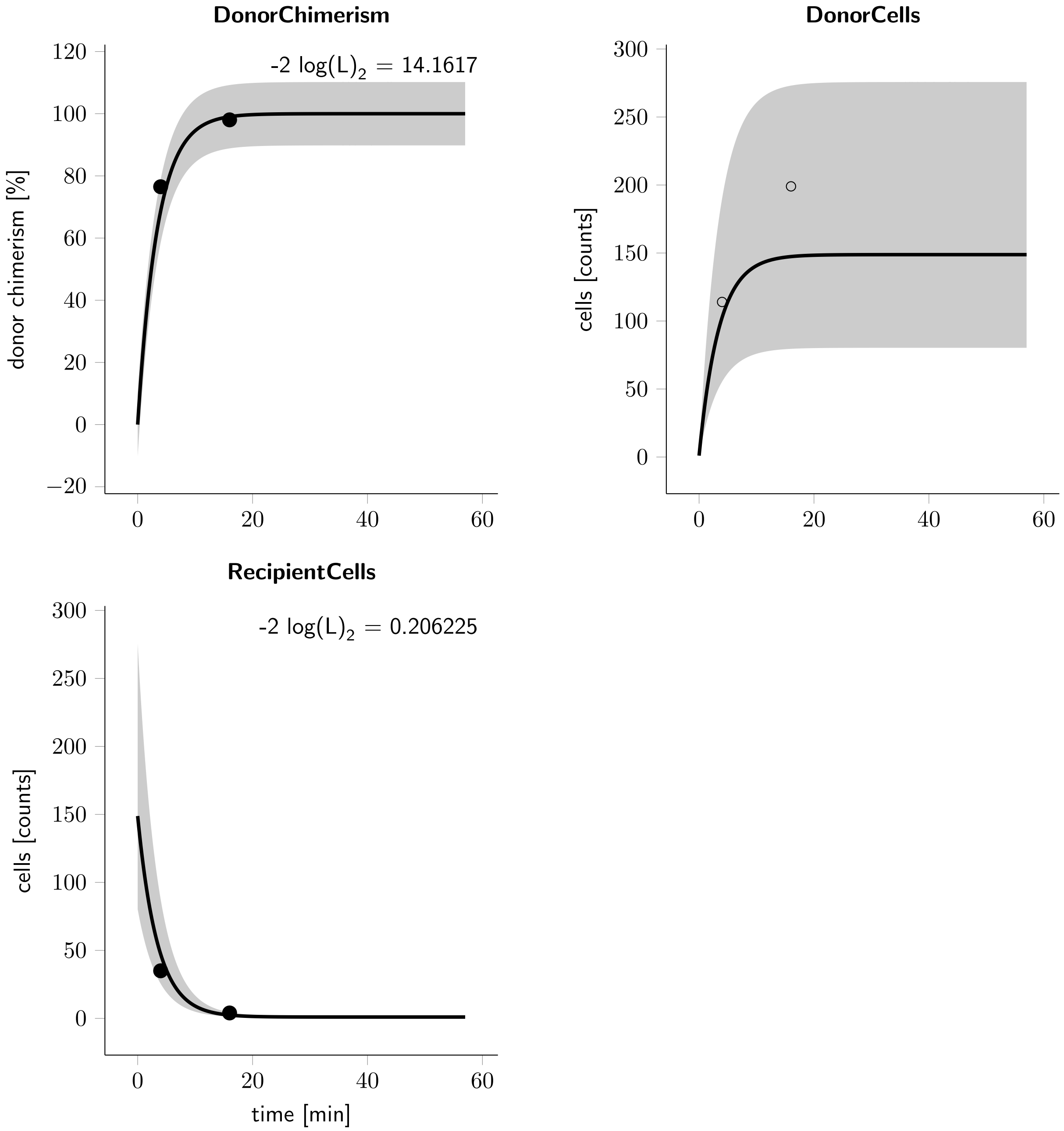


**Figure 9: Cluster_numbers_27Jan2023_ T4 observables and experimental data for the experiment.** The observables are displayed as solid lines. The error model that describes the measurement noise is indicated by shades.

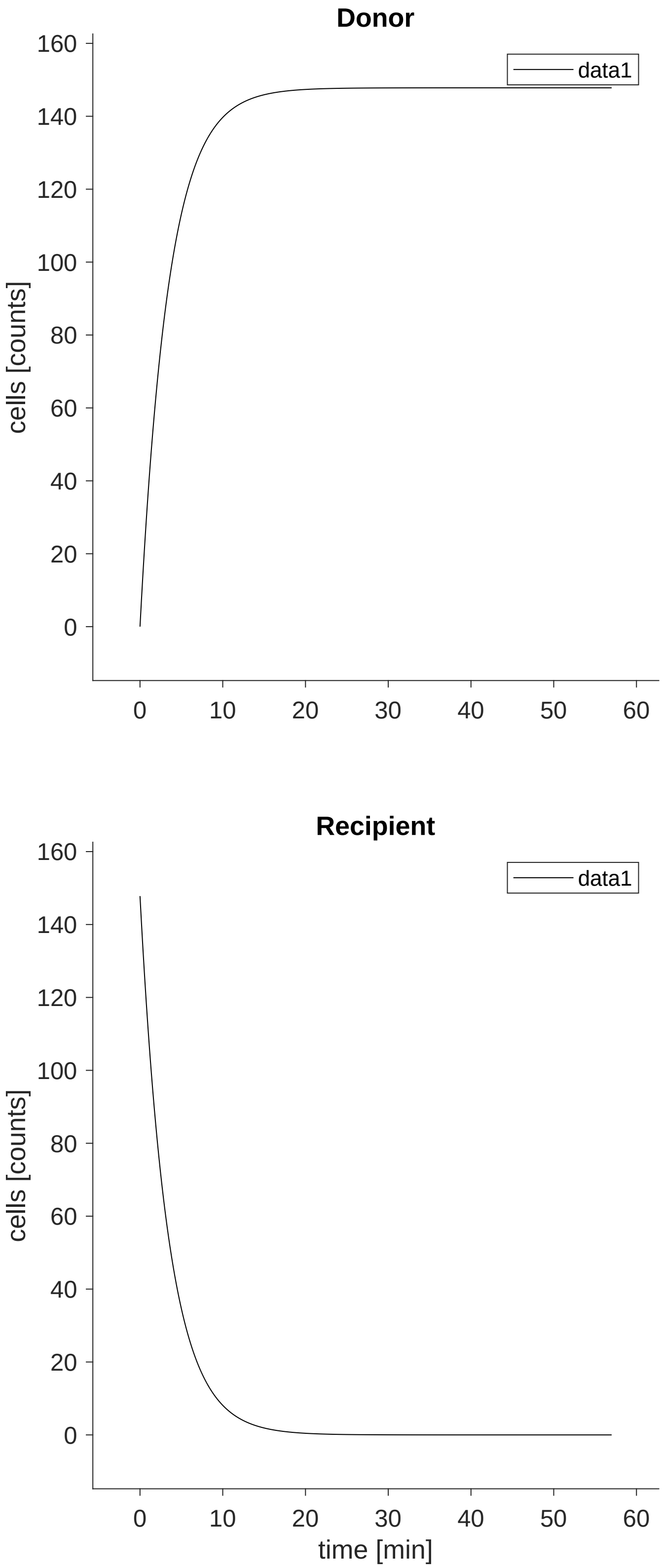




**Figure 10: Cluster_numbers_27Jan2023_ T4 trajectories of the input, dynamic and derived variables.** The dynamical behaviour is determined by the ODE system defined in Section
refHalfLifeModel$_ode$.

| time [min] | DonorChimerism donor chimerism [%] | DonorCells cells [counts] | RecipientCells cells [counts] |
|---|---|---|---|
| 4 | 11.8644 | 28 | 208 |
| 16 | 61.3861 | 62 | 39 |
| 52 | NaN | NaN | NaN |

**Table 6: Experimental data for the experiment Cluster_numbers_27Jan2023_ T5**

#### 1.12 Experiment: Cluster_numbers_27Jan2023_ T5

##### 1.12.1 Comments

data .def file template

##### 1.12.2 Experiment specific conditions

To evaluate the model for this experiment the following conditions are applied.

- **Local condition #6 (global condition #6):**

$$\mathrm{HalfLife\_ClusterID} \rightarrow \mathrm{HalfLife\_ClusterIDT5}$$

$$\mathrm{init\_Recipient} \rightarrow \frac{\mathrm{HalfLife\_ClusterIDT5} \cdot \mathrm{kin\_ClusterIDT5}}{\ln(2)}$$

$$\mathrm{kin\_ClusterID} \rightarrow \mathrm{kin\_ClusterIDT5}$$

##### 1.12.3 Experimental data and model fit

The model observables and the experimental data is shown in Figure 11. The agreement of the model observables and the experimental data, given in Table 6, yields a value of the objective function $\chi^2 = 8.07902$ for 4 data points in this data set. The trajectories of the input, dynamic and derived variables that correspond to the experimental conditions in this experiment are shown in Figure 12.

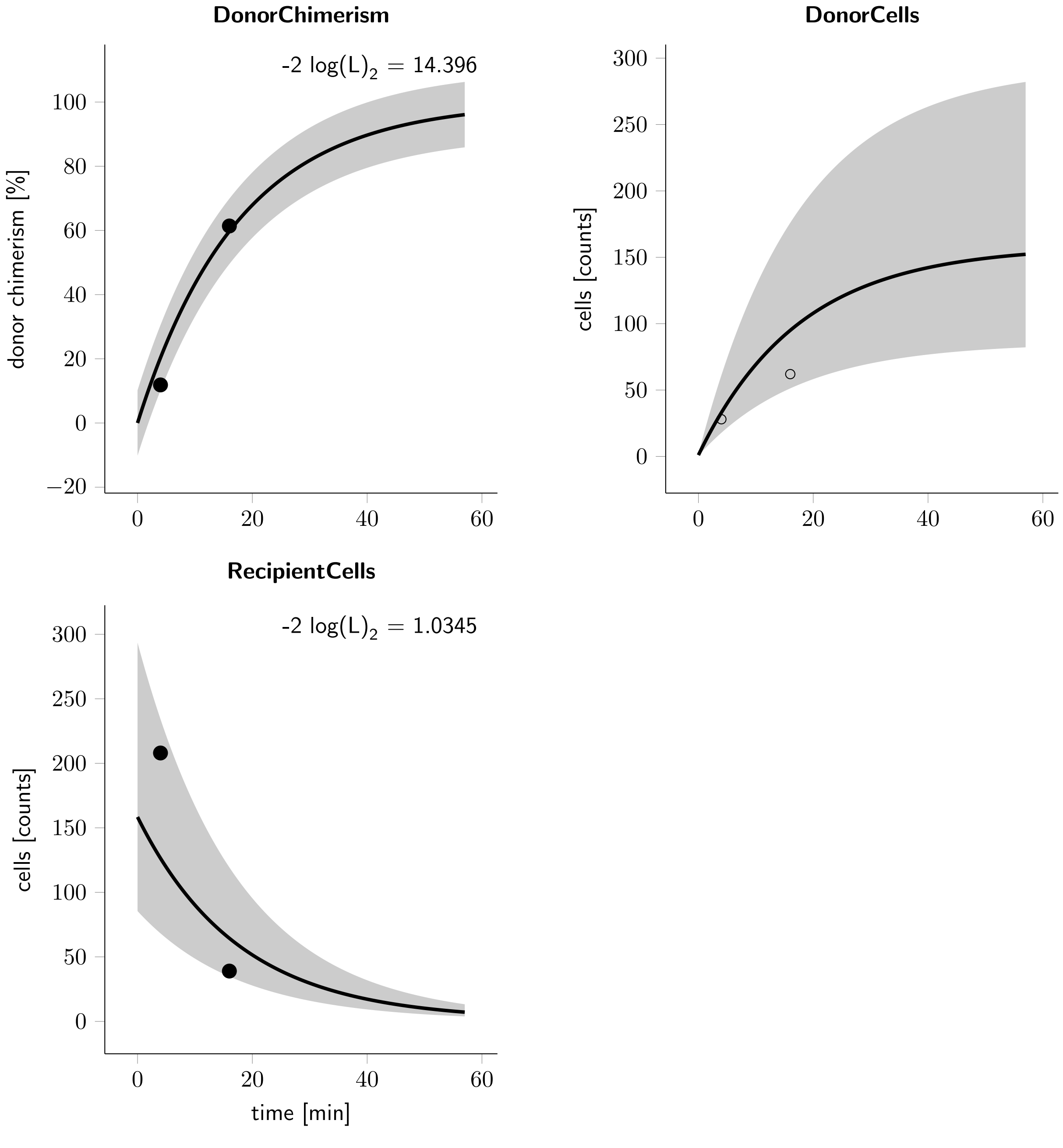


**Figure 11: Cluster_numbers_27Jan2023_ T5 observables and experimental data for the experiment.** The observables are displayed as solid lines. The error model that describes the measurement noise is indicated by shades.

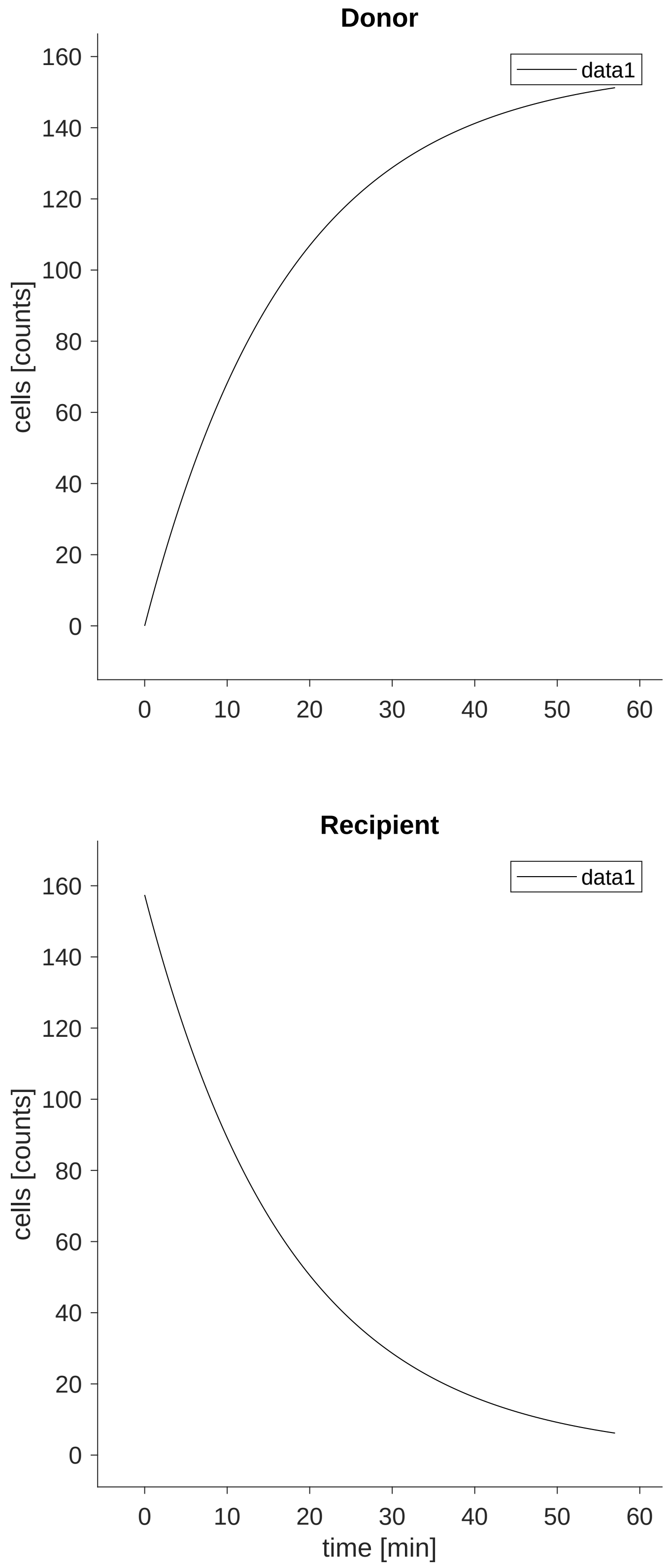


**Figure 12: Cluster_numbers_27Jan2023_ T5 trajectories of the input, dynamic and derived variables.** The dynamical behaviour is determined by the ODE system defined in Section
refHalfLifeModel$_{o}de$.

| time [min] | DonorChimerism donor chimerism [%] | DonorCells cells [counts] | RecipientCells cells [counts] |
|---|---|---|---|
| 4 | 10.2273 | 9 | 79 |
| 16 | 57.3171 | 47 | 35 |
| 52 | NaN | NaN | NaN |

**Table 7: Experimental data for the experiment Cluster_numbers_27Jan2023_ T6**

#### 1.13 Experiment: Cluster_numbers_27Jan2023_ T6

##### 1.13.1 Comments

data .def file template

##### 1.13.2 Experiment specific conditions

To evaluate the model for this experiment the following conditions are applied.

- **Local condition #7 (global condition #7):**

$$\mathrm{HalfLife\_ClusterID} \rightarrow \mathrm{HalfLife\_ClusterIDT6}$$
$$\mathrm{init\_Recipient} \rightarrow \frac{\mathrm{HalfLife\_ClusterIDT6} \cdot \mathrm{kin\_ClusterIDT6}}{\ln(2)}$$
$$\mathrm{kin\_ClusterID} \rightarrow \mathrm{kin\_ClusterIDT6}$$

##### 1.13.3 Experimental data and model fit

The model observables and the experimental data is shown in Figure 13. The agreement of the model observables and the experimental data, given in Table 7, yields a value of the objective function $\chi^2 = 5.43961$ for 4 data points in this data set. The trajectories of the input, dynamic and derived variables that correspond to the experimental conditions in this experiment are shown in Figure 14.

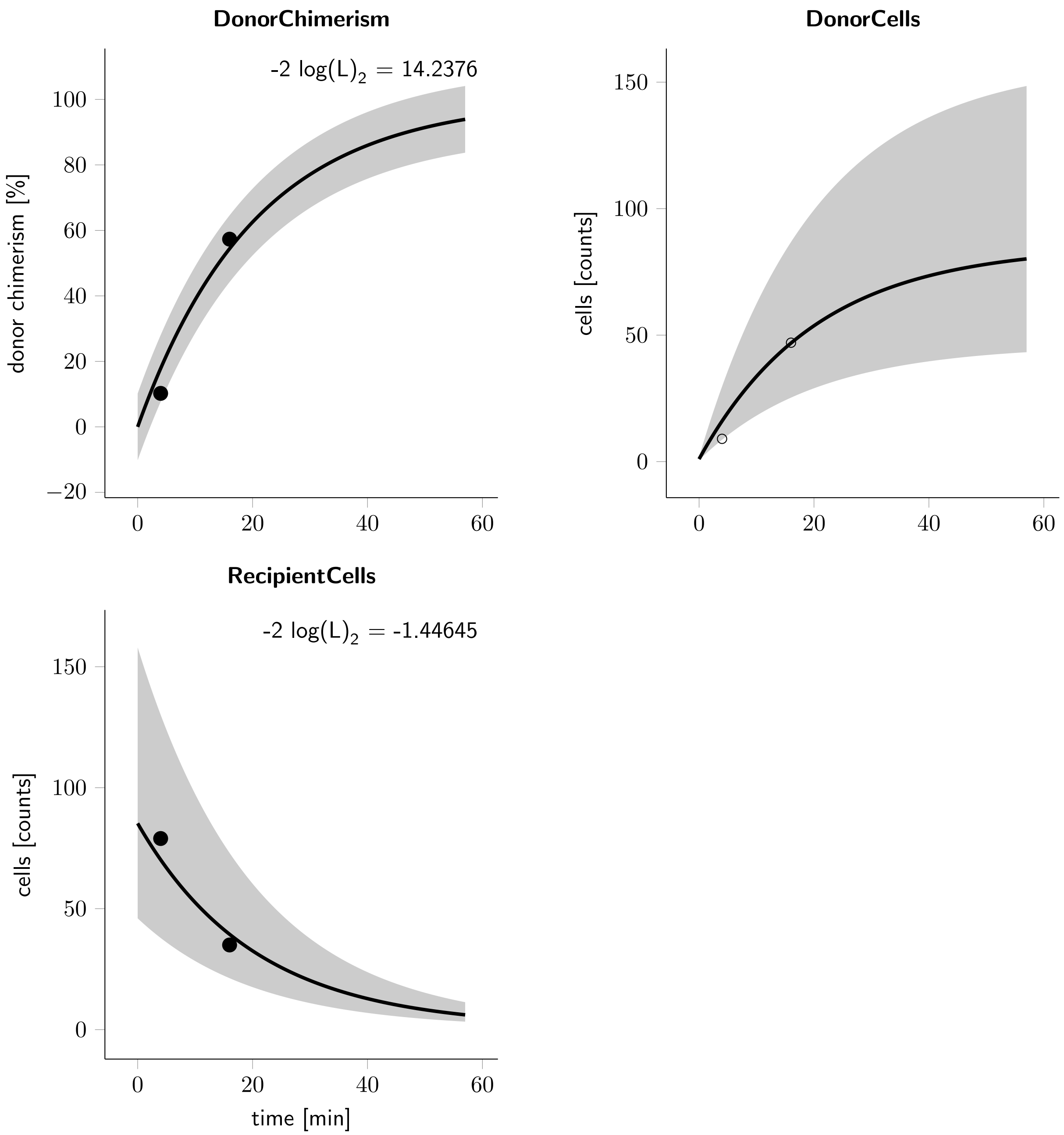


**Figure 13: Cluster_numbers_27Jan2023_ T6 observables and experimental data for the experiment.** The observables are displayed as solid lines. The error model that describes the measurement noise is indicated by shades.

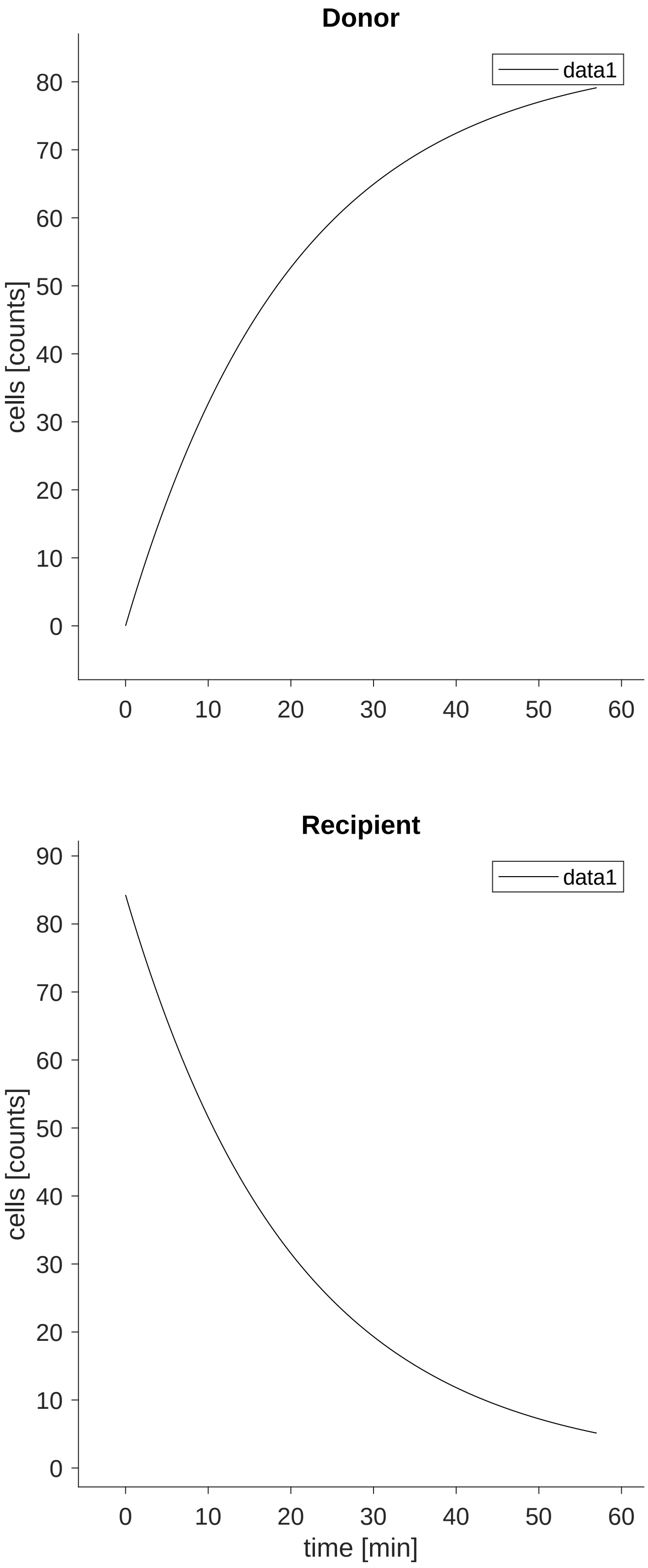




**Figure 14: Cluster_numbers_27Jan2023_ T6 trajectories of the input, dynamic and derived variables.** The dynamical behaviour is determined by the ODE system defined in Section
refHalfLifeModel$_{o}de$.

| time [min] | DonorChimerism<br>donor chimerism [%] | DonorCells<br>cells [counts] | RecipientCells<br>cells [counts] |
|---|---|---|---|
| 4 | 3.77358 | 2 | 51 |
| 16 | 45.1613 | 14 | 17 |
| 52 | NaN | NaN | NaN |

**Table 8: Experimental data for the experiment Cluster_numbers_27Jan2023_ T7**

#### 1.14 Experiment: Cluster_numbers_27Jan2023_ T7

##### 1.14.1 Comments

data .def file template

##### 1.14.2 Experiment specific conditions

To evaluate the model for this experiment the following conditions are applied.

- **Local condition #8 (global condition #8):**

$$\text{HalfLife\_ClusterID} \rightarrow \text{HalfLife\_ClusterIDT7}$$

$$\text{init\_Recipient} \rightarrow \frac{\text{HalfLife\_ClusterIDT7} \cdot \text{kin\_ClusterIDT7}}{\ln(2)}$$

$$\text{kin\_ClusterID} \rightarrow \text{kin\_ClusterIDT7}$$

##### 1.14.3 Experimental data and model fit

The model observables and the experimental data is shown in Figure 15. The agreement of the model observables and the experimental data, given in Table 8, yields a value of the objective function $\chi^2 = 7.04174$ for 4 data points in this data set. The trajectories of the input, dynamic and derived variables that correspond to the experimental conditions in this experiment are shown in Figure 16.

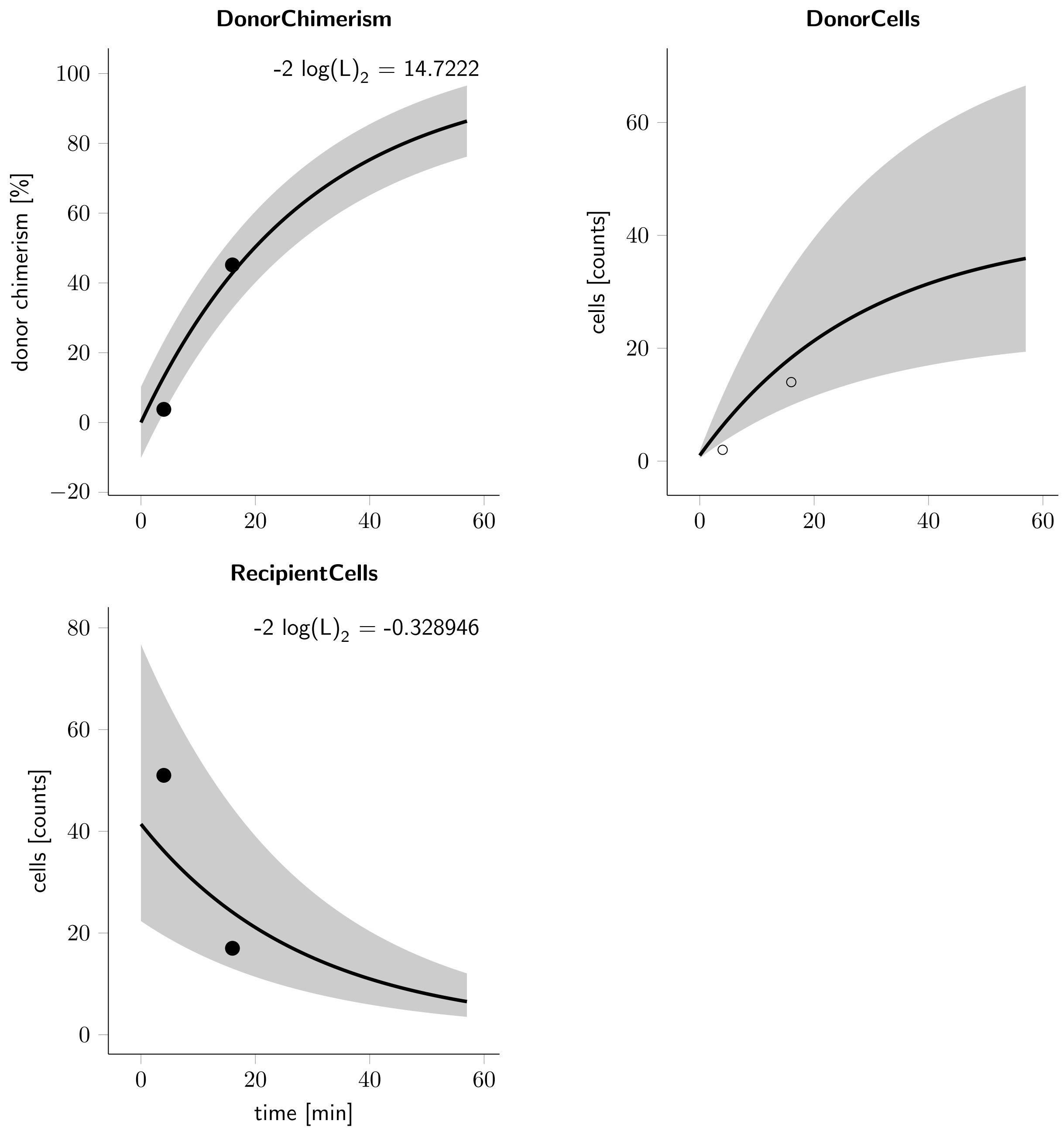


**Figure 15: Cluster_numbers_27Jan2023_ T7 observables and experimental data for the experiment.** The observables are displayed as solid lines. The error model that describes the measurement noise is indicated by shades.

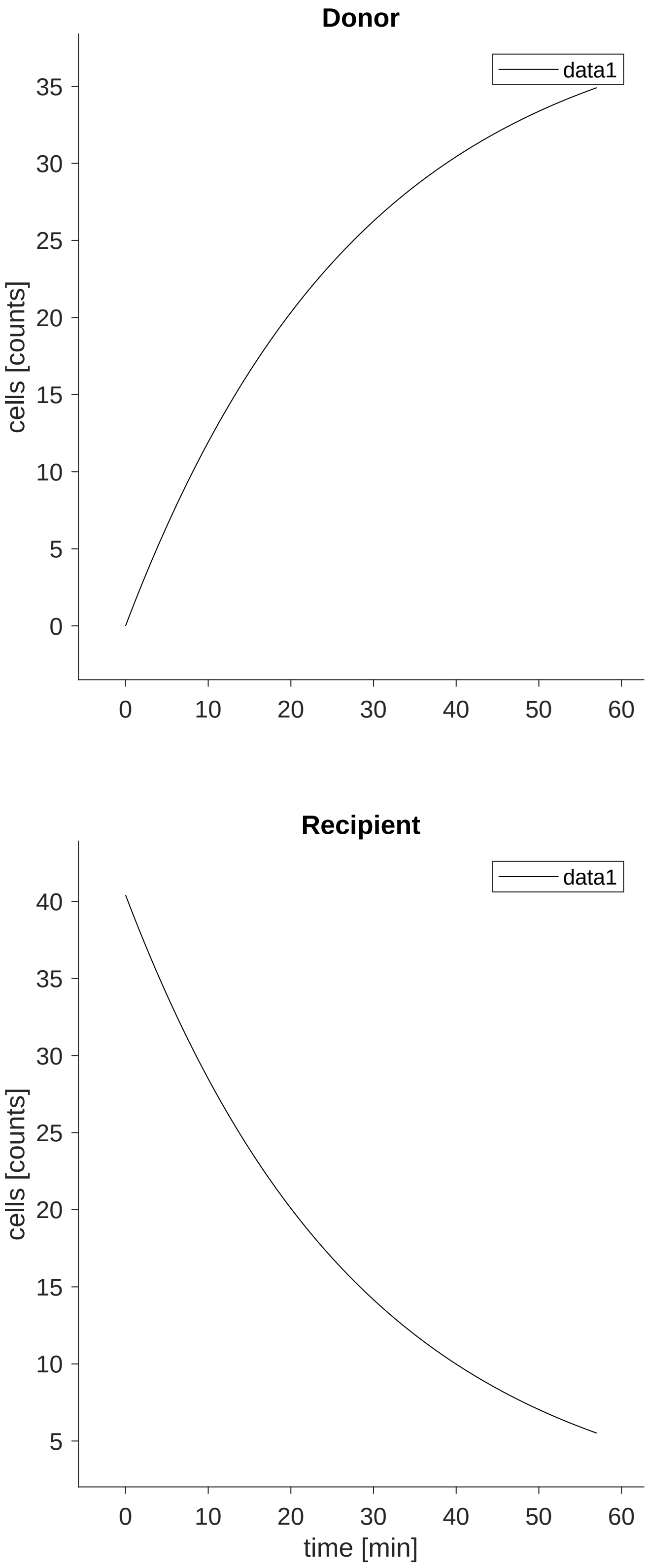




**Figure 16: Cluster_numbers_27Jan2023_ T7 trajectories of the input, dynamic and derived variables.** The dynamical behaviour is determined by the ODE system defined in Section
refHalfLifeModel$_ode$.

| time [min] | DonorChimerism donor chimerism [%] | DonorCells cells [counts] | RecipientCells cells [counts] |
|---|---|---|---|
| 4 | 23.3333 | 14 | 46 |
| 16 | 40.9091 | 9 | 13 |
| 52 | NaN | NaN | NaN |

**Table 9: Experimental data for the experiment Cluster_numbers_27Jan2023_ T8**

#### 1.15 Experiment: Cluster_numbers_27Jan2023_ T8

##### 1.15.1 Comments

data .def file template

##### 1.15.2 Experiment specific conditions

To evaluate the model for this experiment the following conditions are applied.

- **Local condition #9 (global condition #9):**

$$\text{HalfLife\_ClusterID} \rightarrow \text{HalfLife\_ClusterIDT8}$$

$$\text{init\_Recipient} \rightarrow \frac{\text{HalfLife\_ClusterIDT8} \cdot \text{kin\_ClusterIDT8}}{\ln(2)}$$

$$\text{kin\_ClusterID} \rightarrow \text{kin\_ClusterIDT8}$$

##### 1.15.3 Experimental data and model fit

The model observables and the experimental data is shown in Figure 17. The agreement of the model observables and the experimental data, given in Table 9, yields a value of the objective function $\chi^2 = 7.86557$ for 4 data points in this data set. The trajectories of the input, dynamic and derived variables that correspond to the experimental conditions in this experiment are shown in Figure 18.

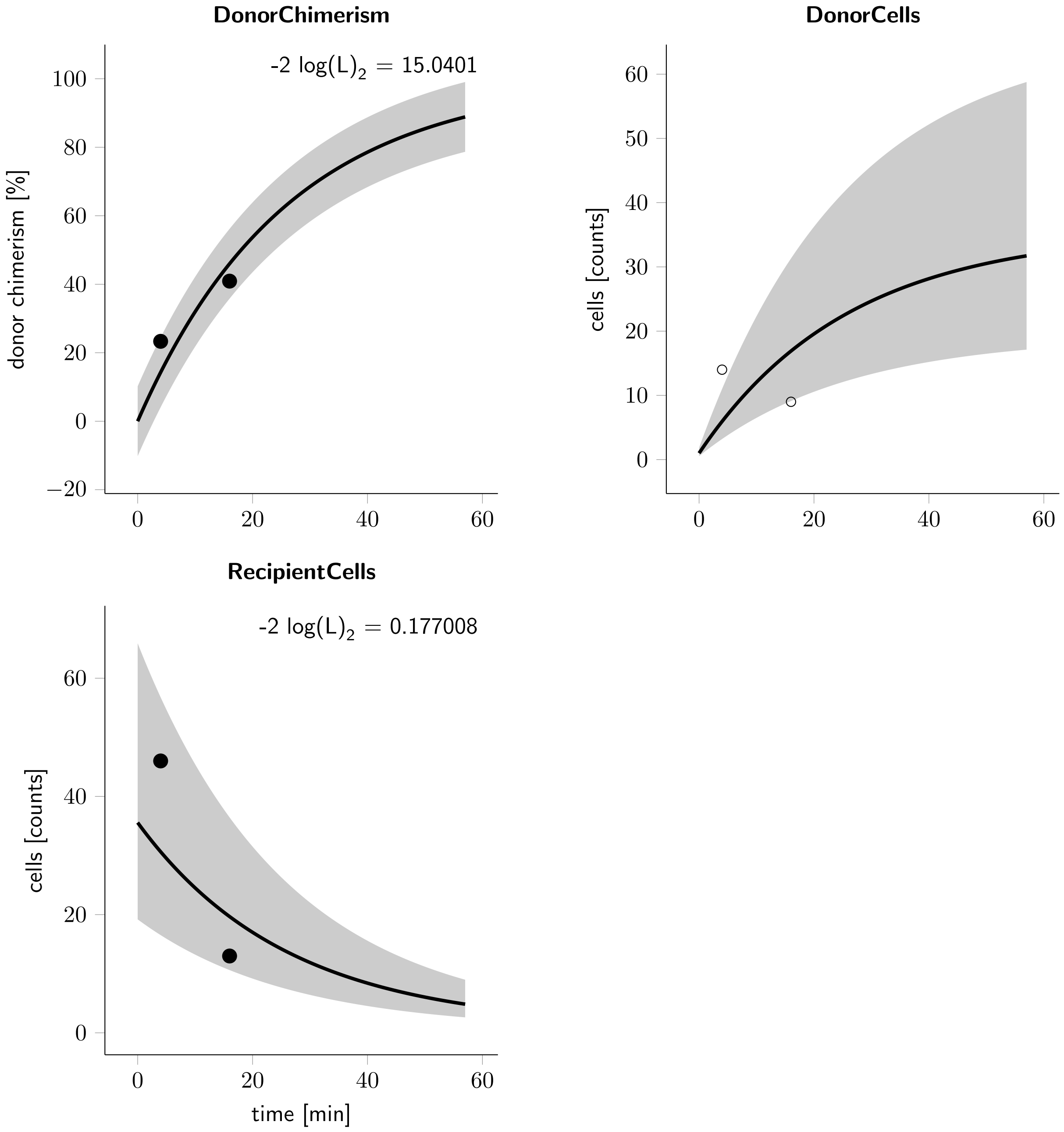


**Figure 17: Cluster_numbers_27Jan2023_ T8 observables and experimental data for the experiment.** The observables are displayed as solid lines. The error model that describes the measurement noise is indicated by shades.

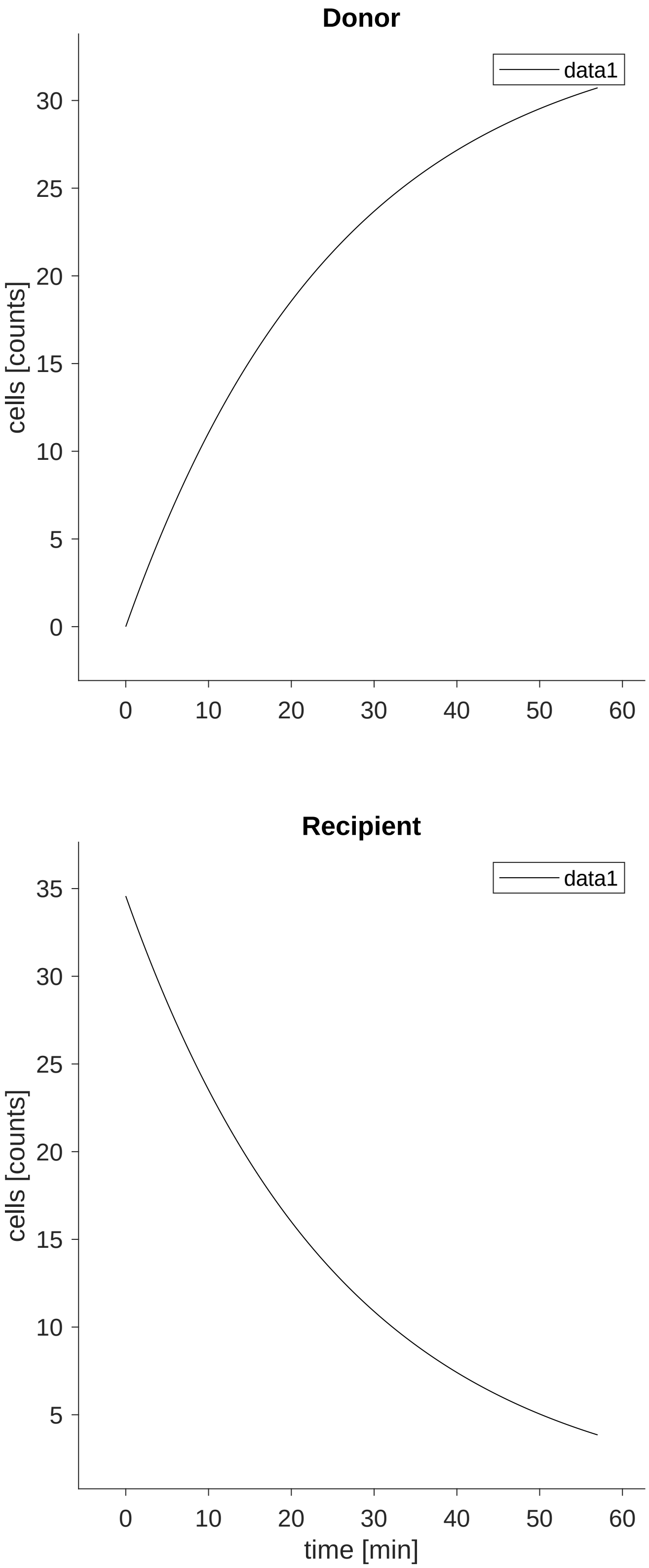




**Figure 18: Cluster_numbers_27Jan2023_ T8 trajectories of the input, dynamic and derived variables.** The dynamical behaviour is determined by the ODE system defined in Section
refHalfLifeModel$_ode$.

| | name | $\theta_{min}$ | $\hat{\theta}$ | $\theta_{max}$ | log | non-log $\hat{\theta}$ | estimated |
|---|---|---|---|---|---|---|---|
| 1 | HalfLife_ClusterIDT0 | -0.8 | +0.7854 | +3 | 1 | $+6.10 \cdot 10^{+00}$ | 1 |
| 2 | HalfLife_ClusterIDT1 | -0.8 | +1.1397 | +3 | 1 | $+1.38 \cdot 10^{+01}$ | 1 |
| 3 | HalfLife_ClusterIDT2 | -0.8 | +1.7504 | +3 | 1 | $+5.63 \cdot 10^{+01}$ | 1 |
| 4 | HalfLife_ClusterIDT3 | -0.8 | +1.2087 | +3 | 1 | $+1.62 \cdot 10^{+01}$ | 1 |
| 5 | HalfLife_ClusterIDT4 | -0.8 | +0.3781 | +3 | 1 | $+2.39 \cdot 10^{+00}$ | 1 |
| 6 | HalfLife_ClusterIDT5 | -0.8 | +1.0863 | +3 | 1 | $+1.22 \cdot 10^{+01}$ | 1 |
| 7 | HalfLife_ClusterIDT6 | -0.8 | +1.1498 | +3 | 1 | $+1.41 \cdot 10^{+01}$ | 1 |
| 8 | HalfLife_ClusterIDT7 | -0.8 | +1.2974 | +3 | 1 | $+1.98 \cdot 10^{+01}$ | 1 |
| 9 | HalfLife_ClusterIDT8 | -0.8 | +1.2553 | +3 | 1 | $+1.80 \cdot 10^{+01}$ | 1 |
| 10 | SD | -5 | -0.5721 | +3 | 1 | $+2.68 \cdot 10^{-01}$ | 1 |
| 11 | SD2 | -5 | +1.0083 | +3 | 1 | $+1.02 \cdot 10^{+01}$ | 1 |
| 12 | kin_ClusterIDT0 | -5 | +1.5254 | +3 | 1 | $+3.35 \cdot 10^{+01}$ | 1 |
| 13 | kin_ClusterIDT1 | -5 | +1.3003 | +3 | 1 | $+2.00 \cdot 10^{+01}$ | 1 |
| 14 | kin_ClusterIDT2 | -5 | +0.6592 | +3 | 1 | $+4.56 \cdot 10^{+00}$ | 1 |
| 15 | kin_ClusterIDT3 | -5 | +1.0796 | +3 | 1 | $+1.20 \cdot 10^{+01}$ | 1 |
| 16 | kin_ClusterIDT4 | -5 | +1.6324 | +3 | 1 | $+4.29 \cdot 10^{+01}$ | 1 |
| 17 | kin_ClusterIDT5 | -5 | +0.9515 | +3 | 1 | $+8.94 \cdot 10^{+00}$ | 1 |
| 18 | kin_ClusterIDT6 | -5 | +0.6167 | +3 | 1 | $+4.14 \cdot 10^{+00}$ | 1 |
| 19 | kin_ClusterIDT7 | -5 | +0.1500 | +3 | 1 | $+1.41 \cdot 10^{+00}$ | 1 |
| 20 | kin_ClusterIDT8 | -5 | +0.1243 | +3 | 1 | $+1.33 \cdot 10^{+00}$ | 1 |

**Table 10: Estimated parameter values**
$\hat{\theta}$ indicates the estimated value of the parameters. $\theta_{min}$ and $\theta_{max}$ indicate the upper and lower bounds for the parameters. The log-column indicates if the value of a parameter was log-transformed. If log $\equiv$ 1 the non-log-column indicates the non-logarithmic value of the estimate. The estimated-column indicates if the parameter value was estimated (1), was temporarily fixed (0) or if its value was fixed to a constant value (2).

### 2 Estimated model parameters

In total 20 parameters are estimated from the experimental data. The best fit yields a value of the objective function $-2\log(L) = 2587.93$ for a total of 37 data points. The model parameters were estimated by maximum likelihood estimation. In Table 10 the estimated parameter values are given. Parameters highlighted in red color indicate parameter values close to their bounds. The parameter name prefix init_ indicates the initial value of a dynamic variable.

### 3 Profile likelihood of model parameters

In order to evaluate the identifiability of the model parameters and to assess confidence intervals, the profile likelihood [**?** ] was calculated. An overview is displayed in Figure 19.

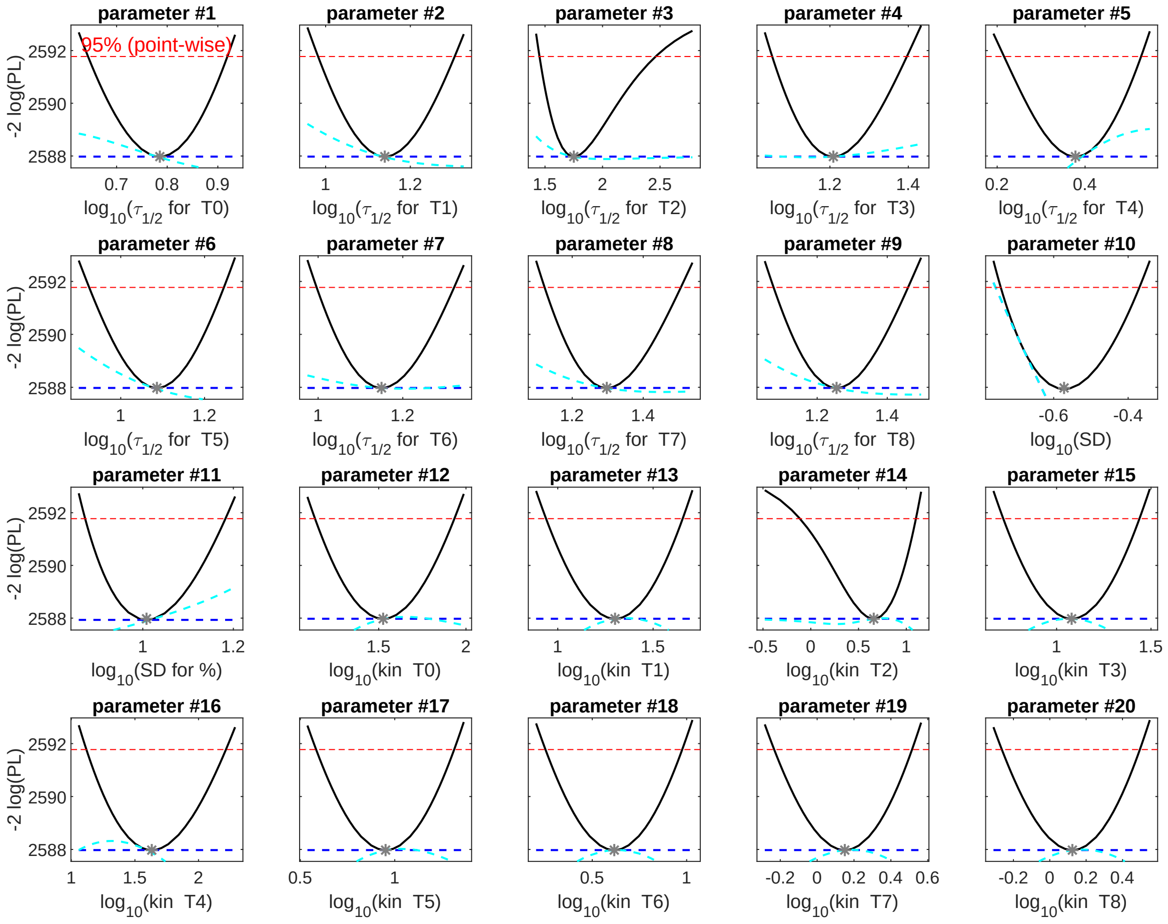




**Figure 19: Overview of the profile likelihood of the model parameters**
The solid lines indicate the profile likelihood. The broken lines indicate the threshold to assess confidence intervals. The asterisks indicate the optimal parameter values.

| | name | $\hat{\theta}$ | $\sigma^-$ | $\sigma^+$ |
|---|---|---|---|---|
| 1 | HalfLife_ClusterIDT0 | +0.785 | +0.643 | +0.920 |
| 2 | HalfLife_ClusterIDT1 | +1.140 | +0.981 | +1.306 |
| 3 | HalfLife_ClusterIDT2 | +1.750 | +1.451 | +2.472 |
| 4 | HalfLife_ClusterIDT3 | +1.209 | +1.052 | +1.398 |
| 5 | HalfLife_ClusterIDT4 | +0.378 | +0.214 | +0.528 |
| 6 | HalfLife_ClusterIDT5 | +1.086 | +0.924 | +1.247 |
| 7 | HalfLife_ClusterIDT6 | +1.150 | +0.996 | +1.322 |
| 8 | HalfLife_ClusterIDT7 | +1.297 | +1.121 | +1.507 |
| 9 | HalfLife_ClusterIDT8 | +1.255 | +1.075 | +1.461 |
| 10 | SD | -0.572 | -0.742 | -0.370 |
| 11 | SD2 | +1.008 | +0.873 | +1.183 |
| 12 | kin_ClusterIDT0 | +1.525 | +1.133 | +1.936 |
| 13 | kin_ClusterIDT1 | +1.300 | +0.936 | +1.657 |
| 14 | kin_ClusterIDT2 | +0.659 | -0.125 | +1.103 |
| 15 | kin_ClusterIDT3 | +1.080 | +0.715 | +1.441 |
| 16 | kin_ClusterIDT4 | +1.632 | +1.116 | +2.219 |
| 17 | kin_ClusterIDT5 | +0.952 | +0.581 | +1.317 |
| 18 | kin_ClusterIDT6 | +0.617 | +0.250 | +0.979 |
| 19 | kin_ClusterIDT7 | +0.150 | -0.235 | +0.517 |
| 20 | kin_ClusterIDT8 | +0.124 | -0.266 | +0.498 |

**Table 11: Confidence intervals for the estimated parameter values derived by the profile likelihood**
$\hat{\theta}$ indicates the estimated optimal parameter value. $\sigma^-$ and $\sigma^+$ indicate 95% point-wise confidence intervals.

### 4 Confidence intervals for the model parameters

In Table 11, 95% confidence intervals for the estimated parameter values derived by the profile likelihood [**?** ] are given.
